# A new foreperiod effect on single-trial phase coherence. Part I: existence and relevance

**DOI:** 10.1101/072371

**Authors:** Joaquin Rapela, Marissa Westerfield, Jeanne Townsend, Scott Makeig

## Abstract

Expecting events in time leads to more efficient behavior. A remarkable early finding in the study of temporal expectancy is the foreperiod effect on reaction times; i.e., the fact that the time period between a warning signal and an impendent stimuli, to which subjects are instructed to respond as quickly as possible, influences reaction times. Recently it has been shown that the phase of oscillatory activity preceding stimulus presentation is related to behavior. Here we connect both of these findings by reporting a novel foreperiod effect on the inter-trial phase coherence triggered by a stimulus to which subjects do not respond. Until now, inter-trial phase coherence has been used to describe a regularity in the phases of groups of trials. We propose a single-trial measure of inter-trial phase coherence and prove its soundness. Equipped with this measure, and using a multivariate decoding method, we demonstrate that the foreperiod duration modulates single-trial phase coherence. In principle, this modulation could be an artifact due to the decoding method used to detect it. We show that this is not the case, since the modulation can also be observed with a very simple averaging method. Although real, the single-trial modulation of inter-trial phase coherence by the foreperiod duration could just reflect a nuisance in our data. We argue against this possibility by showing that the strength of the modulation correlates with subjects’ behavioral measures, both error rates and mean-reaction times. We anticipate that the new foreperiod effect on inter-trial phase coherence, and the decoding method used here to detect it, will be important tools to understand cognition at the single-trial level. In Part II of this manuscript, we support this claim, by showing that attention modulates the strength of the new foreperiod effect in a trial-by-trial basis.

## 1 Introduction

The scientific study of temporal expectation begun with the birth of experimental psychology when Wundt (1874) and Woodrow (1914) discovered the effects on reaction times of the delay between a warning signal and an imperative stimulus to which subjects were instructed to respond as quickly as possible. This delay is called the foreperiod and this effect as the foreperiod effects on reaction times. Here we report a new foreperiod effect on the inter-trial phase coherence (ITPC, the degree of alignment in phases from multiple trials) of the electroencephalogram (EEG) evoked by standard stimuli, to which subjects did not to respond, following a warning signal. We call this effect the standard foreperiod (SFP) effect on ITPC.

When foreperiod durations are fixed across a block of trials, reaction times increase with increasing foreperiod duration (Wundt, 1874; Woodrow, 1914). This effect is commonly explained as arising from a reduced precision of temporal expectancy with increasing time, due to the deterioration of accuracy in time estimation for longer time intervals (Grondin, 2001). When foreperiod durations vary randomly from trial to trial, however, reaction times are commonly found to decrease with increasing foreperiod durations (Näätänen, 1970; Woodrow, 1914). This effect is often explained in terms of conditional probabilities, i.e., the probability that the imperative stimulus will occur in a next small time window given that it has not occurred yet. As time passes during a given foreperiod without the impending stimulus having been presented, the conditional probability of its occurrence increases in the reminder of the foreperiod. The cognitive system presumably learns this changing conditional probability early in the experimental session and exploits it to increase temporal expectation. A review on foreperiod effects appears in Niemi and Naatanen (1981).

Electrophysiological correlates of the foreperiod effect have mainly been investigated through the Contingent Negative Variation (CNV, Walter et al., 1964). The CNV is a slow negative shift in the base line of the EEG that develops between the warning signal and the imperative stimulus. Walter et al. (1964) showed that the CNV does not reflect sensory activity, but the contingency of the warning signal and the imperative stimulus; i.e., the probability that imperative stimulus follows the warning signal. If the imperative stimulus is suddenly omitted the CNV amplitude is reduced, and the CNV disappears after 20-50 trials without the imperative stimulus (Low et al., 1966). Its amplitude is significantly elevated when a motor response is required to the imperative stimulus, compared to when no motor response is needed (Walter et al., 1964). The CNV is not discernible in raw EEG records and requires trial averaging (6-12 trials in normal adults). The CNV is generated in the cortex, requiring inputs from the basal ganglia, and involving cerebral networks where the thalamus plays a critical role (Brunia and van Boxtel, 2001). The investigation of the cognitive functions that are reflected on the CNV has a long and rich history. Walter and colleagues [1964] first discussed the CNV as a sign of expectancy. Tecce (1972) postulated attention as a correlate of the CNV. Gaillard (1977, 1978) proposed that the late E wave of CNV may reflect preparation for the motor response. More recent experiments have linked the CNV with explicit (Macar and Vidal, 2003; Pfeuty et al., 2003, 2005) and implicit (Praamstra et al., 2006) time perception.

That the phase of brain oscillation plays an essential role for understanding human behavior is not a recent finding. Early EEG investigations reported that the instant when a subject performs a voluntary hand movement coincides with a specific phase of the alpha oscillation (Bates, 1951). However, in recent years we have witnessed exceptional new findings on the relation between phase and behavior. External visual (Regan, 1966) and auditory (Galambos et al., 1981) rhythmic stimuli entrain brain oscillations (i.e., drag brain oscillations to follow the rhythm of the stimuli), and this entrainment is modulated by attention in such a way that the occurrence of attended stimuli coincides with the phase of brain oscillations of maximal excitability (Lakatos et al., 2005, 2008, 2013; O’Connell et al., 2011; Besle et al., 2011; Gomez-Ramirez et al., 2011; Zion Golumbic et al., 2013; Cravo et al., 2013; Mathewson et al., 2009, 2011, 2012; Spaak et al., 2014; Gray et al., 2015). Alignment of the phase of brain oscillations to optimize performance has been observed in perception, motor control, and cognition. In perception, this alignment has been related to vision (Valera et al., 1981; Busch et al., 2009; Mathewson et al., 2009, 2012; Cravo et al., 2013; Hanslmayr et al., 2013; Spaak et al., 2014; Cravo et al., 2015; Gray et al., 2015; Milton and Pleydell-Pearce, 2016), audition (Rice and Hagstrom, 1989; Lakatos et al., 2008; Stefanics et al., 2010; Henry and Obleser, 2012; Ng et al., 2012; Neuling et al., 2012; Lakatos et al., 2013; Hickok et al., 2015), and speech (Ahissar et al., 2001; Luo and Poeppel, 2007; Howard and Poeppel, 2010; Cogan and Poeppel, 2011; Howard and Poeppel, 2012; Giraud and Poeppel, 2012; Morillon et al., 2012; Gross et al., 2013; Peelle et al., 2013; Luo et al., 2013; Xiang et al., 2013; Doelling et al., 2014; VanRullen et al., 2014; Park et al., 2015; Zoefel and VanRullen, 2015a, b, 2016). In addition, the alignment of phases of oscillations in the visual brain regions can be triggered crossmodally by auditory stimuli, and vice versa (Thorne et al., 2011; Kayser et al., 2008; Lakatos et al., 2009; Romei et al., 2012). In motor control, optimal alignment of brain oscillations has been found in eye movements (Bosman et al., 2009; Hamm et al., 2010; Saleh et al., 2010; Drewes and VanRullen, 2011), and cortico-spinal excitability (van Elswijk et al., 2010; Berger et al., 2014). It has also been ascribed to reaction speed (Lansing, 1957; Callaway and Yeager, 1960; Stefanics et al., 2010; Drewes and VanRullen, 2011; Thorne et al., 2011). In cognition, the alignment of brain oscillations has been linked to attention (Yamagishi et al., 2008; Lakatos et al., 2008, 2009, 2013; Sauseng et al., 2008; Capotosto et al., 2009; VanRullen et al., 2011; Gomez-Ramirez et al., 2011; Zumer et al., 2014; Liu et al., 2014; Mathewson et al., 2014; Gray et al., 2015; van Diepen et al., 2015), working memory (Bonnefond and Jensen, 2012; Myers et al., 2014), causality judgment (Cravo et al., 2015), stimuli coincidence (Milton and Pleydell-Pearce, 2016), temporal predictions (Samaha et al., 2015; Fujioka et al., 2012), and to the transmission of prior information to visual cortex (Sherman et al., 2016). Specially relevant for this manuscript are reports on effects of stimulus expectancy on phase alignment (Cravo et al., 2011, 2013; Stefanics et al., 2010; Zion Golumbic et al., 2013; Mathewson et al., 2012; Milton and Pleydell-Pearce, 2016). The alignment of phase in groups of trials at one brain region, as studied in this manuscript, has been related to the latency of saccadic eye movements (Drewes and VanRullen, 2011), visual contrast sensitivity (Cravo et al., 2013), visual perception (Busch et al., 2009; Mathewson et al., 2012; Spaak et al., 2014), visual attention (Gray et al., 2015), visual stimuli timing (Milton and Pleydell-Pearce, 2016), top-down visual attention (Yamagishi et al., 2008), speech perception (Zion Golumbic et al., 2013), cross-modal interactions (Thorne et al., 2011; Fiebelkorn et al., 2011), as well as to temporal expectation (Cravo et al., 2011; Stefanics et al., 2010). Also, the alignment of phase across brain regions (Hanslmayr et al., 2007), the coupling of phase across frequencies and brain regions (Fiebelkorn et al., 2013; Sauseng et al., 2008; Händel and Haarmeier, 2009), and the coupling between phase and amplitude (Voytek et al., 2010; Bonnefond and Jensen, 2015; Cravo et al., 2011) is predictive of some aspects of behavior. Reviews of the relation between prestimulus phase, perception, behavior, and cognition appear in Schroeder and Lakatos (2009); Mathewson et al. (2011); VanRullen et al. (2011); Thut et al. (2012); VanRullen et al. (2014). In sum, almost every aspect of behavior, perception, and cognition critically depends on the phase of brain oscillations.

Considering that both the foreperiod duration and the phase of EEG oscillations are related to anticipatory attention, we conjectured that the former could be modulating the latter. In addition, because influences of attention on EEG phase activity can be observed in single trials (e.g., Busch and VanRullen, 2010), we hypothesized that modulations of phase activity by the foreperiod duration could also be seen in single trials. In a previous investigation (Makeig et al., 2002), we showed that ITPC can be a mechanism responsible for the appearance of event-related potential features in trial-averaged measures of EEG activity. Continuing with this line of investigation, here we study effects of expectancy, as induced by the foreperiod duration, on the ITPC. For this, we developed a single-trial multivariate decoding method. We reasoned that if the SFP modulates the ITPC then we should be able to reliably decode the SFP duration (SFPD) from the ITPC evoked by a standard stimulus. In Section 2.2 we show that there exist a significant correlation between the SFPD and the ITPC evoked by a standard in a single trial. In principle, this correlation could be an artifact of the decoding method. In Section 2.3 we show that this is not the case, since the effect of the SFPD on ITPC can be observed by just comparing the average ITPC of trials closer and further away from the warning signal. It could be argued that the SFP effect on ITPC is a nuisance in EEG recordings. We argue against this possibility in Section 2.4, by showing significant correlations between the strength of the SFP effect and behavioral measures (detection errors and reaction speed). In Section 2.5 we study the timing of the SFP effect on ITPC. We conclude with a discussion in Section 3.

## 2 Results

We characterized the foreperiod effect on ITPC in the audio-visual oddball detection task schematized in Figure 1. Subjects had to detect visual or auditory deviants among visual or auditory standards. Attention-shifting cues (LOOK and HEAR) were interspersed among the standards and deviants. After a LOOK (HEAR) cue subjects had to detect visual (auditory) deviants. Thus, the LOOK (HEAR) cue directed attention to the visual (auditory) attended modality. These cues initiated a period of expectancy for the next deviant stimulus and and acted as a warning signal. Further details on the experiment appear in Section 5.1.

**Figure 1:**
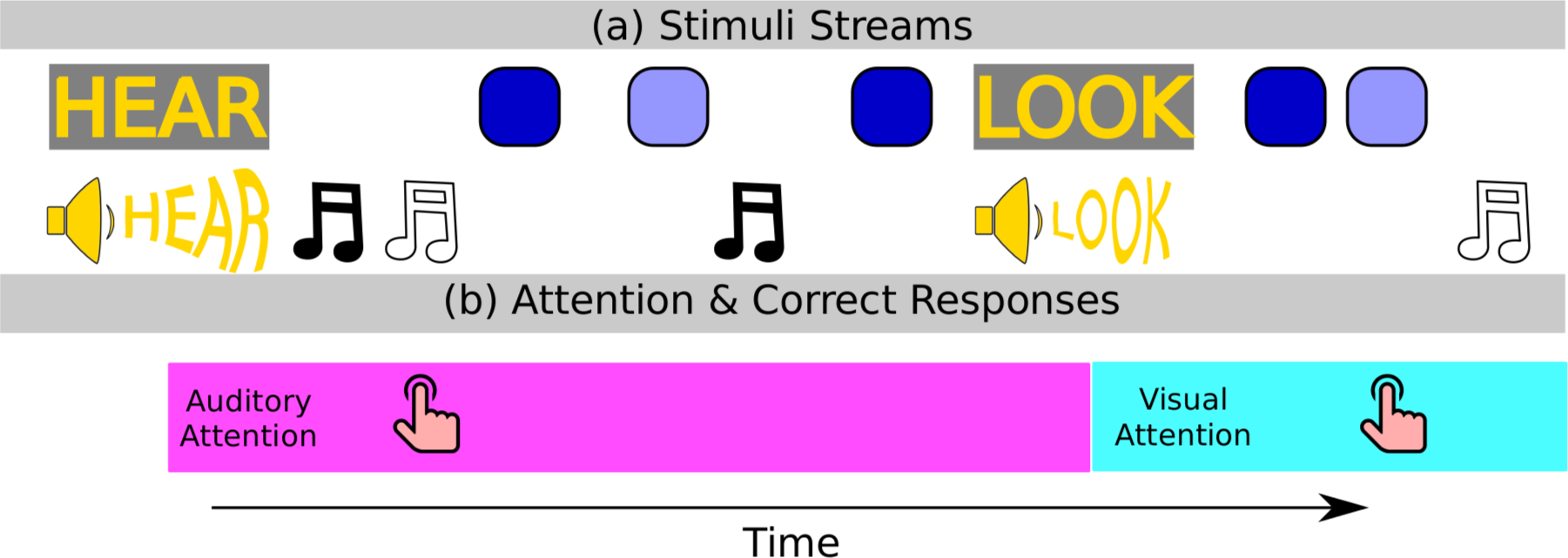
Audio-visual attention-shift experiment. (a) Audio-visual LOOK and HEAR attention-shifting cues, visual standards (dark-blue squares) and deviants (light-blue squares), and auditory standards (low-pitch sounds, filled notes) and deviants (high-pitch sounds, open notes) were presented sequentially in pseudo-random order. (b) After seeing and hearing a LOOK (HEAR) cue, subjects had to indicate with a button press when they saw a visual (heard an auditory) deviant.

All analysis were performed at the level of independent components, ICs, obtained from an Independent component analysis (ICA, Makeig et al., 1996) of the recorded EEG data. Figures 2a and 2b illustrate the main finding of this study. The thin curves are the cosine of the phase (at the peak ITC frequency, Section A.5.4; 8.15 Hz in this figure) of of the 20 epochs with the shortest (Figure 2a) and longest (Figure 2b) SFPD. These epochs were aligned at time zero to the presentation of attended visual standards from IC 13 of subject av124a. The thick red curve is the cosine of the average phase across all epochs (Section A.5.3). Between 150 and 250 ms the phase of the ten epochs with shortest SFPD (thin lines in Figure 2a) are more similar to the average phase (thick line in Figure 2a or in Figure 2b) than the phase of the ten epochs with longest SFPD (thin lines in Figure 2b). This suggests the existence of a foreperiod effect on ITPC where the SFPD modulates the similarity between the phase of a given epoch and the average phase across all epochs. The rest of this paper studies the validity of this effect and investigates its relation with subjects’ behavior (detection errors and reaction times).

**Figure 2:**
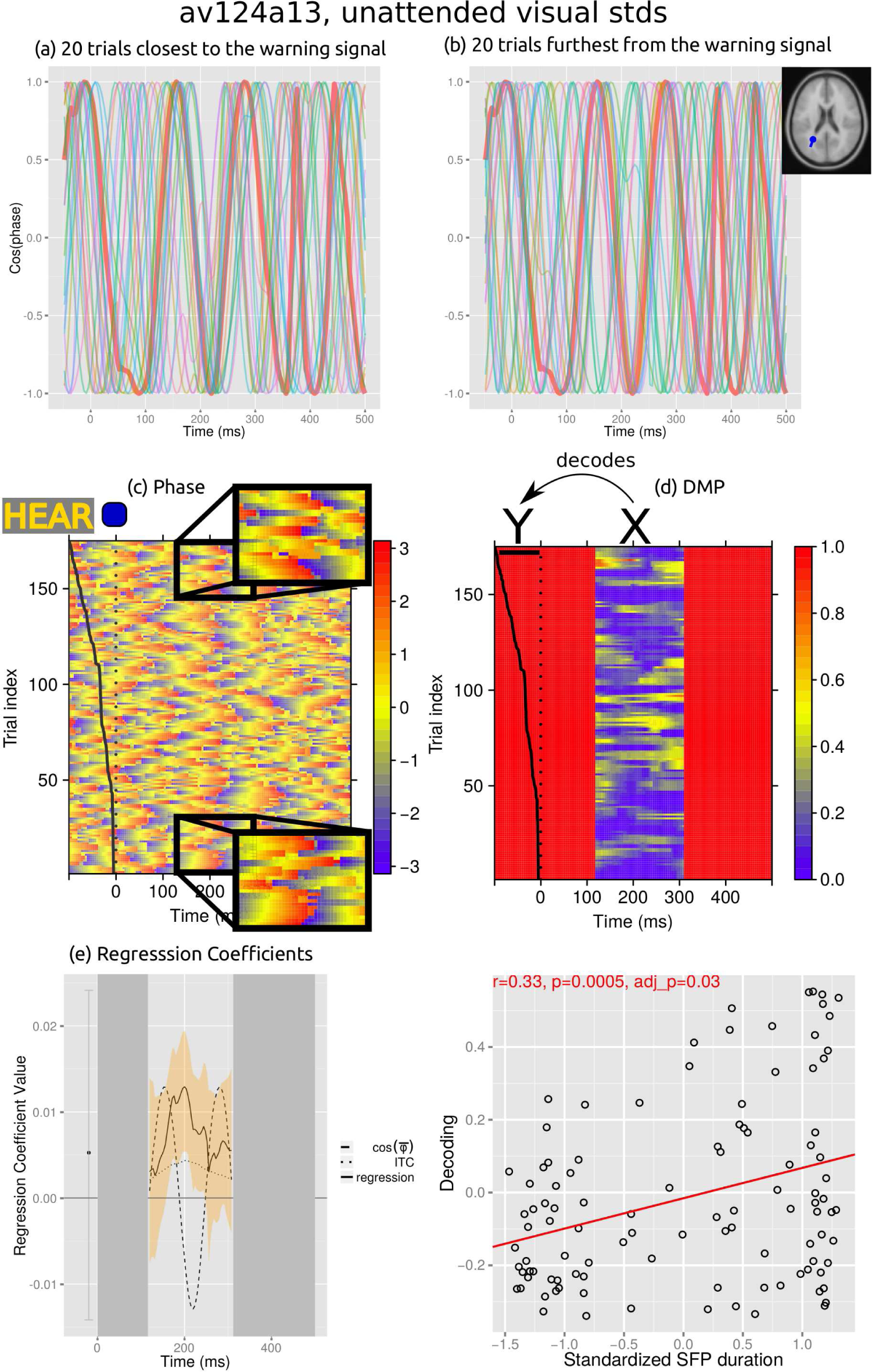
Data used for, and parameters obtained from, the analysis of trials from IC 13 of subject av124a, for epochs aligned to the presentation of unattended visual standards. (a, b) Cosine of the phase of the twenty epochs with the shortest (a) and the longest (b) SFPD. Thin curves are the cosines of the phases of individual epochs, and the thick curve is the cosine of the mean phase (i.e., mean direction, Eq. A.6) across all epochs. Between 150 and 250 ms, the phases of epochs with shortest SFPDs in (a) are more similar to the mean phase than the phases of epochs with longest SFPDs in (b). (c) Phase erpimage. Each horizontal line shows the phase of a single trial. For visualization clarity, the plot has been smoothed across trials using a running median with a window of size three. Time zero (dotted vertical line) corresponds to the presentation of standard visual stimuli. The solid black line indicates the (scaled) times of presentation of the warning signals immediately preceding the standards at time zero. Trials are sorted from bottom to top by increasing SFPD. Insets magnify phases closest (bottom) and furthest (top) from the warning signal. (d) DMP erpimage obtained from the phases in (b), smoothed with a running median across trials of size 5. Solid red regions correspond to time points where the distribution of phases across trials was not significantly different from the uniform distribution (p>0.01, Rayleigh test). (e) The solid line plots the coefficients of the linear regression model used to decode the SFPD (y in panel (b)) from the corresponding DMP values (x in panel (b)) of single trials. The orange band gives 95% bootstrap confidence intervals of the estimated coefficients (Section A.5.14). The dashed line at time t plots the mean phase at time t across all trials. The dotted line is the ITC curve (Section A.5.4). (f) Scatter plot of model decodings versus standardized SFPDs.

### 2.1 Behavioral results

We call deviant foreperiod duration to the delay between the presentation of a warning signal and the next deviant stimulus. We detected significant foreperiod effects on reaction times (i.e.; significant correlations between deviant foreperiod durations and subjects’ reaction times to the corresponding deviant stimuli) in ten combinations of subject and attended modality (26% out of a total of 38 combinations of 19 subjects and two attended modalities). Five of these combinations corresponded to the visual attended modality. These correlations were all positive for the visual attended modality (indicating that the longer the deviant foreperiod duration the longer the subject reaction time), while for the auditory attended modality there was a mixture of two positive and three negative significant correlations. Figure A.8a plots deviant foreperiod durations as a function of reaction times for an example subject and attended modality. We found significant foreperiod effects on detectability (i.e.; significant differences between the median deviant foreperiod duration for hits and misses) in other 10 combinations of subject and attended modality (26%). All of these combinations corresponded to the auditory attended modality and in all of them subjects detected more easily later deviants (i.e.; the median deviant foreperiod was significantly larger for hits than for misses). Figure A.8b plots deviant foreperiod durations for hits and misses for an example subject and attended modality.

Visual deviants were detected more reliably and faster than auditory ones. Mean error rates were 0.22 and 0.09 for detecting auditory and visual stimuli, respectively. A repeated-measures ANOVA, with error rate (in the detection of deviants of the attended modality) as dependent variable and attended modality as independent factor, showed a significant main effect of attended modality (F(1, 17)=39.35, p<0.0001). A posthoc analysis revealed that error rates were smaller when detecting visual than auditory deviants (z=6.45, p=5.59e-11). Mean response times to auditory and visual stimuli were 407 and 377 ms, respectively. A repeated-measures ANOVA, with mean response time (to deviant stimuli of the attended modality) as dependent variable and attended modality as independent factor, showed a significant main effect of attended modality (F(1, 18)=40.29, p<0.0001). A posthoc analysis revealed that mean response time was shorter for visual than auditory deviants (z=6.52, p=3.49e-11).

### 2.2 Measuring a new foreperiod effect on the ITPC elicited by stan-dards

Since we were interested in studying expectancy effects unrelated to motor preparation, we only characterized the foreperiod effect on ITPC for standards, to which subjects were instructed no to respond. Figure 2c shows phase values, at the peak ITC frequency (Section A.5.4), in single trials from IC 13 of subject av124a, and for epochs aligned to the presentation of unattended visual standards. Trials are sorted from bottom to top by increasing SFPD. The solid line represents the presentation time of the warning signal, scaled so that the maximum SFPD fits in the negative time period (the maximum, median, and minimum SFPD in this panel were 2,424 ms, 1,160 ms, and 328 ms, respectively). The top and bottom insets show magnified views of the phase of the trials with longest and shortest SFPDs, respectively. Panels (a) and (b) show the 20 bottommost and topmost trials from panel (c). Apparently, standards with shorter SFPD elicit stronger phase coherence from 150 to 250 ms after the presentation of the warning signal than standards with longer SFPD. Below we use decoding models to assess the validity of this observation.

To study if the SFPD modulates the ITPC evoked by standards, we attempted to decode the former from the latter. Reliable decodings would indicate that the SFPD is correlated with the ITPC evoked by standards. We used the Deviation from the Mean Phase (DMP) to quantify ITPC in single trials (Section 5.4). Figure 2d shows DMP values computed from the phase values in Figure 2c. Time points at which there was not enough statistical evidence to conclude that the phase distribution across different trials was statistically different from the uniform distribution (p>0.01, Rayleigh test) are masked in red. We did not use these points to decode the SFPD, for a reason explained in Section 5.4. For each epoch, we decoded the SFPD of the standard presented at time zero from a 500 ms-long window of DMP values following the presentation of this standard. To avoid possible EEG movement artifacts induced by responses to deviants, we excluded from the analysis trials with deviant stimuli in this 500-ms-long window. We used a linear regression model for decoding (Section A.5.7), and estimated its parameters using a variational-Bayes method (Section A.5.8). To each estimated model corresponds a standard modality (i.e., the modality of the standards used to align the epochs from which the model parameters were estimated; visual standard modality in Figure A.5.10) and an attended modality (i.e., the modality that was attended when the epochs used to estimate the model parameters were recorded; auditory attended modality in Figure A.5.10).

If the delay between the warning signal and the following deviant is longer than a threshold, that depends on the distribution of deviants, the foreperiod effect on reaction times disappears (Botwinick and Brinley, 1962). Similarly, if we estimate decoding models including a large proportion of standards presented long after the warning signal, decodings become non-significant (i.e., the SFP effect on ITPC disappears). To fit decoding models we used data from standards that were presented before a maximum SFP duration after the warning signal. The selection of this maximum is described in Section A.4.

The coefficients of the linear regression model estimated from the SFPDs and DMP values in Figure 2d are shown in Figure 2e. The leave-one-cross-validated decodings from the linear regression model are plotted as a function of SFPD in Figure 2f. To assess the accuracy of these decodings we calculated their robust correlation coefficient with the SFPDs (Section A.5.10). We obtained a correlation coeffi-cient r=0.33, which was significantly different from zero (p=0.0005, permutation test for skipped Pearson correlation coefficient; adjusted p-value for multiple comparisons adj p=0.03, Section A.5.9). This significance, and the fact that the estimated regression model was significantly different from the intercept-only model (p<0.01, likelihood-ratio permutation test, Section A.5.14), suggests that the SFPD is significantly correlated with the ITPC activity evoked by unattended visual standards in IC 13 of subject av124a.

To characterize the SFP effect on ITPC across subjects we grouped all ICs of all subjects using a clustering algorithm (Section A.5.2). Figure 3 shows the obtained clusters and Table A.3 provides additional information about them. To quantify the strength of the standard foreperiod effect on ITPC at a given cluster, standard modality, and attended modality, we computed the proportion of models with decodings significantly correlated with experimental SFPDs, as in Figure 2f. Dots below the image of a cluster in Figure 3 indicate that the decodings of more than 40% of the models, estimated from phase activity in that cluster and from the standard modality and attended modality given by the color of the dot, were significantly correlated with experimental SFPDs. For each cluster, standard modality, and attended modality, the number of models with significant correlations with the SFPDs, as well as the proportion of such models over the total number of models estimated for the cluster, standard modality, and attended modality, are given in Table A.4. The larger number of clusters with more than 40% significant correlations for the visual than the auditory modality (red vs. blue points) and for attended than unattended standards (darker vs. lighter colors) hints that the SFP effect on ITPC is stronger for the visual than the auditory modality, and for attended than unattended standards. For the visual standard modality, clusters where the decoding of more than 40% of models were significantly correlated with SFPDs are localized in parieto-occipital regions, while for the auditory standard modality these clusters are found on fronto-central brain regions, suggesting that the SFP effect on ITPC is somatotopically organized.

**Figure 3:**
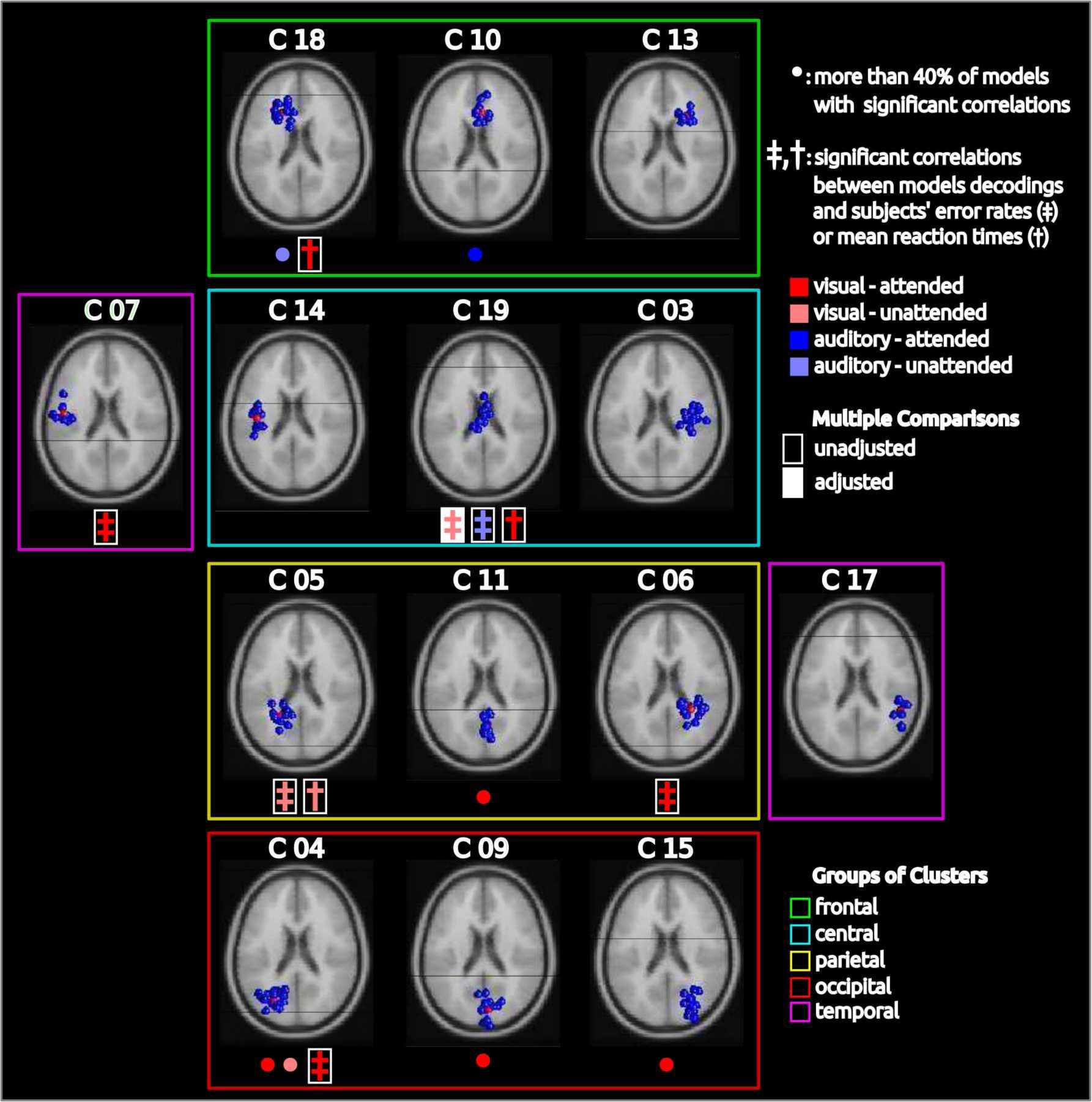
Clusters of ICs, large proportions of significant models, correlations with behavior, and groups of clusters. A blue ball inside a brain slice represents an IC from one subject. Each brain slice displays one cluster of ICs. A colored dot below the image of a cluster indicates that in more than 40% of the models estimated from data from that cluster, and from the standard modality and the attended modality given by the color of the dot, the correlation coefficient between models’ decoding and SFPDs was significantly different from zero (p adj<0.05, Section A.5.9). A dagger (double dagger) signals a significant correlation between models’ decoding power and subjects’ mean reaction times (error rates). Filled (unfilled) rectangles behind daggers indicate that the significance of the corresponding correlation test was corrected (uncorrected) for multiple comparisons. Colored boxes mark groups of clusters used in Section 2.5 to study the timing of the SFP effect on ITPC. The clusters with a large proportion of significant models suggest that the SFP effect on ITPC is stronger for visual than auditory standards and for attended than unattended standards, and that the effect is somatotopically organized (see text). The significant correlations between the strength of the SFP effect on ITPC and subjects’ behavioral performance show that the effect is behaviorally relevant.

From the successful decodings of SFPDs from ITPC activity triggered by standards, we infer that the SFPD is modulating the ITPC. However, other events, such at the warning signal, could be modulating this ITPC. We argue against this possibility in Section A.1.

### 2.3 Direct evidence for the SFP effect on ITPC

In the previous section we used linear regression models to decode SFPDs from ITPC activity following the presentation of standards. Since these decodings were reliable, we inferred that the SFPD is correlated with the ITPC activity triggered by standards. Here we use a simple method to gain more direct evidence of the existence of this correlation.

We compared the averaged phase decoherence of trials with the shortest and the longest SFPDs. The existence of a significant difference in these values would be direct evidence for the SFP effect on ITPC. To measure phase decoherence in a group of trials, we computed the mean DMP of the trials in the group. Section A.5.6 proves that, at times of high phase coherence, the mean DMP is related to the ITC, a standard measure of averaged ITPC. The red and blue lines in the top row of Figure 4 plot the mean DMP of the 20% trials furthest from and closest to the warning signal, respectively. Panels in the middle row plot the difference between the red and blue lines in the corresponding panels of the top row. Plots in different columns correspond to phase activity extracted from different subjects, ICs in the left parieto-occipital cluster 04, and attended visual stimuli. For subject av124a and IC 07 (left column), the mean DMP was significantly larger for trials with the longest SFPD between 0 and 100 ms after the presentation of standard stimuli, but smaller between 200 and 320 ms (Figures 4a and 4d). Differently, for subject av115a and IC 4 (central column), the mean phase decoherence was significantly larger for trials with the longest SFPD between 200 and 400 ms (Figures 4b and 4e). And for subject av113a and IC 17 (right column), the mean phase decoherence was significantly larger for trials with the longest SFPD between 0 and 250 ms (Figures 4c and 4f).

**Figure 4:**
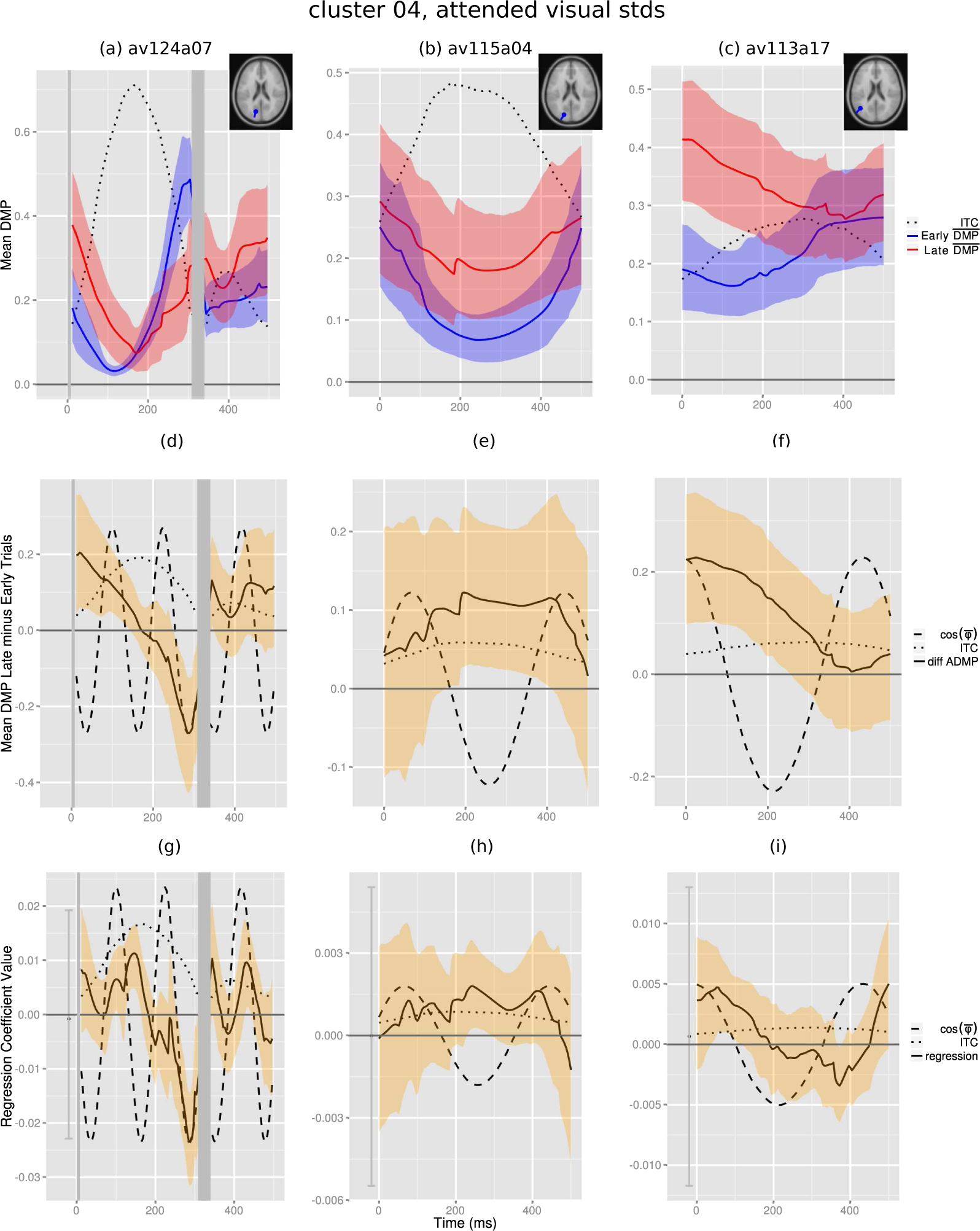
Relation between differences in phase coherence in trials furthest from and closest to the warning signal and regression models’ coefficients. Top row: averaged DMP evoked by the 20% standards closest to (blue curve) and furthest from (red curve) the warning signal. The dotted line plots the ITC computed from all trials. Middle row: the solid line plots the difference in averaged DMP evoked by trials furthest from minus closest to the warning signal (i.e., the red minus the blue curves in the top row). Bottom row: standardized coefficients of regression models. Colored bands in all panels represent 95% confidence intervals. The dotted and dashed lines in the panels of the middle and bottom rows plot the arbitrarily scaled ITC and the cosine of the mean phase from all trials, respectively. The top and middle rows demonstrate that ITPC depends on the SFPD, and the similarity between the middle and bottom rows show that the coefficients of a regression model indicate whether standards further from the warning signal evoke more coherence or decoherent activity than standards closer to it.

The linear regression method is statistically more powerful than the the simple method for testing the association between the SFPD and the ITPC activity triggered by standards, since it uses all trials in a statistically optimal way, and not just extreme trials in and ad-hoc manner. Another advantage of the linear regression method, compared with more general function approximation methods, such as artificial neural networks, is that the regression coefficients are readily interpretable. A positive regression coefficient indicates that trials with longer SFPD correspond to larger values of DMP, or larger decoherence, at the time of the regression coefficient, compared to trials with shorter SFPDs. This is because to decode a long SFPD a positive regression coefficient needs to be multiplied by a large DMP value (assuming that the reminding regression coefficients are kept fixed). Similarly, a negative regression coefficient implies that trials with longer SFPD correspond to smaller phase decoherence, at the time of the regression coefficient. To validate this interpretation, and the soundness of the decoding methodology, the bottom row in Figure 4 plots the regression coefficients of the decoding models corresponding to the top rows. Figure 4g shows significantly positive regression coefficients between 100 and 200 ms and significantly negative coefficients between 240 and 300 ms. Therefore, according to the previous interpretation, trials with longer SFPDs should correspond to more decoherent activity between 100-200 ms and to less decoherent activity between 240 and 300 ms. And this is what we observed in Figures 4a and 4d. Similar consistencies hold for the other columns. The normalized crosscorrelation (Section A.5.14) between the difference in averaged DMP (middle row in Figure 4) and the corresponding regression coefficients (bottom row of Figure 4) was 0.80 for (d)-(g), 0.39 for (e)-(h), and 0.38 for (f)(i). Across all models significantly different from the intercept-only model, the first, second (median), and third quartiles of the normalized crosscorrelation distribution were 0.40, 0.71 and 0.87, respectively. These results indicate that on average regression coefficients were similar to differences in average DMP, validate the previous interpretation of regression coefficients, and support the inference that reliable decodings by the regression models indicate modulations of ITPC by the SFPD.

### 2.4 The SFP effect on ITPC is correlated with subjects’ behavior

In the previous sections we presented evidence suggesting that the SFPD influences the ITPC evoked by standards. Here we show that this is a relevant effect, since its strength is correlated with subjects’ error rates and mean reaction times.

The ordinate in Figure 5a gives the correlation coefficient between decodings from a model and experimental SFPDs, for all subjects with an IC in the central-midline cluster 19 and unattended visual standards. The abscissa provides subjects’ error rates. The significant negative correlation between models’ decoding accuracies and subjects’ error rates (r=-0.91, p=0.0004, adj p=0.05) shows that the higher is the association between SFPD and ITPC in a subject, the better is his or her behavioral performance (error rate). Figure 5b is as Figure 5a, but for attended visual standards and the left parieto-occipital cluster 04. We also found significant correlations between the decoding accuracy of models and subjects mean reaction times, as illustrated in Figures 5c and 5d. Points marked in green in these figures represent outliers detected in the calculation of robust correlation coefficients (Section A.5.10). Note that the decoding models were optimized to decode SFPDs. These significant correlations resulted without fitting the decoding models to behavioral data.

**Figure 5:**
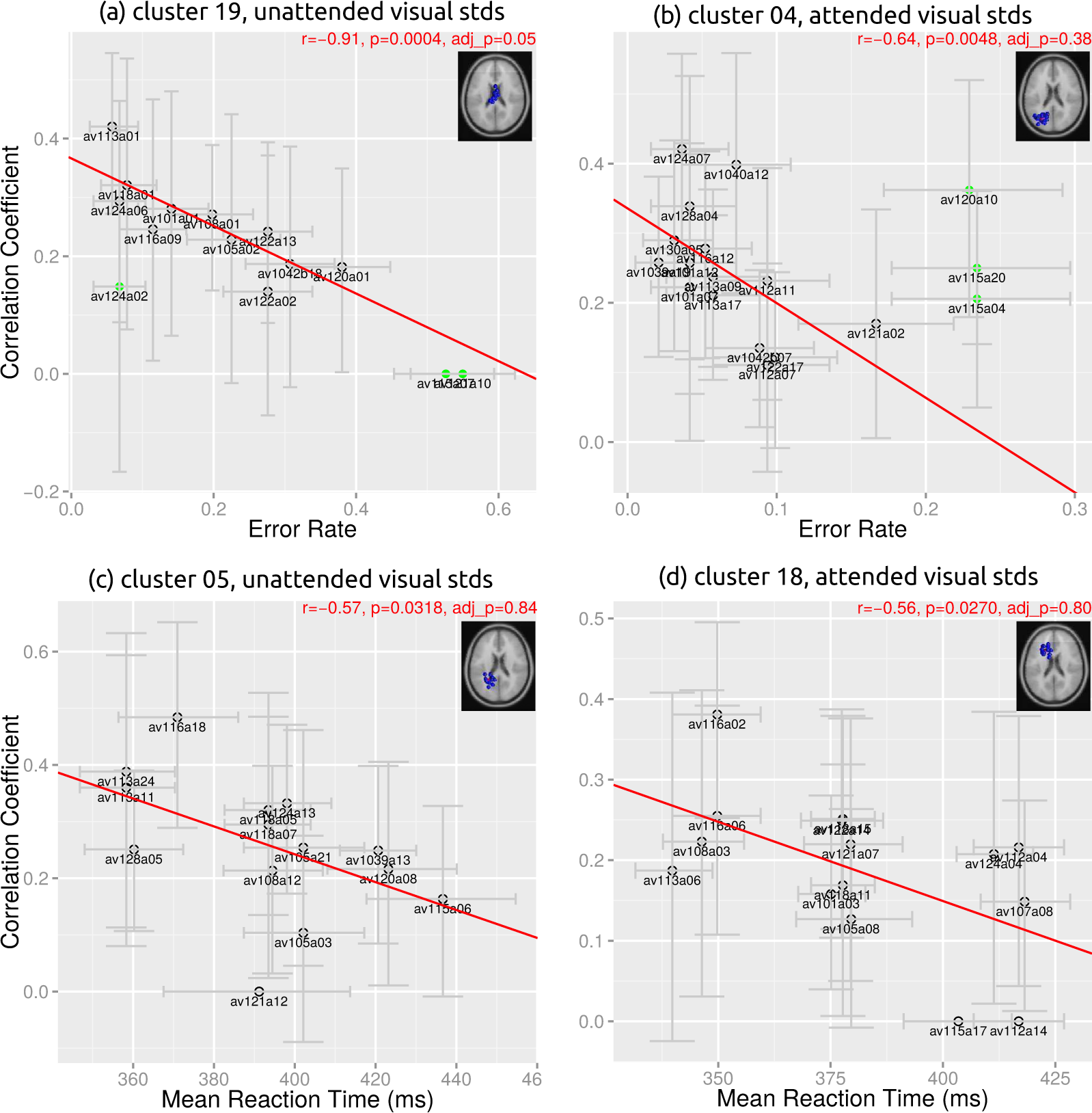
The strength of SFP effect on ITPC (i.e., the accuracy of the decoding models) is related to subjects’ behavior (error rates in (a, b) and mean reaction times in (c, d)), attesting the behavioral relevance of the SFP effect on ITPC. Larger decoding accuracy (i.e., larger correlation coefficients between models decodings and SFPDs; ordinate in all panels) corresponds to lower error rates (abscissa in (a, b)), as indicated by negative correlation coefficients, r, in (a, b), and this correspondence is stronger in models estimated from phase activity triggered by unattended standards (e.g., |r| is larger in (b) than in (a)). Larger decoding accuracy seems to correspond to faster (slower) mean reaction times (abscissa in (c, d)) in models estimated from phase activity triggered by attended (unattended) standards (e.g., r in in (c) and (d) is negative and positive, respectively), but this correspondence is not as conclusive as for error rates (see text).

A colored double and simple dagger below the image of a cluster in Figure 3 indicates a significant correlation between the decoding accuracy of models estimated from phase activity in the cluster and subjects error rates and mean reaction times, respectively. The color of the dagger corresponds to the standard modality and the attended modality of the data used to estimate the models’ parameters. Table A.5 shows the correlation coefficients, and corresponding p-values, between the decoding accuracy of models and subjects error rates. Table A.6 is as Table A.5, but for mean reaction times. All significant correlations between model’s decoding accuracies and subject’s error rates were negative (Table A.5), indicating that the larger the detection performance of a subject, the larger the strength of the SFP effect on ITPC. The absolute value of these correlations was larger for attended than unattended standards (p<1e-04; permutation test). For mean reaction times, the three significant correlations were also negative (Table A.6), showing that subjects with strongest SFP effect on ITPC reacted the slowest. However, differently from error rates, positive correlation for mean reaction times almost reach significance (e.g., cluster 06 and unattended auditory standards, or cluster 13 and unattended visual standards, Table A.6). Correlations (both significant and non-significant) were stronger for error rates than for mean reaction times and stronger for the visual than the auditory standard modality. An ANOVA with the absolute value of the correlation coefficient as dependent variable showed significant main effects of behavioral measure type (i.e., error rate or mean reaction time; F(1, 109)=7.26, p=0.0082) and for standard modality (F(1, 109)=8.22, p=0.005). A posthoc analysis indicated that the mean absolute value of the correlation coefficient was larger for error rates than for mean reaction times (p=0.005; Tukey test) and larger for the visual than the auditory standard modality (p=0.0082; Tukey test). Further information on this ANOVA appears in Section A.5.12.

The above correlations with error rates, although significantly different from zero at the 0.05 confidence level for univariate hypothesis tests, were found after computing a total of 56 correlations (14 clusters x 2 standard modalities x 2 attended modalities). Due to the multiple-comparison problem (Westfall and Young, 1993) the probability that at least one of these correlations was found significant while there was no association between subjects’ error rates and models’ decoding accuracies is very large (p ≃ 0.94). Only the correlation for the central cluster 19 and unattended visual standards (Figure 5a) remained significant after adjusting for multiple comparisons using the resampling procedure described in Section A.5.9. It is thus highly probable that some of the five unadjusted significant correlations with error rates that did not survive the multiple comparison test occurred by chance. However, note that, as expected, all of these correlations were negative, which is a very rare event under the assumption that all of these correlations occurred by chance (i.e., the probability of finding five negative correlations assuming equal probability for positive and negative correlations is p = 0.5^5^ = 0.03). Thus, not all of the five unadjusted significant correlations that did not survive the multiple comparison adjustment may have occurred by chance. None of the correlations with mean reaction times survived the multiple comparisons adjustment. Nonetheless, since the brain regions were these unadjusted significant correlations occurred have been previously connected to foreperiod effect on reaction times, as we discuss below, it is possible that not all these correlations occurred by chance.

### 2.5 Timing of the SFP effect on ITPC

In Section 2.3 we advanced an interpretation of the coefficients of the decoding model, and used a simple trial averaging procedure to verify that this interpretation was sound. As shown in Figures 2e and 4d-f, modulations of ITPC by the SFPD, as represented by the coefficients of decoding models, are not constant in time, but fluctuate in an oscillatory manner. In the 500 ms window following the presentation of a standard, these coefficients displayed one or more peaks, as in Figures 2e and 4g-h, respectively. Details on the identification of these peaks are given in Section A.5.13. The time of the largest peak corresponds to the latency after the presentation of standards where modulations of ITPC by the SFPD are strongest. In this section we use this time to represent the timing of the SFP effect on ITPC, and study how this timing varies across brain regions, standard modalities, and attended modalities.

The median time of the largest peak of the decoding model coefficients was 292 ms, with a 95% confidence interval of [284, 300] ms. We grouped the clusters of ICs into five groups, as indicated in Figure 3, and examined the mean time of the largest peak for each group of clusters and attended modality (Figure 6a), and for each group of clusters and and standard modality (Figure 6b). We found that peaks occurred earlier when attention was oriented to the auditory than to the visual modality (Figure 6a). This modulation by attention was strongest at the central group of clusters, where the peak of the models’ coefficients occurred earlier for auditory than visual standards (Figure 6b). Therefore, the timing of the SFP effect on ITPC was most strongly modulated, by the attended modality and by the standard modality, at central brain regions. In addition, for visual standards, the SFP effect on ITPC occurred earlier at occipital than central brain regions (Figure 6b).

**Figure 6:**
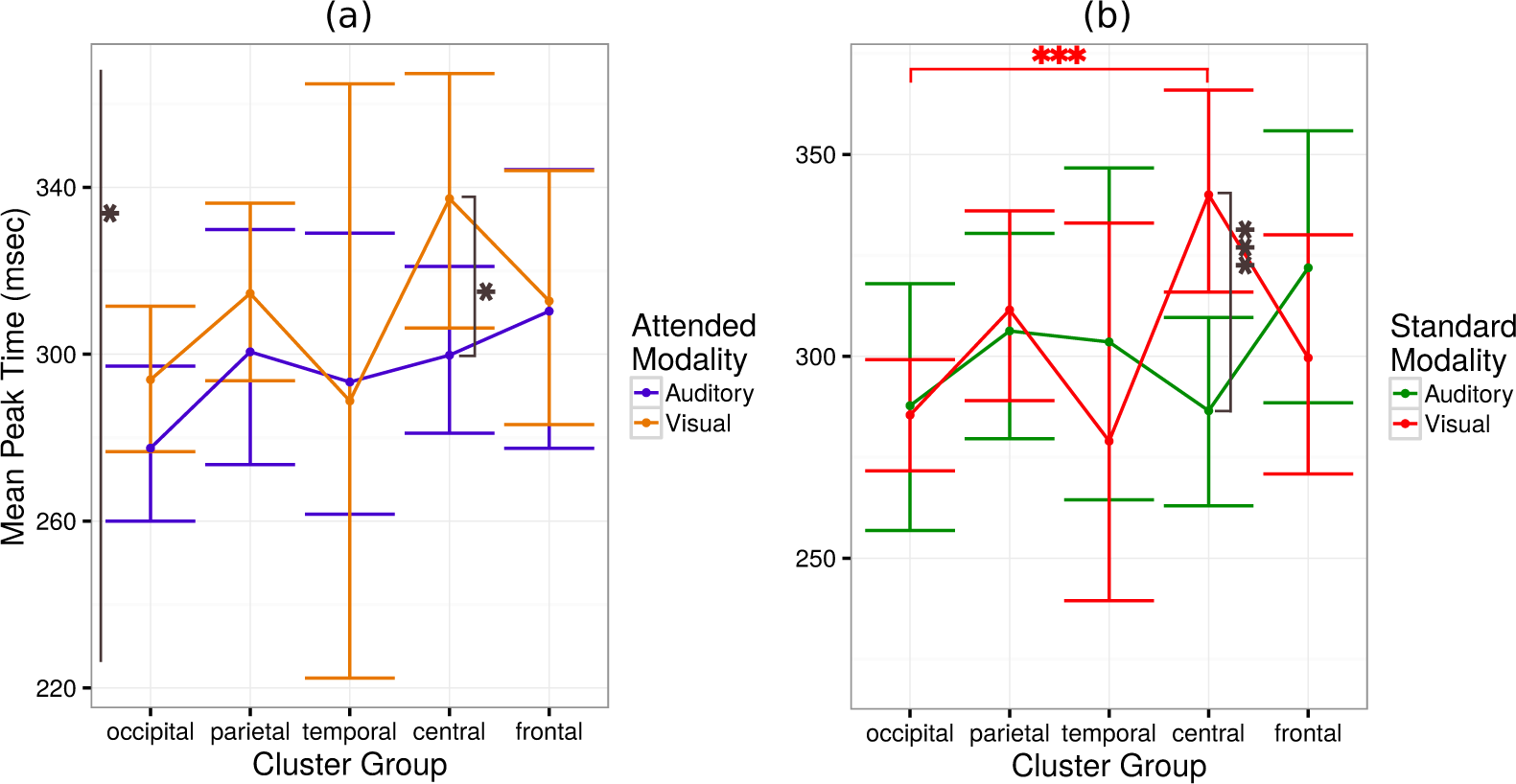
Timing of the SFP effect on ITPC. (a) Mean coefficients peak times for models corresponding to the auditory (blue points) and visual (orange points) attended modality, as a function of group of clusters. On average, when attending to the audition, the SFP effect on ITPC occurred earlier than when attending to vision. (b) Mean coefficients peak times for models corresponding to the auditory (green points) and visual (red points) standard modality, as a function of group of clusters. On average, for visual standards, the SFP effect on ITPC occurred earlier in the occipital than in central brain regions. The effect was most strongly modulated over the central brain region by both the attended modality (panel (a)) and by the standard modality (panel (b)).

A first ANOVA, using data from all models significantly different from the intercept only model (p<0.01; likelihood-ratio permutation test, Section A.5.14), with time of the largest peak of the coefficients as dependent variable, found a significant main effect of attended modality (F(1,230)=4.259, p=0.0402). A posthoc test showed that the peak occurred earlier for the auditory than the visual attended modalities (z=1842, p=0.0327; black asterisk to the left of Figure 6a). A second ANOVA restricted to models corresponding to the visual standard modality found a significant main effects of group of clusters (F(4,135)=4.4073). A posthoc analysis revealed that the peak was earlier for the occipital than the central group of clusters (z=4.163, p=3.14e-05; red asterisks in Figure 6b). A third ANOVA restricted to models corresponding to the central group of clusters found significant main effects of attended modality (F(1,43)=4.5061, p=0.0396) and of standard modality (F(1,43)=14.2173, p=0.0005). A posthoc analysis showed that the peak was earlier when attention was oriented to the auditory than visual modality (z=2.178, p=0.029360; black asterisk next to Central in Figure 6a) and earlier for auditory than visual standards (z=3.406, p=0.000659; black asterisks in Figure 6b).

## 3 Discussion

Here we reported a new foreperiod effect on the ITPC activity triggered by standards. We demonstrated that, when visual or auditory standards are preceded by a warning signal (Figure 1), ITPC is modulated by the delay between the warning signal and the standards (Figures 2, clusters with a colored dot in Figure 3, and blue entries in Table A.4). We used a decoding method to detect the new foreperiod effect. We demonstrated that this effect is not an artifact of the decoding method, since it can be observed using simple trial averages (Figure 4). Importantly, the strength of the new foreperiod effect (i.e.; the accuracy in decoding the SFPD from the phase activity elicited by a standard) is correlated with subjects’ detection performance and reaction speed (example and summary of correlations with detection performance in Figures 5a and 5b, clusters with a double dagger in Figure 3, and blue entries in Table A.5; with reaction speed in Figures 5c and 5d, clusters with a dagger in Figure 3, and blue entries in Table A.6). The effect occurred earlier when attention was oriented to the auditory modality (Figure 6a), for visual standards it happened earlier at occipital than in central brain regions (Figure 6b), and modulations of the timing of the effect by the attended modality or by the standard modality was stronger over a central brain regions (Figures 6a and 6b).

Phase coherence has been used to describe the synchrony between different recording electrodes (e.g., Lachaux et al., 1999; Hanslmayr et al., 2007) or, as in this study, to characterize the alignment of phases across multiple trials at a single site (e.g., Busch et al., 2009; Thorne et al., 2011; Cravo et al., 2015). Previous measures of phase alignment at a single site were the result of averaging data across multiple trials. However, phase coherence may vary from trial to trial, and important information could be lost in trial averages. We have previously proposed the ERP image plot, a useful visualization of trial-by-trial consistencies in a set of event-related data trials (Makeig et al., 2004; Delorme et al., 2015). To build this plot, trials are first sorted in order of a relevant data or external variable and then plotted as a color-coded two-dimensional image, as in Figure 2c. ERP image plots have proven useful in revealing trial-to-trial consistencies otherwise hidden in trial averages (Makeig et al., 1999; Jung et al., 2001; Delorme et al., 2007; Onton and Makeig, 2009). However, methods are lacking to establish the statistical significance of apparent features in these plots. Here we used a multivariate linear regression model to assess the significance of the observation, from the ERP image in Figure 2c, that the ITPC evoked by standards was modulated by the SFPD. We thus found a new foreperiod effect on the single-trial ITPC triggered by standards. Importantly, this effect would have been lost if we had indiscriminately averaged phases across all trials.

The fact that models accurately decoded the SFP duration from the ITPC triggered by standard stimuli does not necessarily mean that these two quantities are correlated with each other. In principle, a sufficiently complex model can decode arbitrarily accurately any experimental variable from any physiological measurements, if the complex model overfitted the data (Geman et al., 1992). To avoid overfitting we used leave-one-out crossvalidation to generate model decodings. To further validate that the obtained results were not an artifact of our decoding method, we showed that similar modulations of ITPC were obtained by a simple trial averaging procedure (Section 2.3). In addition, we developed surrogate controls to further support the hypothesis that (1) the SFP duration is the experimental variable causing the SFP effect on ITPC and (2) the ITPC triggered by standards is the aspect of brain activity modulated by the SFP effect on ITPC (Section A.1). Further evidence for the reliability of our results comes from the significant correlations between the decoding power of the models and subjects behavioral measures (Section 2.4) and from the anatomical specificity of the clusters with a large proportion of significant models (Section 2.2).

### 3.1 What psychophysical processes generate the SFP effect on ITPC?

The current investigation suggests that temporal expectancy plays an important role in the generation of the SFP effect on ITPC, although further investigations are needed to validate this suggestion, as discussed below. First, the strongly periodic stimulation in our experiment should have induced temporal expectations in subjects. This claim is supported by the observation of reliable foreperiod effects on reaction times in 26% of the subjects and attended modalities (Section 2.1) and by previous arguments relating the foreperiod effect on reaction times to expectancy (e.g., Niemi and Naatanen, 1981). Second, temporal expectations contribute to faster reaction times and improved perception (e.g., Correa, 2010), while the strength of the SFP effect on ITPC was larger in subjects achieving larger detection rates and faster mean reaction times (Figure 5 and Tables A.6 and A.5). Third, although it is possible to develop temporal expectancies for unattended stimuli, these expectancies are stronger for attended stimuli. Hence, the strength of the foreperiod effect on ITPC should be larger for attended than unattended stimuli (part II of this manuscript). Fourth, as we discuss in the next section, most of the brain regions where we observed significant correlations between the strength of the SFP effect on ITPC and behavioral measures, have been implicated in temporal expectation.

An unresolved question in the field of temporal expectation is whether it affects motor (i.e., rate of motor response), premotor (i.e., response preparation), perceptual (i.e., buildup of information about the stimulus), or executive (i.e., decision mechanism) stages. The majority of the evidence suggests a motor influence (e.g., Sanders, 1998; Brunia and Boelhouwer, 1988; Coull and Nobre, 1998), pre-motor effects have also been reported (e.g., Bausenhart et al., 2006; Hackley and Valle-Inclan, 1999; Müller-Gethmann et al., 2003), and more recent studies have shown influences on perceptual (e.g., Correa et al., 2005; Mento et al., 2013; Lange, 2009; Rolke, 2008) and executive (e.g., Naccache et al., 2002; Correa et al., 2010) functions. Results from the current study suggest that the aspect of temporal expectation measured by the SFP effect on ITPC mostly affects high-order visual processing. That the strength of the SFP effect on ITPC is significantly correlated with stimuli detectability (Figure 5a, and 5b and Table A.5) and that the SFP effect on ITPC is calculated from EEG activity triggered by standard stimuli to which subjects did not respond, indicates that the temporal expectation measured by the SFP effect on ITPC affects non-motor activity. Also, the long latencies after the presentation of standards at which modulations of the SFP were largest (the median time of the largest peak of the regression coefficients was 292 ms, Section 2.5) argues against a relation between the temporal expectation measured by SFP effect on ITPC and early sensory stages. Lastly, that the SFP effect on ITPC was substantially stronger for the visual than for the auditory standard modality (as indicated in Section 2.2, Figure 3, and Table A.6, the number of clusters with a large proportion of models significantly different from the intercept-only models was larger for the visual standard modality and, as shown in Section 2.4, Figure 3, and Tables A.6 and A.5, the number of clusters showing significant correlations between models’ decodings and behavioral measures, as well as the strength of these correlations, was larger for the visual than the auditory standard modality) suggests that the effects of the temporal expectation measured by the new foreperiod effect may be specific to the visual modality. Note, however, that the weakness of the SFP effect on ITPC on the auditory modality could be due to the fact that our 32-channel EEG recording system may not have covered auditory brain regions with sufficient density.

### 3.2 Relations to previous investigations

Previous investigations have shown that brain regions associated with clusters whose ITPC activity is significantly correlated with detection performance (colored cells in Table A.5) or with reaction speed (colored cells in Table A.6) are related to temporal expectancy, as discussed next.

Electrophysiological correlates of temporal expectation generated by the foreperiod effect have mainly been investigated through the CNV. Studies demonstrating that the CNV amplitude is maximal at the vertex (Walter, 1967), that the the supplementary motor area (SMA) and the anterior cingulate cortex (ACC) are the sources of the CNV’s O wave (Cui et al., 2000; Zappoli et al., 2000; Gomez et al., 2001), and that the premotor cortex appears to give rise to the CNV’s E wave (Hultin et al., 1996), indicate the central brain region associated with cluster 19 is related to attention orienting and response preparation. Also, more recent studies have shown that posterior sites contribute to the generation of the E wave of the CNV (Gomez et al., 2001, 2003). Of particular interest are the asymmetric activations in the left middle-occipital cortex observed by Gomez et al. (2003, Figures 2 and 3). These studies suggest that activity in the left parieto-occipital region associated to cluster 04 (Figure A.9) may be related to perceptual anticipation.

Findings from a CNV study focused on attention in time by Miniussi et al. (1999) are in close agreement with results reported here. In this study subjects were instructed to detect a target stimulus as quickly as possible. Two central cues which predicted if the subsequent target would occur 600 or 1400 ms after cue onset. Their analysis showed that the influence of the foreperiod duration on event-related potentials elicited by the cue changed over time. The earliest significant influence appeared 280 ms post-cue and occurred in left-parieto-occipital electrodes, and by 480 ms post-cue this difference has moved to the vertex (Figure 7 in Miniussi et al., 1999). Thus, the significant correlation between the strength the SFP effect on ITPC and error rates observed in the left-parieto-occipital cluster 04 and the central-midline cluster 19 (Figure A.9) agrees with the significant influences observed in Miniussi et al. (1999) in left-parieto-occipital and vertex electrodes. In addition, the observation that the SFP effect on ITPC occurred earlier in occipital than in central clusters (Figure 6b) agrees with the finding by Miniussi et al. (1999) that significant foreperiod influences appeared earlier in left-parieto-occipital electrodes than in electrodes surrounding the vertex.

**Figure 7:**
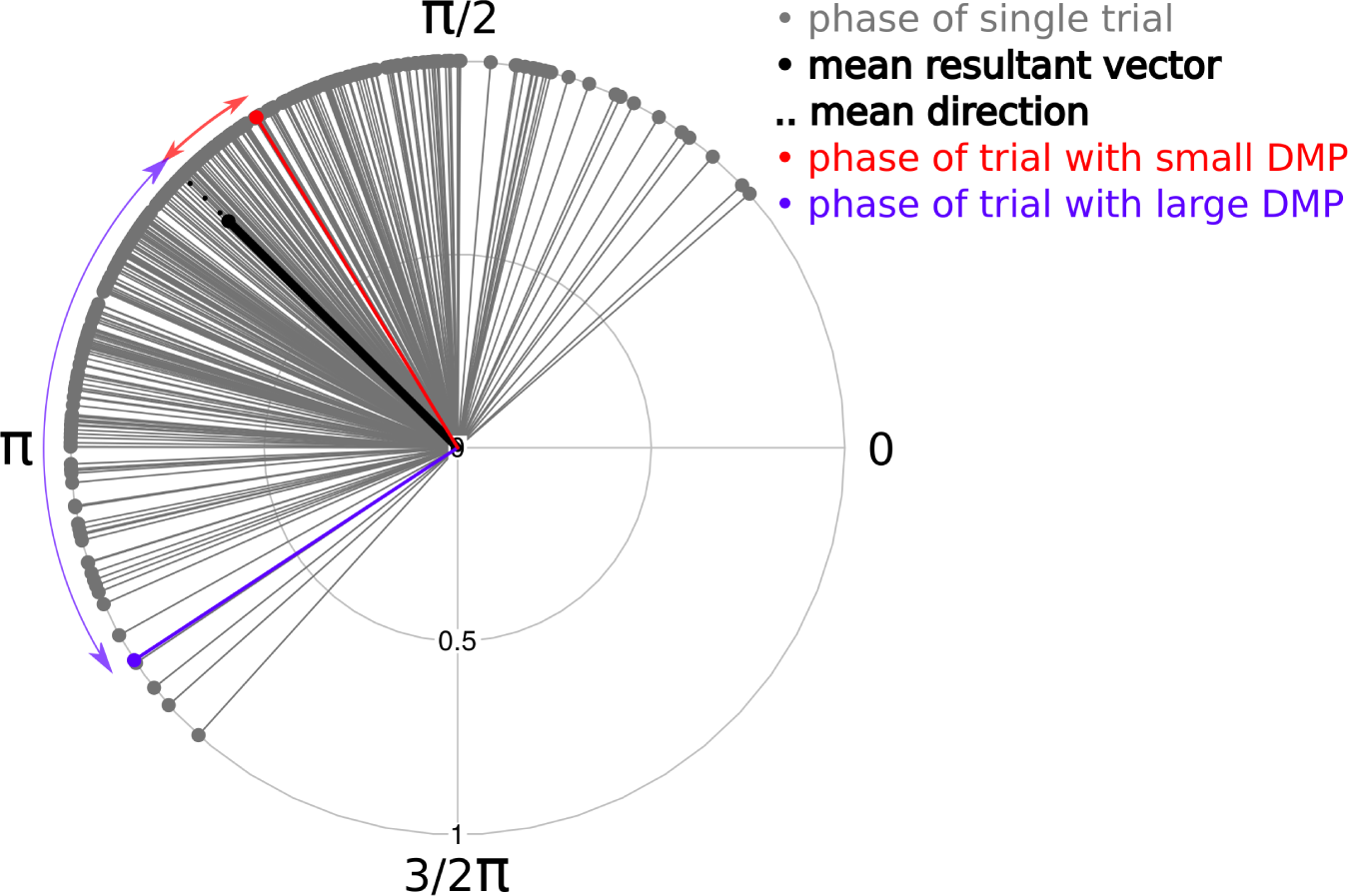
Computation of the DMP. Given a set of phases, represented by gray unit vectors, one first calculates the mean of these unit vectors (the mean resultant vector, black solid vector) and extracts the phase of this mean vector (mean direction, Eq. A.6, black dotted unit vector). Then the deviation from the mean phase for a given phase (e.g., the phase corresponding to the red unit vector) is a measure of distance (i.e., circular variance, Eq. A.5) between this phase and the mean direction (e.g., red arc). The red and blue unit vectors correspond to phases with small and large DMPs, respectively.

Fixed temporal expectations of when an event is likely to occur are marked by activity in left premotor and parietal areas. However, if the event has still not appeared by the expected delay, the right prefrontal cortex updates current temporal expectations (Coull, 2009). The left prefrontal cluster 18 and left parietal cluster 5 (Figure A.9) may be related to correspondingly left-lateralized fMRI activations in attention on time tasks (e.g., Coull and Nobre, 1998). Updates of temporal expectations occurred in the present study, since it used variable foreperiod durations. Previous lesion (Vallesi et al., 2007) and fMRI (Vallesi et al., 2009; Coull et al., 2000) studies have shown involvement of activity in the right prefrontal cortex related to updating temporal expectations in variable foreperiod duration tasks. Thus, we expected to observe a significant correlation between the strength of the SFP effect on ITPC and reaction times in the right prefrontal cluster 13 (Figure A.9). An early fMRI study on attention on time compared brain regions activated by spatial and temporal attention (Coull and Nobre, 1998). This and other studies (e.g., Coull et al., 2004) found a region next the left insula only activated during temporal attention. Interestingly, many of the ICs in cluster 7 are close to the insula (Figure A.9).

Cravo and colleagues specifically studied how temporal expectation modulates the ITPC of neural oscillations in the EEG of humans. In a first study, Cravo et al. (2011) asked if temporal expectations influenced low-frequency rhythmic activity in the theta range (4-8 Hz). They found that temporal expectations modulate reaction times simultaneously with the amplitude, ITC, and phase-amplitude coupling in central-midline electrodes. Interestingly, they reported increased values of ITC at times of higher expectancy. The authors interpreted these modulations as reflecting motor preparation. Our finding of a significant correlation between the SFP effect on ITPC and detectability in the central cluster 19 (Figure 5a) is consistent with the report in Cravo et al. (2011). But, as we discussed above, we believe the modulations of ITPC observed in this manuscript are related to non-motor neural processes. The study by Cravo and colleagues used three foreperiod durations between a warning signal and a Go/No-go target. Differently from previous and from this study (see below), the authors found no modulations of ITC with the foreperiod duration. A possible explanation for this difference is that Cravo and colleagues averaged ITCs across subjects before testing for modulations by the foreperiod duration. These modulations may have been present at the single subject level, but obscured in the average, which highlights the relevance of avoiding averages across subjects. In a second study Cravo et al. (2013) studied the effect of phase entrainment by repetitive visual stimulation of rhythmic activity in the delta range (1-4 Hz) over occipital cortex in an orientation discrimination task. They showed that 150 ms before the presentation of a target stimulus the phase of delta oscillations clustered around the phase corresponding to the largest contrast gain. Considering the effects of expectancy in central brain regions in the Go/No-Go task reported by Cravo et al. (2011), and those in occipital brain regions in the orientation discrimination task reported in Cravo et al. (2013), the authors argued that “the sites where temporal expectation modulate low-frequency oscillations … are highly dependent on the task demands.” However, in Cravo et al. (2011) effects were only studied, or at least reported, over central regions, while in Cravo et al. (2013) effects were only reported over occipital regions. It is possible that extending their analysis to multiple brain regions could show effects over occipital areas in Cravo et al. (2011) and over central areas in Cravo et al. (2013) and invalidate their argument. This highlights the relevance of reporting results over all brain regions.

In an MEG investigation of the effect of attention on primary visual cortex activity, Yamagishi et al. (2008) showed that ITC over a left-occipital brain region was significantly correlated with subjects’ orientation discrimination ability when attention was oriented toward, but not away from, a grating located on the right visual field. The authors concluded that the phase-resetting measured by ITC reflects an attention- and vision-related top-down modulation to primary visual cortex. Using a different brain-imaging modality, the investigation by Yamagishi et al. (2008) supports our previous finding that ITPC is related to behavioral performance, and the finding of part II of this manuscript that ITPC is modulated by attention. Also, it is an interesting coincidence that Yamagishi et al. (2008) found significant correlations between ITC and behavior in four time frequency points between 200 and 300 ms after an attention-orienting cue, and that we found the median of the largest SFP effect on ITPC at 292 ms (Section 2.5). However, our study and that of that Yamagishi et al. (2008) differ in an important regard; they used a trial-averaged measure of ITPC while we used a single-trial one. This single-trial analysis allowed us to find a new foreperiod effect, which shows that ITPC is not constant across trials, but varies systematically as a function of the SFPD. To determine if in our study ITC was also related to behavioral performance, we correlated the peak ITC value following the presentation of a standard (measured as indicated in Section A.5.4) with error rates and mean reaction times for all subjects, standard modalities, and attended modalities. Details of this investigation are reported in Section A.2, Figure A.2, and Tables A.1 and A.2. Similarly to Yamagishi et al. (2008), we found that in the left-parieto-occipital cluster 04 ITC was significantly correlated with detection performance for attended, but not unattended, visual stimuli. However, in the right-parieto-occipital cluster 15 we found a significant correlation between the ITC triggered by attended auditory standards and detection performance, and in the central parieto-occipital cluster 09 we found a significant correlation between the ITC triggered by unattended auditory standards and detection performance. These results appear to contradict the conclusions from Yamagishi et al. (2008) that phase reset over occipital regions reflects vision-related top-town modulations and that these modulations are related to attention. However, this contradiction may be due to differences between our experiments (our temporal-versus their spatial-attention experiments, or our detection versus their discrimination experiments) or differences in the selection of the time-frequency bin used to compute ITC (we used the subject-, standard modality-, and attended modality-specific peak ITC frequency while Yamagishi et al. (2008) used the same time-frequency point for all subjects). The previous significant correlations between peak ITC values and subjects’ error rates were found after computing 56 correlations. None of these correlations survived the multiple-comparison adjustment (Section A.5.9). Therefore, some of the these significant correlations could have appeared by chance. However, we suspect that not all of them are spurious since eight of the nine unadjusted significant correlations were negative. Assuming that these correlations occurred by chance, the probabilities of obtaining positive and negative correlations should be equal. Under this assumption, the probability of obtaining eight out of nine significant correlations is very small (p < 0.02). In our study, the significant correlations between ITC and behavior and those between the SFP effect on ITPC and behavior appear to reflect different neural processes since, with the exception of the correlation with error rates in the left parieto-occipital cluster 04, correlations with ITC occurred in different clusters, standard modalities, and attended modality, than those with the SFP effect on ITPC (compare daggers in Figures 3 and A.2). For example, most significant correlations between ITC and error rates error rates corresponded to auditory standards (Figure A.2 and Table A.1), while those between the SFP effect on ITPC and error rates corresponded to visual standards (Figure 3 and Table A.5).

**Figure A.1:**
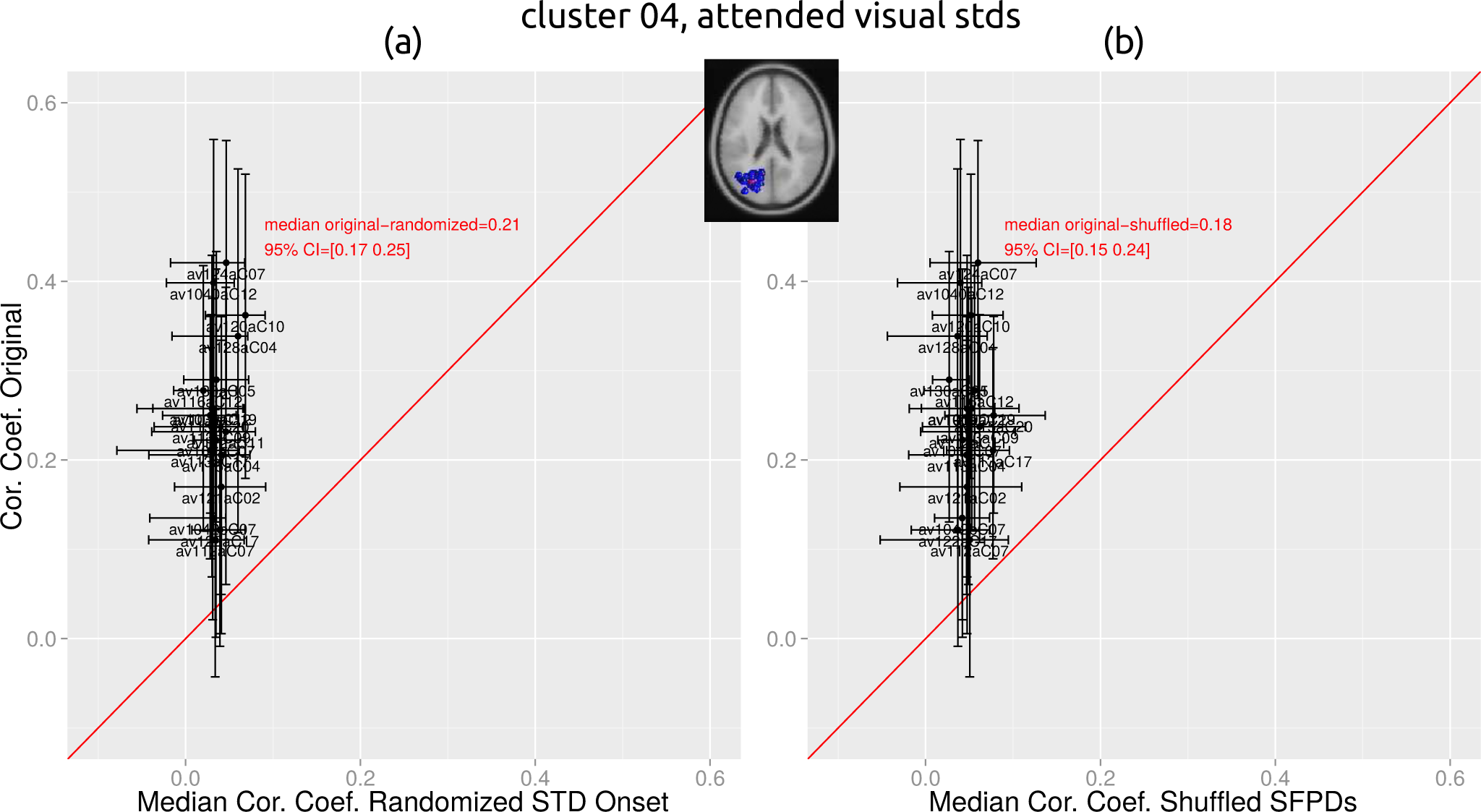
Controls on the SFP effect on ITPC. Panel (a) plots correlation coefficients for models fitted to original datasets, with epochs aligned to the presentation time of standards, versus correlation coefficients for models fitted to surrogate datasets, with epochs aligned to random times. Panel (b) is as panel (a) but for surrogate datasets with SFPDs shuffled among trials. Both panels correspond to the left parieto-occipital cluster 04 and attended visual standards. Correlations for models fitted to original datasets are significantly larger than those from models fitted to surrogate datasets, supporting our assumption that in the SFP effect on ITPC the ITPC triggered by standards is the modulated variable (panel (a)) and that the SFPD is the modulating variable (panel (b)).

**Figure A.2:**
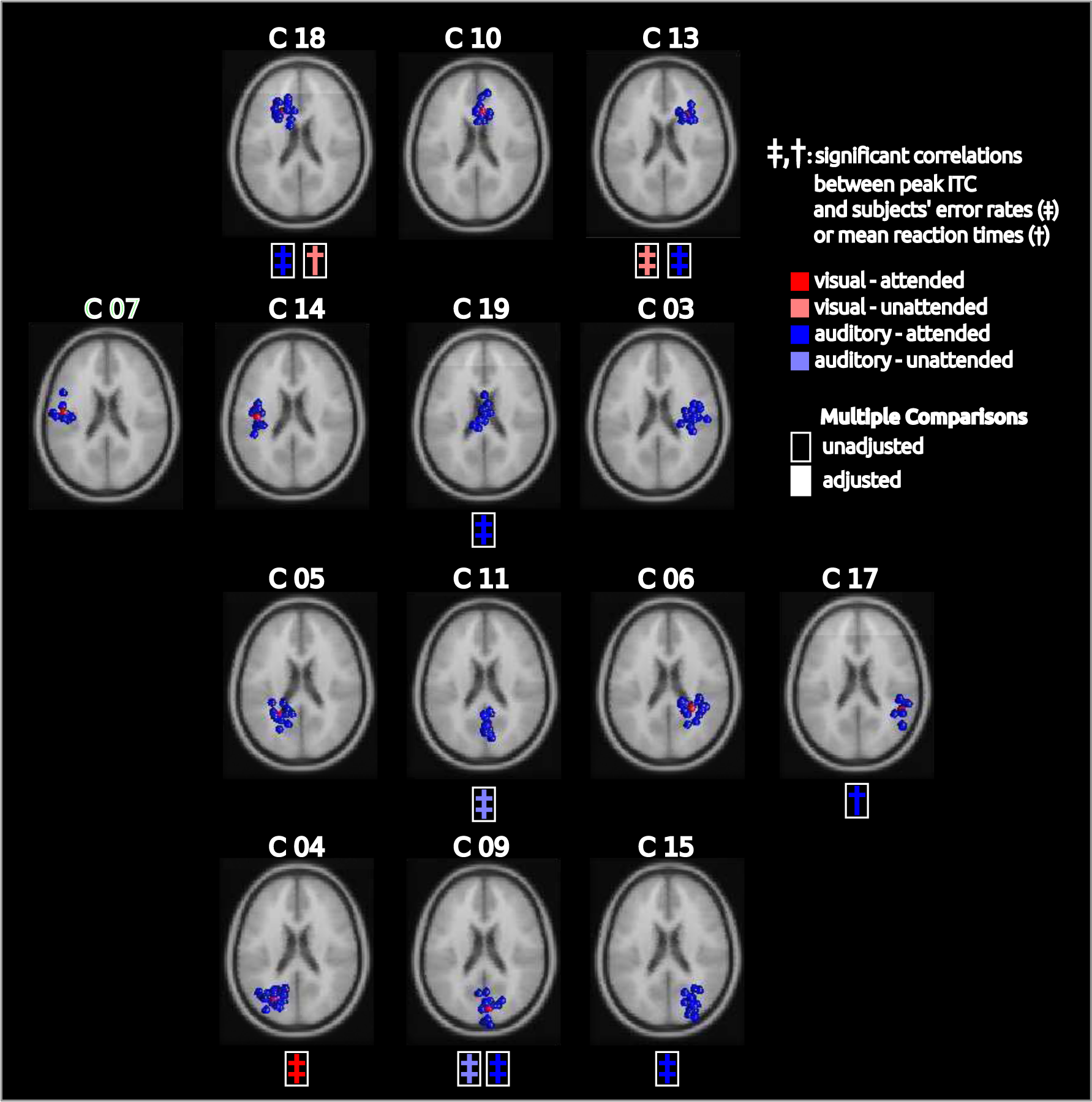
Clusters of ICs, and correlations between peak ITC values and subjects’ behaviors. A dagger (double dagger) shows a significant correlation, unadjusted for multiple comparisons, between peak ITC values and subjects’ mean reaction times (error rates). With the exception of the left parieto-occipital cluster 04 and attended visual standards, significant correlations between peak ITC values and subjects’ behaviors occur in different clusters, standard modalities and attended modalities, than correlations between models’ decodings and subjects’ behaviors (Figure 3).

**Table A.1:** Correlations between peak ITC values and subjects’ error rates. Each cell shows the correlation coefficient, r, and p-values unadjusted, p, and adjusted, adj p, for multiple comparisons. Blue cells highlight correlations with p<0.05.

| Cluster | Standard Modality | Visual Attention |  |  | Auditory Attention |  |  |
| --- | --- | --- | --- | --- | --- | --- | --- |
| | | $r$ | $p$ | $adj\_p$ | $r$ | $p$ | $adj\_p$ |
| 03 | Visual | -0.36 | 0.1624 | 1.00 | 0.35 | 0.2162 | 1.00 |
|  | Auditory | -0.51 | 0.0656 | 0.99 | 0.15 | 0.6132 | 1.00 |
| 04 | Visual | -0.53 | 0.0382 | 0.96 | -0.14 | 0.5882 | 1.00 |
|  | Auditory | 0.52 | 0.0678 | 0.99 | -0.32 | 0.1938 | 1.00 |
| 05 | Visual | -0.36 | 0.1944 | 1.00 | 0.13 | 0.6906 | 1.00 |
|  | Auditory | -0.24 | 0.4678 | 1.00 | -0.38 | 0.2162 | 1.00 |
| 06 | Visual | 0.03 | 0.9206 | 1.00 | -0.01 | 0.9718 | 1.00 |
|  | Auditory | -0.23 | 0.4138 | 1.00 | -0.35 | 0.1852 | 1.00 |
| 07 | Visual | -0.49 | 0.1682 | 1.00 | 0.20 | 0.6042 | 1.00 |
|  | Auditory | -0.39 | 0.2816 | 1.00 | -0.33 | 0.3748 | 1.00 |
| 09 | Visual | 0.03 | 0.9092 | 1.00 | -0.34 | 0.2234 | 1.00 |
|  | Auditory | -0.62 | 0.0190 | 0.80 | -0.58 | 0.0278 | 0.90 |
| 10 | Visual | -0.38 | 0.2706 | 1.00 | 0.31 | 0.4092 | 1.00 |
|  | Auditory | -0.13 | 0.7218 | 1.00 | -0.10 | 0.7738 | 1.00 |
| 11 | Visual | 0.12 | 0.7058 | 1.00 | 0.08 | 0.7788 | 1.00 |
|  | Auditory | -0.69 | 0.0130 | 0.67 | -0.29 | 0.3292 | 1.00 |
| 13 | Visual | -0.24 | 0.4756 | 1.00 | 0.77 | 0.0056 | 0.40 |
|  | Auditory | -0.34 | 0.3394 | 1.00 | -0.72 | 0.0162 | 0.75 |
| 14 | Visual | -0.50 | 0.0854 | 1.00 | -0.39 | 0.2208 | 1.00 |
|  | Auditory | -0.27 | 0.3572 | 1.00 | -0.31 | 0.3626 | 1.00 |
| 15 | Visual | 0.25 | 0.3156 | 1.00 | -0.02 | 0.9458 | 1.00 |
|  | Auditory | -0.35 | 0.2036 | 1.00 | -0.54 | 0.0196 | 0.81 |
| 17 | Visual | -0.25 | 0.5424 | 1.00 | -0.06 | 0.8630 | 1.00 |
|  | Auditory | 0.48 | 0.2202 | 1.00 | -0.02 | 0.9368 | 1.00 |
| 18 | Visual | -0.34 | 0.2390 | 1.00 | -0.03 | 0.9120 | 1.00 |
|  | Auditory | -0.21 | 0.4862 | 1.00 | -0.67 | 0.0242 | 0.87 |
| 19 | Visual | 0.15 | 0.5896 | 1.00 | 0.18 | 0.5638 | 1.00 |
|  | Auditory | -0.04 | 0.9056 | 1.00 | -0.85 | 0.0010 | 0.11 |

**Table A.2:** Correlations between peak ITC values and subjects’ mean reaction times. Same format as Table A.1.

| Cluster | Standard Modality | Visual Attention |  |  | Auditory Attention |  |  |
| --- | --- | --- | --- | --- | --- | --- | --- |
|  |  | r | p | adj-p | r | p | adj-p |
| 03 | Visual | -0.22 | 0.4036 | 1.00 | 0.25 | 0.3388 | 1.00 |
|  | Auditory | -0.46 | 0.0602 | 0.97 | -0.09 | 0.7218 | 1.00 |
| 04 | Visual | -0.08 | 0.7416 | 1.00 | 0.00 | 0.9882 | 1.00 |
|  | Auditory | 0.02 | 0.9322 | 1.00 | -0.28 | 0.2512 | 1.00 |
| 05 | Visual | 0.15 | 0.5894 | 1.00 | 0.41 | 0.1536 | 1.00 |
|  | Auditory | -0.06 | 0.8322 | 1.00 | -0.19 | 0.5250 | 1.00 |
| 06 | Visual | -0.28 | 0.2808 | 1.00 | 0.00 | 0.9882 | 1.00 |
|  | Auditory | -0.23 | 0.4088 | 1.00 | -0.33 | 0.2104 | 1.00 |
| 07 | Visual | -0.04 | 0.9086 | 1.00 | 0.08 | 0.8406 | 1.00 |
|  | Auditory | 0.05 | 0.8878 | 1.00 | 0.03 | 0.9334 | 1.00 |
| 09 | Visual | -0.16 | 0.5782 | 1.00 | -0.05 | 0.8532 | 1.00 |
|  | Auditory | 0.12 | 0.6622 | 1.00 | -0.31 | 0.2766 | 1.00 |
| 10 | Visual | 0.19 | 0.5968 | 1.00 | 0.40 | 0.2500 | 1.00 |
|  | Auditory | 0.39 | 0.2844 | 1.00 | -0.04 | 0.9142 | 1.00 |
| 11 | Visual | -0.21 | 0.4930 | 1.00 | -0.17 | 0.5838 | 1.00 |
|  | Auditory | -0.43 | 0.1400 | 1.00 | 0.26 | 0.3822 | 1.00 |
| 13 | Visual | 0.25 | 0.4262 | 1.00 | 0.54 | 0.0746 | 0.99 |
|  | Auditory | 0.09 | 0.7674 | 1.00 | -0.26 | 0.4130 | 1.00 |
| 14 | Visual | -0.24 | 0.4220 | 1.00 | -0.17 | 0.5694 | 1.00 |
|  | Auditory | -0.27 | 0.3618 | 1.00 | -0.22 | 0.4852 | 1.00 |
| 15 | Visual | -0.23 | 0.3250 | 1.00 | -0.03 | 0.9086 | 1.00 |
|  | Auditory | 0.03 | 0.8986 | 1.00 | -0.23 | 0.3204 | 1.00 |
| 17 | Visual | -0.18 | 0.6160 | 1.00 | 0.06 | 0.8692 | 1.00 |
|  | Auditory | -0.15 | 0.6830 | 1.00 | -0.74 | 0.0250 | 0.78 |
| 18 | Visual | -0.31 | 0.2542 | 1.00 | -0.58 | 0.0240 | 0.76 |
|  | Auditory | 0.15 | 0.5918 | 1.00 | -0.21 | 0.4384 | 1.00 |
| 19 | Visual | -0.39 | 0.1592 | 1.00 | -0.02 | 0.9502 | 1.00 |
|  | Auditory | -0.13 | 0.6522 | 1.00 | -0.17 | 0.5450 | 1.00 |

An investigation by Busch et al. (2009) also showed that stimulus detectability is related to prestimulus phase. Below we highlight a few differences between this study and ours. The purpose of this comparison is not to belittle the research by Busch et al. (2009), which we believe was excellent, but to emphasize a few novel characteristics of our analysis. First, the ITC and the phase coherence index used in the main text of Busch et al. (2009) are phase measures for groups of trials, while the DMP used in this article is a phase measure for single trials. Thus, we were able to decode the SFP duration from DMP values in a trial-by-trial basis, while these single-trial decodings are not possible with phase measures for groups of trials. In the supplementary Figure 2, Busch et al. (2009) reported the results of using a classifier to predict in a single trial if a stimulus will be detected based on the phase at a single time-frequency point. The fact that, after averaging the predictions of the classifiers of different subjects, they found a correlation between the predictions of the classifiers and the detection hit ratio is remarkable. However the analysis method is limited by using only one time-frequency point to predict the detectability of a stimulus. The models used in this manuscript used a 500 ms window of phase values to decode the SFP duration. That is, the second difference between the study by Busch et al. (2009) and ours is that they used univariate models, while we used multivariate ones. Note that we have optimized models to decode SFP durations, and then found significant correlations between the decoding power of these models and behavior without fitting model parameters to behavioral data. That a model optimized to predict behavior from a stimulus feature (e.g., ITPC) achieves reliable predictions does not necessarily mean that the stimulus feature is relevant to the behavior. A sufficiently complex model can predict arbitrarily closely any behavior from an irrelevant stimulus feature, if the model overfit the data (Geman et al., 1992). Although there exist methods to minimize the risks of overfitting, such as using separate pieces of data to estimate the parameters of a model and to evaluate its predictive power (the method used in Busch et al. (2009)), this risk never disappears when optimizing a model to predict behavior. Differently, the procedure used in this article to show that ITPC is related to behavior does not suffer from the overfitting problem, since no model parameter was fitted to behavioral data. The third difference is that the study by Busch et al. (2009) averaged data across subject while we analyzed different subjects separately. This single-subject analysis allowed to observe that those subjects whose brain activity was more strongly modulated by the SFP duration were those who performed better Fourth, Busch et al. (2009) reported the analysis of a single electrode (Fz), while our study described the analysis of the activity of ICs over the whole brain. As we discussed above in relation to the research by Cravo and collaborators, we believe it is important to report effects across the whole brain to show how hypothesis supported by activity in a given brain region generalize across the brain. Fifth, Busch et al. (2009) analyzed activity from EEG channels, while the present study characterized the activity of ICs. Applied to EEG, ICA finds maximally independent sources generating recorded potentials. Scalp representations of these sources resemble fields generated by current dipoles inside the brain, and biological arguments suggest that these maximally-independent sources reflect the synchronized activity of neurons in compact cortical regions (Delorme et al., 2012). Thus, our analysis most probably characterized the activity of cortical sources while that of Busch et al. (2009) described the activity of mixtures of these sources. Sixth, the study by Busch et al. (2009) characterized phase correlates of visual processing, while our study investigated phase correlates of both visual and auditory processing. The multi-modal nature of our experiment allowed us to see that the SFP effect on ITPC was substantially stronger in the visual than in the auditory modality. An important advantage of the study by Busch et al. (2009) over ours is that, for each subject separately, they calibrated the visual stimuli so that the detection rate of the subject was close to 50%. No such individualized calibration was performed in our experiment, and the averaged detection rate across subjects was 86%. Having a similar number of trials where subjects perceived and failed to perceive the stimulus facilitates the estimation of the phase correlates of stimulus perception. However, the fact that the methods used in this manuscript could find reliable phase correlates of stimulus detection with sub-optimal stimulation shows that these methods could be applied to a larger number of EEG experiments where stimulation was not optimized for each subject.

### 3.3 What neural processes generate the SFP effect on ITPC?

What brain processes can make the phases of standards closer to the warning signal more aligned at some times but more misaligned at other times than the phases of of standards further away from the warning signal? Also, how can it happen that phase alignment fluctuates in an oscillatory manner, as indicated in Figures 4a-c and A.7? Large values of ITPC are typically associated with phase resets, like those that occur upon the presentation of visual stimuli (e.g., Makeig et al., 2002). But a single phase-reset event is not the answer to the previous question, since after it all trials should have a similar phase, so that there would be no difference in ITPC between trials closer to and further away from the warning signal. A single phase-reset event can neither explain the fluctuations in ITPC. These fluctuations are reminiscent of known variations in perception and reaction times that have been related to the phase of low-frequency neural rhythms (Bartley and Bishop, 1932; Bishop, 1932; Jarcho, 1949; Lindsley, 1952; Lansing, 1957; Callaway and Yeager, 1960; Dustman and Beck, 1965; Trimble and Potts, 1975; Valera et al., 1981; Mathewson et al., 2009; Busch et al., 2009; Mathewson et al., 2012; Busch and VanRullen, 2010; Bonnefond and Jensen, 2015; Liu et al., 2014; Drewes and VanRullen, 2011; Thorne et al., 2011; Fiebelkorn et al., 2011; Chakravarthi and 2012; Cravo et al., 2015; Milton and Pleydell-Pearce, 2016), reviewed in VanRullen and Koch (2003). However, if after a reset the phase evoked by standards oscillated identically across trials there would not be a difference in ITPC between trials closer and further-away from the warning signal.

A brain mechanism generating the above effects should be able to replicate the fluctuations in ITPC shown in Figures 4a-c and A.7. We make three observation: First, these fluctuations appear to oscillate at a very low frequency (i.e., less than 1 Hz), independently of the frequency at which the phase of trials was measured (the latter frequency is that of the cosine of the mean phase given by the dashed curve in Figures 4a-c and A.7). Coincidentally, fluctuations at 1 Hz in visual detectability have been reported by Fiebelkorn et al. (2011). Also, fluctuations in somatosensory detectability (between 0.01 and 0.1 Hz) have been described by Monto et al. (2008). Second, in some figures the phase of the oscillations in ITPC at time zero differs between early and late trials (e.g., the phase of the blue curve corresponding to early trials in Figure 4a is more advanced than that of the red curve corresponding to late trials). Third, as shown in Figure 5, these fluctuations are related to subjects’ behaviors.

We propose a simple model for the generation of low-frequency oscillations that can account for the previous observations. The component of the recorded potential corresponding to trial *i*, at frequency *f*, and time t is given by Eq. 1:

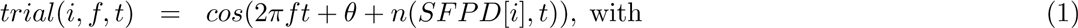

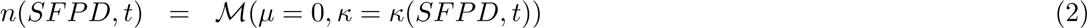

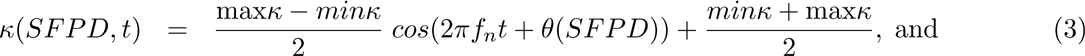

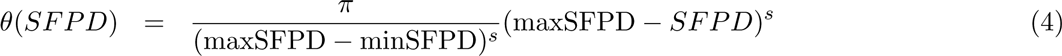

To account for the first previous observation, we assume that the phase of the cosine is contaminated by an additive noise *n* (Eq. 2) following a von Mises distribution with a precision parameter, *κ*, varying sinusoidally on time (Eq. 3). When the precision of this noise is small and large we should observe decoherent and coherent activity, respectively, and, because the noise precision varies sinusoidally in time, we should observe alternations between coherence and decoherence, as shown in Figures 4a-c and A.7. To account for the second previous observation, we make the phase at time zero of the precision of the noise vary smoothly as a function of the SFPD (Eq. 4). To account for the third observation, we speculate that subjects achieving better detection performance were those showing larger modulations in the precision of the noise. For these subjects the ITPC should be more different between trials closer to and further away from the warning signal, and models should more reliably decode the SFPD from ITPC activity, in agreement with Figure 5. The parameters of the previous model are the frequency, *f*, the noiseless phase, *θ*, and the vector of SFPDs, *SF P D*, in Eq. 1; the minimum and maximum values, *minκ* and *maxκ*, and the frequency, *f_n_*, of the noise precision in Eq. 3; and the steepness of the change in the noise precision phase as a function of the SFPD, *s*, in Eq. 4.

To validate the previous speculations we simulated data following the model in Eq. 1 (Figure A.3) and fitted a decoding model to this data, as described in Section A.3. With this simulated data we obtained oscillations in averaged DMP (Figures A.4a, b), as in Figure 4a-c. We also attained oscillations in the difference between the averaged DMP between trials closer to and further away from the warning signal (Figures A.4c, d), as in Figure 4d-f. In addition, we verified that a model fitted to data with larger fluctuations of the precision of the noise generated more accurate decodings than those of a model fitted to data with smaller modulations of this precision, supporting the previous argument on the relation between fluctuation of ITPC and subjects’ behaviors. As explained in Section A.3, the correlation coefficient for a model fitted to data with smaller fluctuations of the precision of the noise was r = 0.23 (95% CI=[0.12, 0.34]), which was significantly smaller than that for a model fitted to data with larger fluctuations of the precision of the noise r = 0.51 (95% CI=[0.43, 0.59]).

**Figure A.3:**
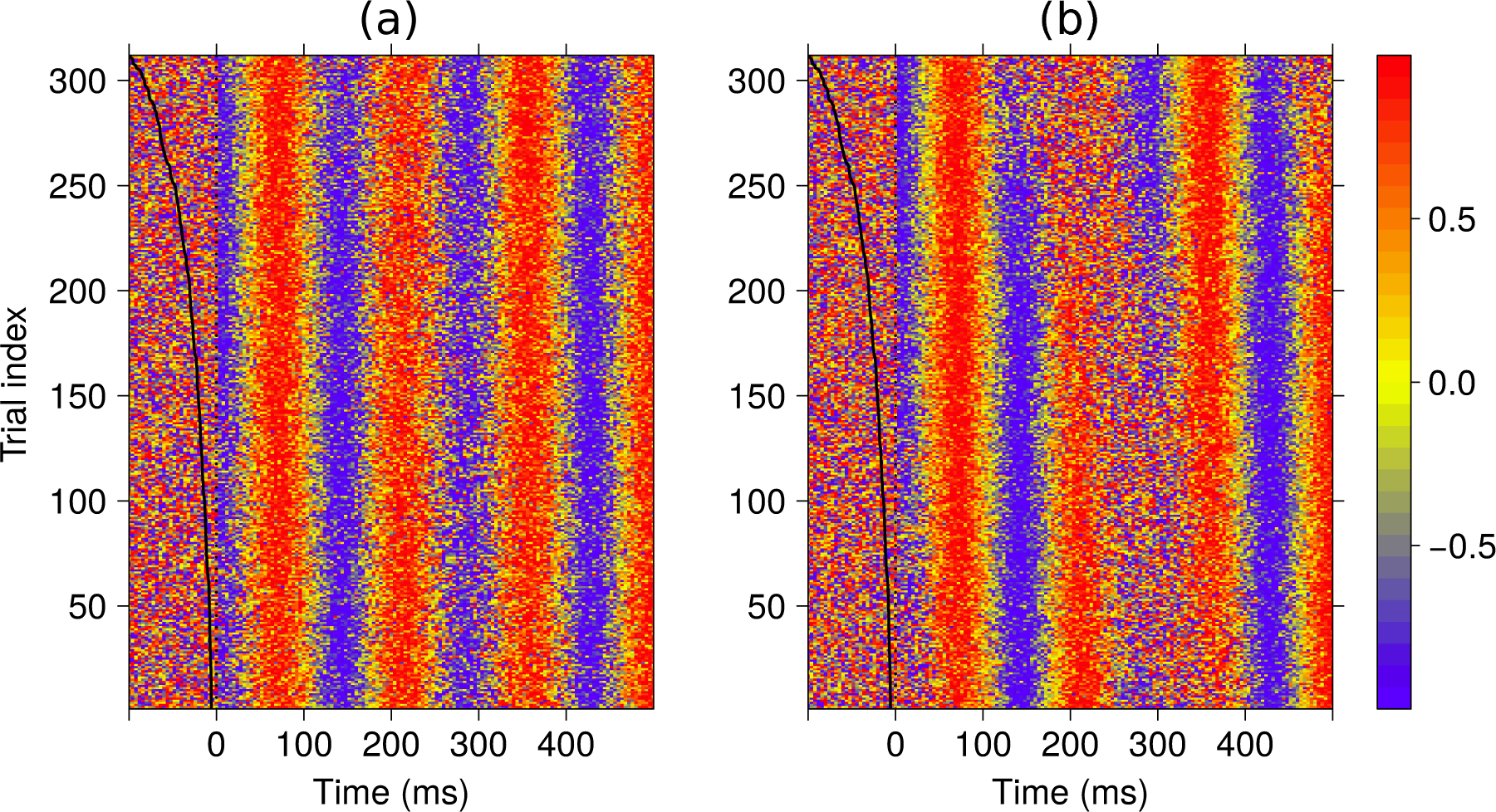
Oscillations simulated according to Eq. 1 with phase noise *n* having small (*minκ* = 1.51, *maxκ* = 3.5, panel (a)) and large (*minκ* = 0.01, *maxκ* = 5.0, panel (b)) fluctuations in precision (Eq. 3).

These results show that a simple sinusoidal oscillation with a noise process added to its phase captures salient features of the SFP effect on ITPC, suggesting that this model is a good first description of how the SFPD modulates the phase of neural oscillations. Key properties of this noise process are its precision oscillating in time and its SFPD-dependent phase at time zero.

### 3.4 Single-trial predictive methods for EEG

Single-trial predictive models are seldomly used with EEG recordings to characterize the neural basis of psychophysical process (but see Sajda et al., 2009; Pernet et al., 2011b). These models are mostly applied to EEG recordings in brain computer interface applications (e.g., Nicolas-Alonso and Gomez-Gil, 2012; Wolpaw et al., 2002). However, these applications are usually focused on the predictive power of these models, and not so much on the brain mechanisms underlying the psychophysical process they attempt to predict. This omission is striking, since single-trial predictive models have been very successful for understanding the neural basis of psychophysical processes using fMRI recordings (e.g., Haxby et al., 2001; Haynes and Rees, 2005; Kamitani and Tong, 2005; Kay et al., 2008; Naselaris et al., 2009; Serences and Boynton, 2007; Reey et al., 2010; Polyn et al., 2005; Mitchell et al., 2008; Abrams et al., 2011; Hampton and O’Doherty, 2007). For reviews see Naselaris et al. (2011); Tong and Pratte (2012). Note that applying single-trial predictive methods to EEG recordings offers the possibility of understanding the neurophysiological basis of behaviors that occur at fast time scales, which cannot be studied with typical fMRI recordings. For example, it would have been impossible to find that the SFPD modulates the ITPC triggered by standards since these modulations occur in a 500 ms-long time window that cannot be resolved with the two seconds sampling rate of typical fMRI experiments.

LIMO EEG (Pernet et al., 2011a) is an existing single-trial predictive method for EEG recordings. It attempts to predict the activity of electrodes using experimental factors as inputs. The method described in this manuscript differs from the previous one in that the EEG recordings are used as inputs to predict an experimental factor (i.e., the SFP duration). That is, LIMO EEG is an encoding method, aiming at describing how experimental factors are encoded in EEG activity, while the method introduced here is a decoding one, seeking to predict experimental factors from EEG activity. An advantage of decoding over encoding methods is that they can be readily used to determine that a certain pattern of brain activity (e.g., ITPC) is related to a particular behavioral characteristic (e.g., reaction times). This can be done by using the decoding method to decode an experimental factor that is related to the particular behavior (e.g., the SFPD is related to reaction times). Then the decoding of the method should be related to the behavior, as we showed in this article. On the contrary, is not straightforward to relate patterns of brain activity and behavior with encoding methods. Another advantage of the method used here compared to LIMO EEG is that the former method is a multivariate test, while the latter uses univariate tests. Both methods can be used to test if a scalar trial attribute (e.g., the SFPD) is related to a time-varying trial attribute (e.g, the ITPC at time t). However, the method introduced here uses information across all time points to test if a time-dependent trial attribute at a given time is related to the scalar trial attribute, while LIMO EEG only uses information at one time point to test if the time-dependent trial attribute at that point is related to the scalar trial attribute. If the time-varying trial attribute is correlated across time (as happens with most EEG time-varying attributes) a multivariate method should be more powerful than a univariate one.

The construction of predictive methods for EEG data requires to address two signal processing problems: the curse of dimensionality and multicollinearity of the EEG regressors. The curse of dimensionality refers to the exponential increase of the volume of a space as its dimensionality grows (Bellman, 1961). In predictive models the dimensionality of the input space is given by the number of predictors. To estimate accurate models one should use training data densely covering the volume of the input space. However, if the input space is very high dimensional, collecting enough training data to densely cover the “exponentially large” volume of the input space becomes unfeasible. For example, in EEG predictive methods one would like to use all EEG samples in an interval preceding an event of interest (i.e., presentation of a standard) as regressors to predict an stimulus property (i.e., the SFP duration). However, if this interval is long and/or the EEG sampling rate is large, the number of regressors will be very large and the predictive method will become impractical, due to the curse of dimensionality. Multicollinearity in the inputs is another important problem in predictive models because, among other things, it yields parameters estimates with large variance (Belsley et al., 2004), which in turn limits the inferences that can be made from these estimates.

The predictive method used in this article was a simple one. To address the previous signal processing problems we used a regularized linear regression model (Section A.5.7) and estimated its parameters using a variational method (Section A.5.8). We have developed more elaborated predictive methods for the characterization of responses of visual cells from their responses to (correlated) natural stimuli (Rapela et al., 2006, 2010). These are nonlinear methods specifically designed to address the curse of dimensionality and multicollinearity problems. Using nonlinear predictive models, carefully validated with behavioral data, to characterize EEG recordings could generate superior predictions and be able to model a broader range of psychophysical processes than those achievable with linear models. We will apply these more advanced predictive model to EEG recordings in future investigations.

In summary, the decoding method used in this article is just a first step in using EEG activity to decode behavior and there exist ample possibilities for further improvements.

### 3.5 Future work

In future investigations we will perform new experiments to test how psychophysical factors modulate the SFP effect on ITPC. For this we will build on thoughtful investigations on the psychophysical factors associated with the CNV. Is the SFP effect on ITPC related to expectancy (Walter et al., 1964)? Would it disappear if responses to deviants are not required? Would it be attenuated if a proportion of deviants do not require response? Is the SFP effect on ITPC linked to motor preparation (Gaillard, 1977, 1978)? Does the emergence of this effect requires motor responses? What is the relation between the SFP effect on ITPC and intention to respond (Low et al., 1966)? Would the strength of this effect be directly proportional to the anticipated force needed in the motor response? Is the SFP effect on ITPC related to motivation (McCallum and Walter, 1968)? Would the strength of this effect be augmented when subjects are instructed to concentrate hard and respond quickly to deviants? How does this effect relate to arousal (Tecce, 1972)? Would the relation between arousal and this effect be U-shaped, as is the relation between arousal and CNV amplitude (Tecce, 1972, Figure 6b)? Is the SFP effect on ITPC linked to to explicit (Macar and Vidal, 2003; Pfeuty et al., 2003, 2005) or implicit (Praamstra et al., 2006) time perception? Would this effect reflect the perceived duration of a target interval when subjects compare it to the duration of a memorized interval? We will address these questions in future investigations.

## 4 Summary

We claimed that phase coherence is a key aspect of neural activity currently under used for the characterization of EEG recordings. We proposed a single-trial measure of phase coherence (Figure 7) an a method to relate it to experimental variables (Figure 2). Using this measure and method we demonstrated a novel foreperiod effect on single-trial phase coherence. We showed that the new foreperiod effect is not an artifact of the proposed method since it can be observed in simple trial averages (Figure 4). We demonstrated the relevance of the new foreperiod effect by reporting strong correlations between the strength of the effect and subjects’ error rates and mean reaction times (Figure 5). We argued that temporal expectancy plays an important role in the generation of the new foreperiod effect. We hope the present manuscript has provided convincing evidence for the relevance of phase coherence for the characterization of EEG recordings and that it has shown that single-trial decoding methods can provide useful information about brain processes. We anticipate that the new foreperiod effect on phase coherence will lead to as many important insights about anticipatory behavior as the foreperiod effect on reaction times has done until now.

## 5 Methods

### 5.1 Experiment information

We analyzed the experimental data first characterized in Ceponiene et al. (2008). Below we summarize features of this data relevant to the present study; further details are given in the previous article.

#### 5.1.1 Subjects

We only characterized the younger-adult subpopulation in Ceponiene et al. (2008), comprising 19 subjects (11 females) with a mean age of 25.67 ± 5.94 years.

#### 5.1.2 Stimuli

Stimuli were sequentially presented in a visual and an auditory stream (Figure 1a). Auditory stimuli were 100 ms duration, 550 Hz (deviants) and 500 Hz (standards) sine-wave tones. These tones were played via two loudspeakers located at the sides of a 21-inch computer monitor. Visual stimuli were light-blue (deviants) and dark-blue (standards) filled squares subtending 3.3 degrees of visual angle, presented on a computer monitor for 100 ms on a light-gray background. Interspersed among the deviant and standard stimuli were attention-shifting cues. These cues were presented bimodally for 200 ms, by displaying the words HEAR (LOOK) on orange letters on the computer monitor and simultaneously playing the words HEAR (LOOK) on the loudspeakers. The inter-stimulus interval (ISI) between any two consecutive stimuli varied pseudo-randomly between 100 and 700 ms as random samples from short, medium, and large uniform distributions, with supports [100-300 ms], [301-500 ms], and [501-700 ms], respectively. In each block, the ISI of 132, 60, and 72 pseudo-randomly chosen stimuli were drawn from the short, medium, and large ISI distributions, respectively. Stimuli were presented in blocks of 264 for a duration of 158 seconds. Each block consisted of 24 cue stimuli (12 visual) and 240 non-cue stimuli (120 visual). The 120 non-cue stimuli of each modality comprised 24 deviants and 96 standards. A video of the experimental stimuli appears at http://sccn.ucsd.edu/∼rapela/avshift/experiment.MP4

#### 5.1.3 Experimental design

The experiment comprised FOCUS-VISION, FOCUS-AUDITION, and SWITCH blocks. In FOCUS-VISION (AUDITION) blocks subjects had to detect visual (auditory) deviants and ignore attention-shifting cues. In SWITCH blocks these cues became relevant, and after a LOOK (HEAR) cue subjects had to detect visual (auditory) deviants. The type of block was told to the subject at the beginning of each block. Subjects pressed a button upon detection of deviants. Each subject completed four FOCUS-VISION blocks, four FOCUS-AUDITION blocks, and 12 SWITCH blocks. Here we only characterize SWITCH blocks. Correct responses in SWITCH blocks are shown in Figure 1b. After a LOOK (HEAR) cue subjects oriented their attention to the visual (auditory) modality, as indicated by the magenta (cyan) segments in Figure 1b. A warning signal is a stimulus initiating a period of expectancy for a forthcoming deviant. LOOK (HEAR) cues were the warning signals in the SWITCH blocks characterized in this study.

### 5.2 EEG acquisition

Continuous EEG was recorded from 33 scalp sites of the International 10-20 system, with xxx electrodes and a yyy amplifier, and digitized at 250 Hz. The right mastroid served as reference.

### 5.3 Characterized epochs

For each IC of each subject we built four sets epochs, aligned at time zero to the presentation of attended visual standards, attended auditory standards, unattended visual standards, and unattended auditory standards, with a duration of 500 ms (Figure 2c) We call standard modality to the modality of the stimuli used to align a set of epochs (visual or auditory), and we call attended modality to the attended modality corresponding to a set of epochs (e.g., the visual (auditory) modality for epochs aligned to the presentation of attended (unattended) visual standards). The mean number of epochs aligned to the presentation of attended visual standards, attended auditory standards, unattended visual standards, and unattended auditory standards was 150, 136, 99, and 94, respectively. Note that the mean number of epochs aligned to the presentation of attended stimuli was considerably larger than that aligned to the presentation of unattended stimuli, for both the visual and auditory standards. Thus, for testing the influence of attention on the SFPD effect on ITPC (Section 2.4) we equalized the number of epochs used to fit attended and unattended models. For each attended model, we selected a random subset of the attended epochs of size equal the number of epochs used to fit the corresponding unattended model. Figure A.10a displays histograms of deviant latencies, with respect to the presentation time of the previous standard, for all combinations of standard modalities and attended modalities. To avoid possible movement artifacts due to the response to deviants, we excluded from the analysis epochs where deviants of the attended modality were presented in the 500 ms after the presentation of the standard at time zero. The number of deviants excluded from the analysis is given by the sum of counts to the left of the red vertical lines in Figure A.10a. Figure A.10b shows histograms of SFPDs for all combinations of standard modalities and attended modalities. The vertical green line correspond to the mean SFPD, which was shorter for epochs where attention was directed to the visual (1.9 sec, left panels) than to the auditory modality (2.4 sec, right panels). A repeated-measures ANOVA with SFPD as dependent variable showed a significant main effect of attended modality (F(1, 16264)=234.31, p<0.0001) and a posthoc analysis indicated that the mean SFPD was shorter in epochs corresponding to visual than to auditory attention (z=15.31, p<2e-16). Further information on this ANOVA appears in Section A.5.12.

### 5.4 Deviation from the mean phase

Let *θ*_0_ be the phase of a given trial, and *θ*_1_, …, *θ_N_* be the phases of the trials in a reference set, all phases measured at the same time-frequency point. Then, the deviation from the mean phase is a measure of the difference between the phase of the given trial (i.e., *θ*_0_) and the mean phase of all trials in the reference set, (i.e., mean direction, 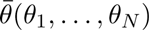, Eq. A.6), as illustrated in Figure 7. The circular variance (*CV*, Eq. A.5) is used to measure this difference:

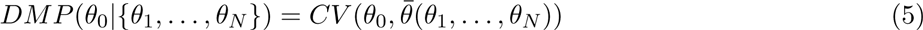

The DMP is zero (one) if the phase of the given trial is equal (opposite) to the mean phase of the trials in the reference set. The DMP has previously been used to investigate interelectrode phase coherence with EEG recordings (Hanslmayr et al., 2007, Figure 2c). It is an appealing measure of single-trial phase coherence since, as demonstrated in Section A.5.6, when there is large phase concentration (i.e., ITC≃ 1) the average value of the DMP approaches the ITC, a widely used measure of averaged ITPC (Section A.5.4). When there is not large phase concentration, the mean phase cannot be estimated reliably, and therefore the DMP becomes unsound. Here we only computed DMP values at times where the distribution of phases across trials was significantly different from the uniform distribution (p<0.01, Raleigh test).

## Acknowledgments

We thank Dr. Steve Hillyard for suggesting Figure 4 and to investigate the influence of the maximum SFPD in the decodings of the models (Figure A.5). We thank Dr. Cyril Pernet for suggesting the use of robust correlation coefficients. We also thank Mr. Robert Buffington for computer assistance. Most of the research in this paper was performed with free software. Computations were done with R (R Core Team, 2012), the article was written in LATEX (Lamport, 1994), figures were prepared with Inkscape (The Inkscape Team, 2004), all in a personal computer and in clusters of personal computers running Linux (Torvalds, 2008). The only paid software used in this investigation was MATLAB (MATLAB, 2013), employed solely for initial preprocessing with ICA (Makeig et al., 1996).

## Glossary

attended modality: the attended modality (visual or auditory) corresponding to a set of epochs (Section 5.3).
DMP: (deviation from the mean phase) single-trial measure of ITPC (Section 5.4).
IC: (independent component) component from an ICA decomposition (Section A.5.1).
ITC: (inter trial coherence) a measure of ITPC in a group of trials (Section A.5.4).
ITPC: (inter trial phase coherence) degree of phase alignment across multiple trials. Here we use the DMP and the ITC as measures of ITPC for single trials and for groups of trials, respectively (Section 1).
SFP: (standard foreperiod) interval between the presentation of a warning signal and a standard stimulus (Section 1).
SFPD: (standard foreperiod duration) duration of a SFP (Section 1).
standard modality: the modality of the standard stimuli (visual or auditory) used to align a set of epochs (Section 5.3).
warning signal: stimulus initiating a period of expectancy for a forthcoming impendent stimulus. The LOOK and HEAR attention-shifting cues are the warning signals in this study (Section 5.1.3).

## A Appendix

### A.1 Controls on the SFP effect on ITPC

The previous sections demonstrated the existence of the SFP effect on ITPC, and argued for its relevance by showing that its strength correlates with subjects’ perceptual abilities and reaction speeds. From the successful decoding of SFPDs from the ITPC following the presentation of standards we inferred that the SFPD (i.e., the modulation variable) modulates the ITPC evoked by standards (i.e., the modulated variable). Below we present evidence supporting the inference that the modulated variable is the ITPC evoked by standard stimuli (Section A.1.1), and that the modulating variable is the SFPD (Section A.1.2).

#### A.1.1 The ITPC triggered by standards is the modulated variable

Besides the presentation of standards, any other experimental event (e.g., the presentation of a warning signal) could have generated ITPC modulations that allowed models to decode, from a 500 ms-long segment of phase coherence starting at time t_0_ after the presentation of a warning signal, that the warning signal was presented t_0_ ms before the start of the segment. We tested this possibility by building surrogate datasets, identical to the original ones, with the exception that epochs were aligned to random times, instead of being aligned to the presentation time of standards, and then fitting decoding models to these surrogate datasets.

For each epoch in an original dataset we built a corresponding epoch in the corresponding surrogate dataset. To construct the surrogate epoch, the onset time of the standard on the original dataset was shifted by a random number between a minimum and a maximum value. The minimum value was the negative of the SFPD, in order to guarantee that the surrogate standard onset occurred after the previous warning signal. The maximum value was the minimum among the latency of the next deviant and the latency of the next warning signal, minus 500 ms. In this way a surrogate standard occurred before the next deviant and the next warning signal, and neither deviants or warning signals appeared in the 500 ms-long window used to decode the SFPD. Since the onset time of standards determines the DMP values used as independent variables in the linear regression model (Section A.5.7), the surrogate datasets only randomized the independent variables of this model. In the surrogate datasets, the distribution of phases at every moment in time was not significantly different from the uniform distribution (p>0.01, Raleigh test). In the main analysis, to decode SFPDs, we only used DMP values at time points at which the distribution of phases was significantly different from the uniform distribution. If we had applied this same criterion with the surrogate epochs we would have no time points to decode SFPDs. Thus, for epochs in surrogate datasets we used the same time points as in the original epochs to decode SFP durations.

For each original dataset, we repeated the above randomization 30 times. For each surrogate dataset, we fitted a decoding model (in exactly the same way as with the original dataset), and computed the correlation coefficient between the model decodings and the SFPDs. We used the median of these 30 correlation coefficients to measure the decoding power of models fitted to surrogate datasets. For models derived from ICs in the left parieto-occipital cluster 04 and the attended visual standards, Figure A.1a plots the decoding power of models fitted to the original dataset versus that of models fitted to surrogate datasets. The decoding power of models fitted to the original dataset was significantly larger than that of models fitted to surrogate datasets. The median pairwise difference between the decoding power of models fitted original datasets minus that of models fitted to surrogate datasets was 0.21, which was significantly larger than zero (95% confidence interval [0.17, 0.25]).

Note that the estimation of the decoding power for models fitted to surrogate datasets was computationally very expensive. For each cluster, standard modality, and attended modality, we fitted approximately 28,500 models, corresponding to 19 subjects × around 50 maximal SFPDs to select the optimal one (Section A.4) × 30 repetitions of the randomization procedure. For this reason, we limited this control to one representative cluster, standard modality, and attended modality.

This control supports the claim that the ITPC evoked by standards is the experimental variable modulated by the SFPD. In particular, it constitutes evidence against the possibility that the observed modulations on ITPC were evoked by the warning signal, since in this case modulations of ITPC by the warning signal should not be affected by the randomization of the onsets of standards.

#### A.1.2 The SFP duration is the modulating variable

It is possible that from the DMP values triggered by standards one could reliably decode any variable, and not only the SFPD. This would be the case, for example, if the decoding model were overfitted to the data. Note, however, that the estimation and the model evaluation methods used here were designed to avoid overfitting (i.e., we used a regularized error function for parameter estimation, Section A.5.8, and cross-validation to evaluate the goodness of fit of models, Section A.5.14). To control for this possibility, we proceed as in Section A.1.1, but with surrogate datasets with shuffled SFPDs. That is, instead of assigning to each trial its corresponding SFPD, we assigned the SFPD of a randomly chosen (without replacement) trial. Since the SFP is the dependent variable in the linear regression model (Section A.5.7), the surrogate dataset only shuffled the dependent values of this model.

Figure A.1b is as Figure A.1a, but for control models estimate from surrogate datasets with shuffled SFPDs. The decoding power of models estimated from original datasets was significantly larger than that of models estimated from the shuffled surrogate datasets. The median pairwise difference between the correlation coefficient for a model estimated from the original dataset minus that for a model estimated from the corresponding surrogate dataset was 0.18, which was significantly different from zero (95% confidence interval: [0.15, 0.24]). Thus, Figure A.1b supports the inference that the SFPD is the experimental variable modulating the ITPC elicited by standards.

### A.2 Correlations between ITC and behavior

For each IC in a cluster and for a given standard modality and attended modality we extracted the ITC value at the peak time and frequency (Section A.5.4). Then we computed the correlation coefficient between peak ITC values of ICs and error rates or mean reaction times of the subjects corresponding to the ICs. Tables A.1 and A.2 show the obtained correlation coefficients and corresponding p-values for error rates and mean reaction times, respectively, across all clusters, standard modalities, and attended modalities. Highlighted in blue are entries with uncorrected p-values smaller than 0.05. A double dagger (dagger) below the image of a cluster in Figure A.2 indicates a significant correlation between peak ITC values and error rates (mean reaction times) in the corresponding cluster and in the standard modality and attended modality given by the color of the double dagger (dagger).

### A.3 Simulations of the SFP effect on ITPC

We simulated the model in Eq. 1 twice, with the precision of the phase noise *n* displaying larger (max*κ* = 5.00, min*κ* = 0.01) and smaller (max*κ* = 3.50, min*κ* = 1.51) oscillations and with parameters *f* = 7 Hz, *θ* = π, *f_n_* = 3 *Hz*, and *s* = 2. For these simulations we used the SFPDs, *SF P D* in Eq. 1, from one experimental session (subject av124a and attended visual standards). The simulated oscillations are shown in Figure A.3 as an erpimage (Makeig et al., 2004) sorted by SFPDs (black curve to the left of time zero). Figure A.3a and A.3b correspond to larger and smaller fluctuations in the precision of the phase noise, respectively. Figure A.4 is as Figure 4 but for the simulated data. The fluctuations in averaged DMP in trials closest to and furthest from the warning signal, those in their difference, and fluctuations in the estimated model coefficients are similar for the simulated and experimental data. The correlation coefficient for the model fitted to data with smaller fluctuations of the precision of the noise was *r* = 0.23 (95% CI=[0.12, 0.34]), which was significantly smaller than that for the model fitted to data with larger fluctuations of the precision of the noise *r* = 0.51 (95% CI=[0.43, 0.59]).

**Figure A.4:**
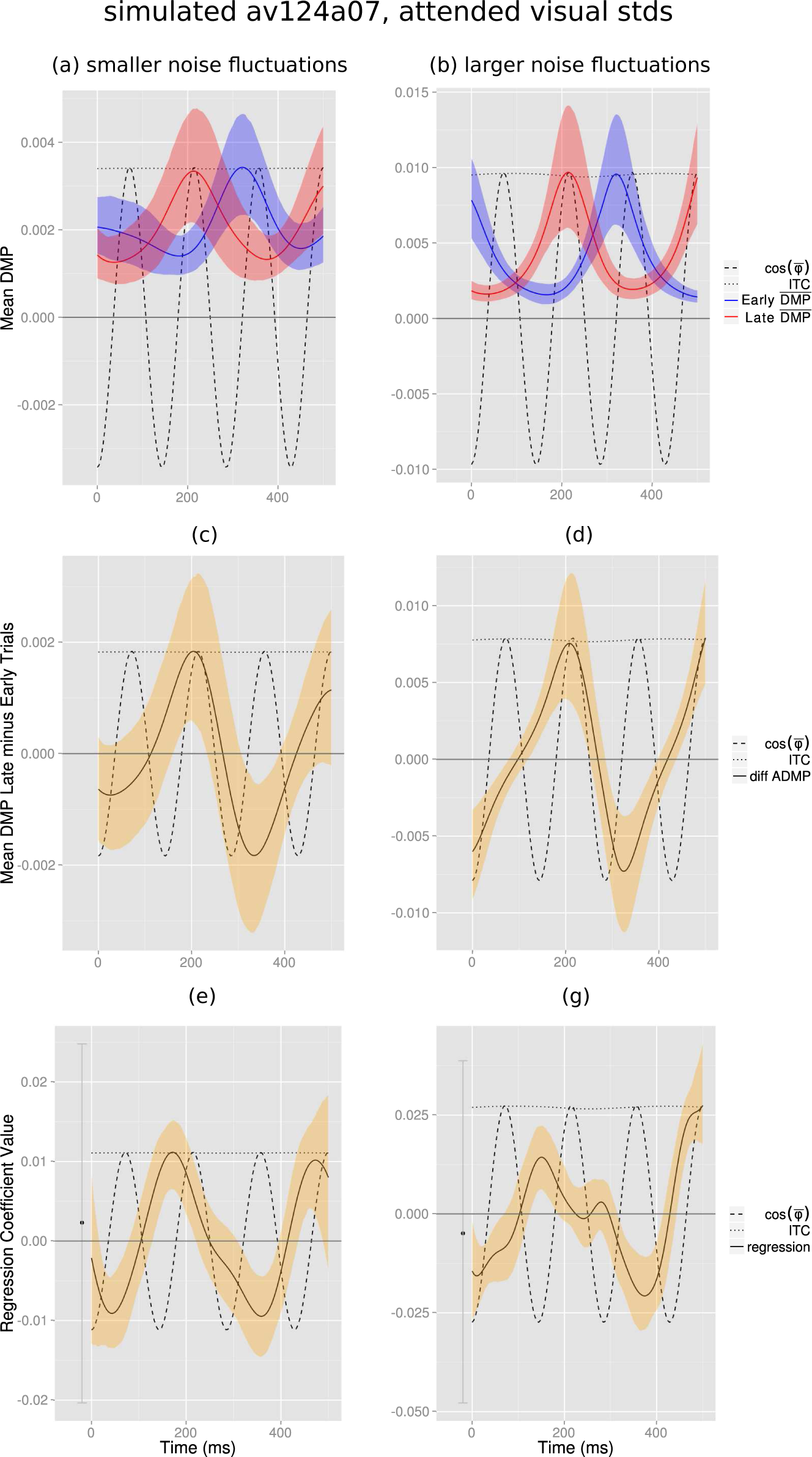
Average DMP in trials furthest from and closest to the warning signal and regression coefficients for data simulated from Eq. 1. Same format as Figure 4. That the panels in this figure capture salient features of those in Figure 4, generated from neural recordings, suggests that the simple oscillatory model in Eq. 1 is a good first description of how the SFPD modulates the phase of neural oscillations.

### A.4 Selection of the optimal maximum SFPD

To fit decoding models we used data from standards that were presented before an optimal maximum SFPD after the warning signal. For models fitted to data from IC 05 of subject av130a, and unattended visual standards, Figure A.5 plots the decoding power of models as a function on their maximum SFPD. We see that the decoding power of models varied smoothly as a function of this maximum SFPD. For each estimated model we selected the maximum SFPD that optimized the decoding power of the model as the optimal maximum SFPD. We chose maximum SFPDs between one second and the largest SFPD of any trial in the data, since maximum SFPDs shorter than one second may be contaminated by modulations from the nearby warning signal. For IC 05 of subject av130a and unattended visual standards, we selected an optimal maximum SFP duration of 2.8 seconds (time highlighted in red in Figure A.5).

**Figure A.5:**
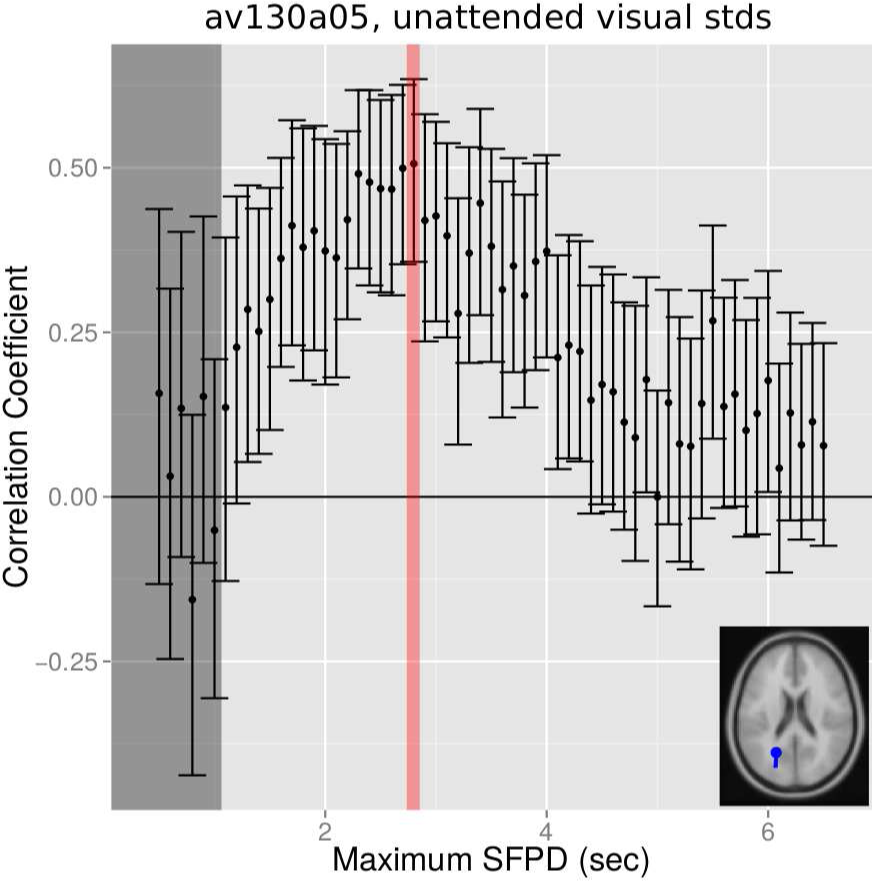
Selection of the optimal maximum SFPD for the decoding model for IC 05 of subject av130a and unattended visual standards. Decoding models were estimated with maximum SFPDs between 500 ms and the largest SFPD in a set of trials, in steps of 100 ms (abscissa), and the correlation coefficient were computed between models’ decodings and experimental SFPDs (ordinate). The SFPD at which this correlation coefficient was largest was selected as the optimal maximum SFPD (red bar). To avoid possible modulations of the ITPC by the warning signal, we excluded from this selection SFPDs shorter than one second (gray bar).

The optimal maximum SFPD gives the range of SFPDs at which modulations of ITPC by the SFPD are stronger. The mean of the optimal maximum SFPD for models corresponding to the visual and auditory attended modality were 2.4 and 2.7 seconds, respectively. A repeated-measures ANOVA with optimal maximum SFPD as dependent variable found a significant main effect of attended modality (F(1,579)=8.45, p=0.0038). A posthoc analysis indicated that the mean of the optimal maximum SFPD was significantly shorter for models estimated from epochs where attention was directed to the visual than to the auditory modality (z=2.91, p=0.0018). Further information on this ANOVA appear in Section A.5.11. Therefore, modulations of ITPC by the SFPD were stronger earlier when attention is directed to the visual than to the auditory modality.

In Section 5.3 we showed that the mean SFPD was significantly shorter in epochs where attention was directed to the visual than to the auditory modality. Coincidentally, in this section we demonstrated that the time window with the strongest modulations of ITPC by the SFPD occurred earlier when attention was directed to the visual than to the auditory modality. This coincidence suggests that subjects had learned the distribution of SFPDs and that their modulations of ITPC reflected this learning.

### A.5 Supplementary methods

#### A.5.1 ICA data preprocessing

For each subject, we performed an ICA decomposition on his/her EEG recorded brain potentials (Makeig et al., 1996). Briefly, if x ∈ ℝ^*N*^ is an *N*-dimensional random vector representing the EEG activity in *N* channels, we estimated a mixing matrix *A* ∈ ℝ^*N*×*N*^ so that

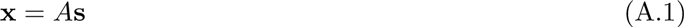

and the components of the random vector s ∈ ℝ^*N*^ were maximally independent. These components are called the independent components (ICs) through the manuscript. We used the AMICA algorithm (Palmer et al., 2007) with one model to compute this decomposition. Prior to the ICA decomposition, the EEG data was high- and low-pass filtered with cutoff frequencies of of 1 and 50 Hz respectively. After this decomposition, non-brain ICs (e.g., components corresponding to eye blinks, lateral eye movements, muscle activity, bad channels, and muscle activity) were removed from the ICA decomposition. We kept, an average, of 26 components per subject, out of the *N* = 32 components obtained in the ICA decomposition.

#### A.5.2 Clustering of ICs

All ICs of all subjects were grouped into clusters according to the proximity of their equivalent dipole locations (see below) using the k-means algorithm. This algorithm groups points of a D-dimensional metric space into clusters, in such a way that the sum of the distances between points and their corresponding cluster centroid is minimized. We represented each IC by the three-dimensional point given by the Talairach coordinates of its equivalent dipole location. A free parameter in k-means is the number of clusters. We set this parameter to 17 to obtain clusters of reasonable coarseness. Clusters 8, 12, and 16 were not analyzed because they contained too few ICs, 4, 4 and 5, respectively (Table A.3).

**Table A.3:**
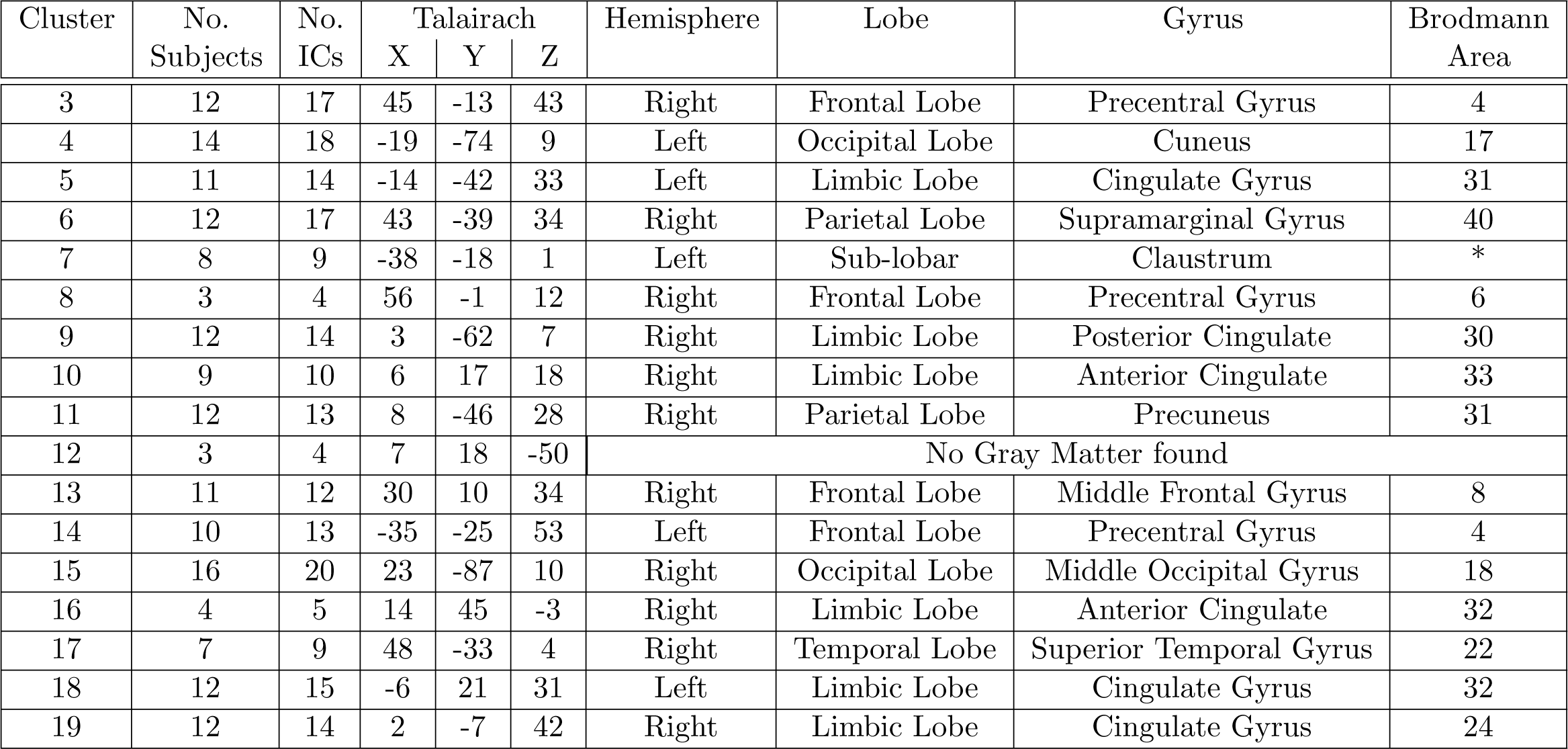
Information about clusters of ICs. The anatomical labels associated with the Talairach coordinates of the clusters’ centroids were extracted using the Talairach client (Lancaster et al., 1997, 2000)

##### Equivalent Dipole location

The scalp map of the ith IC is the ith column of the mixing matrix A in (A.1). For each IC, the location of the electric current dipole whose projection in the scalp best matched the IC scalp map was estimated using the DIPFIT2 plugin of the EEGLAB software. The location of this dipole is the equivalent dipole location of the IC.

#### A.5.3 Circular statistics concepts

This section introduces concepts from circular statistics (Mardia, 1972) used below to define ITC and DMP. Given a set of circular variables (e.g., phases), *θ*_1_, …, *θ*_*N*_, we associate to each circular variable a two-dimensional unit vector. Using notation from complex numbers, the unit vector associated with variable *θ*_*i*_ is:

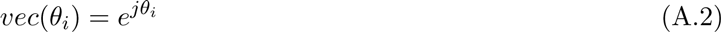

The *resultant vector*, R, is the sum of the associated unit vectors:

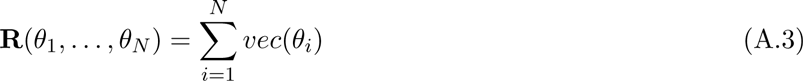

The *mean resultant length*, 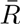, is the length of the resultant vector divided by the number of variables:

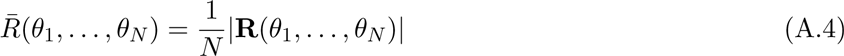

The *circular variance*, *CV*, is one minus the mean resultant length:

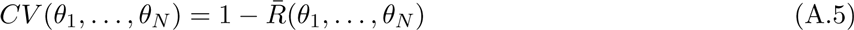

The *mean direction*, 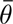, is the angle of the resultant vector:

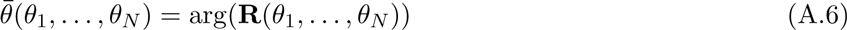

Note that the mean direction (and therefore the DMP) is not defined when the resultant vector is zero, since the angle of the zero vector is undefined.

#### A.5.4 ITC and Peak ITC frequency

The Inter-Trial Coherence (ITC) is a measure of ITPC resulting from averaging phase information among multiple epochs (Tallon Baudry et al., 1996; Delorme and Makeig, 2004). To compute ITC we extracted epochs from one second before to three seconds after the presentation of standards. Then, a continuous wavelet transform was performed on these epochs, using the Morlet wavelet with three significant cycles, eight octaves, and 12 frequencies per octave. With a 250 Hz EEG sampling rate, the previous parameters furnished a Morlet transform with a time resolution of 70.71 ms, and a frequency resolution of 4.5 Hz, both at 10 Hz. The function cwt in the Rwave package of the language R (R Core Team, 2012) was used to compute the continuous wavelet transform. This transform provided the phases of every trial, for frequencies between 0.5 and 128 Hz, and for times between the start and end of the epoch. The ITC of a set of phases *θ*_1_, …, *θ*_n_ (gray vectors in Figure 7) is the mean resultant length (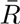, Eq. A.4) of these phases (length of black vector in Figure 7):

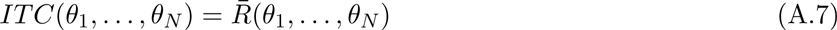

For each IC of every subject, standard modality, and attended modality, we used the corresponding set of epochs to calculate ITC values between the start and end times of the epochs, and between 0.5 Hz to 125 Hz. We selected the peak of these values between 100 and 500 ms after the presentation of the standard at time zero, and between 1 and 14 Hz. The frequency corresponding to this peak (i.e., the peak ITC frequency) was then used to measure the single-trial phase values, as described in Section A.5.5. The median peak ITC frequency and its 95% confidence interval were 4.93 Hz and [4.59, 5.21] Hz, respectively, and the median time of the ITC peak and its 95% confidence interval were 215 ms and [212, 226] ms, respectively. ITC values between -200 and 500 ms around the presentation of attended visual standards, from IC 5 of subject av130a are shown in Figure A.6. The black cross in this figure marks the peak ITC value, occurring at peak frequency 8.15 Hz.

**Figure A.6:**
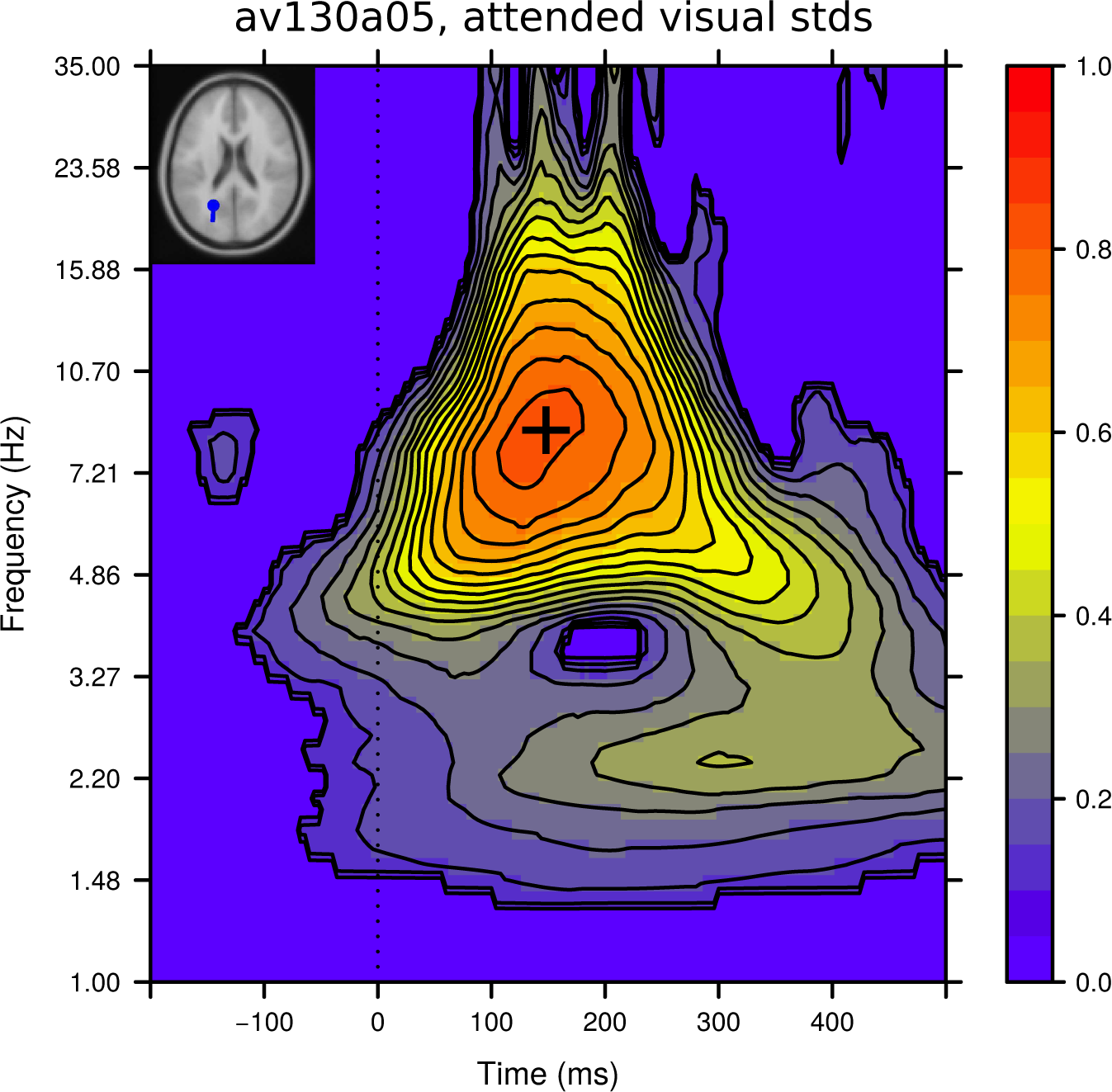
Example set of ITC values and selected peak from IC 05 of subject av130a and attended visual standards. The peak ITC value (black cross) was selected between 100 and 500 ms and between 1 and 14 Hz. The selected peak ITC frequency, time, and amplitude were 8.52 Hz, 148 ms, and 0.83, respectively. Non-significant ITC values (p>0.01, Rayleigh uniformity test) are masked in light blue.

#### A.5.5 Measuring phases in single trials

After selecting the peak ITC frequency (Section A.5.4) for an IC of a subject, a standard modality, and an attended modality, we computed a Gabor transform of all epochs. These epochs started one second before and ended three seconds after the presentation of standards. For this calculation, we used the function cgt in the Rwave package of the language R (R Core Team, 2012). We adjusted the scale of the Gabor’s Gaussian window to obtain three significant sinusoids at the frequency of the Gabor transform. The phases of single trials were extracted from the complex coefficients of this Gabor transform. Figure 2c illustrates calculated phases for IC 13, of subject av124a, and unattended visual standards. Although we computed phases for the whole epoch duration, between -1 and 3 seconds, this figure only shows phases between -100 and 500 ms. To avoid border effects, epochs were clipped to their final size, between 0 and 500 ms, after the computation of this Gabor transform.

#### A.5.6 Relation between the averaged DMP and the ITC

The DMP is a measure of phase decoherence, while the ITC is a measure of phase coherence. Here we derive an upper bound for the averaged DMP, 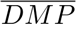, by the ITC (Proposition 1). From this bound we infer that to minimal values of the 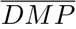 (i.e., 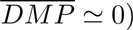) correspond large values of ITC (i.e., *ITC* ≃ 1), and that the maximal value of the 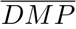 is ½.

##### Proposition 1

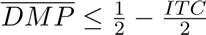

*Proof*. We first rewrite DMP

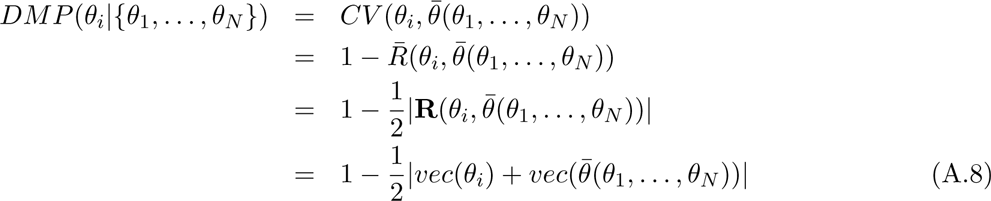

The first equality comes from Eq. 5, the second one from Eq. A.5, the third one from Eq. A.4, and the fourth one from Eq. A.3. We also rewrite 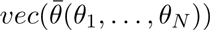:

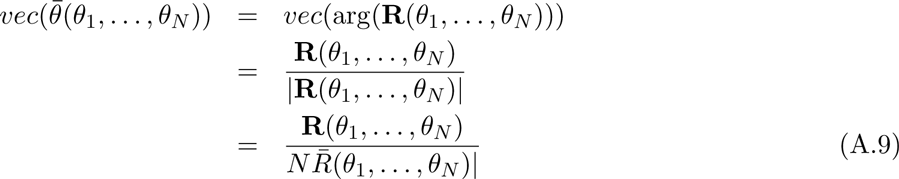

The first equality comes from Eq. A.6, the second one from the fact that *vec* applied to arg of any vector gives the vector normalized to unit length, the third equality comes from Eq. A.4. Then,

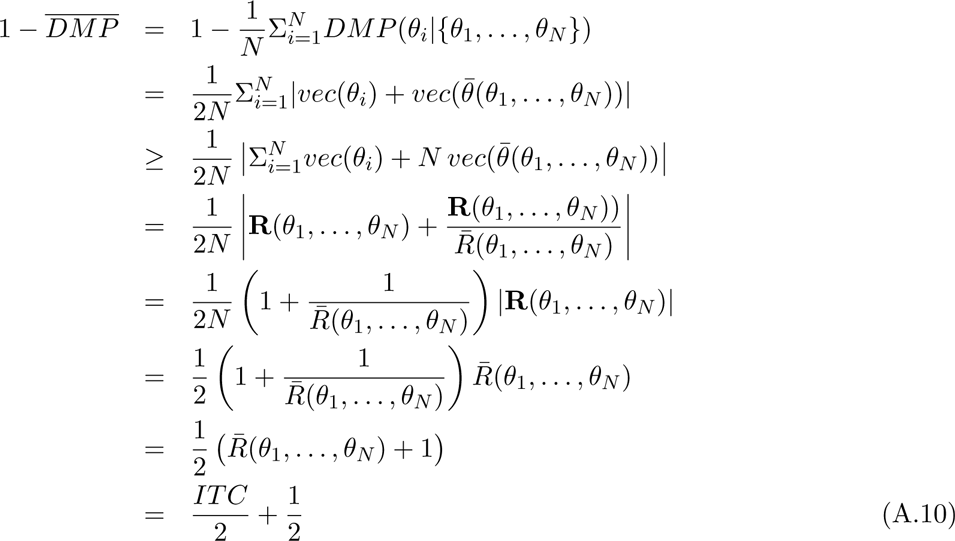

The first equality comes from the definition of 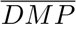, the second one from Eq. 5, the third one from the triangle inequality, the fourth one from Eq. A.3 and Eq. A.9, the fifth one from taking R(*θ*_1_, …, *θ*_N_) as common factor, the sixth one from Eq. A.4, the seventh one from distributing 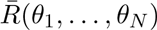, and the eight one from Eq. A.7. The proposition follows by re-arranging Eq. A.10.

Because 0 ≤ *DMP* ≤ 1 it follows that 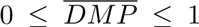. When ITC is maximal (i.e., ITC=1) Proposition 1 shows that 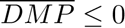, thus 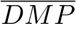 must achieve it minimal value (i.e., 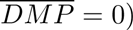 = 0). Also, this proposition shows that ½ is an upper bound for 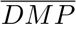, corresponding to zero ITC.

#### A.5.7 Linear regression model

We used a linear regression model to decode the SFPD of trial *n*, *y*[*n*], from the samples of DMP values from this trial, *x*[*n*, ·].

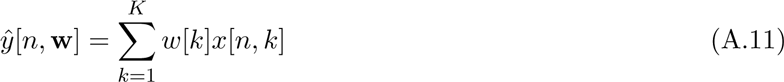

where 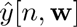 is the decoded SFPD, *w*[*k*] is a regression coefficient, *x*[*n*, *k*] is the DMP value for trial *n* at sample time *k*, and *K* is the number of sample points in the 500 ms-long time window following the presentation of standards for which the distribution of phases across trials was significantly different from the uniform distribution (p<0.01; Rayleigh test).

#### A.5.8 Method to estimate the coefficients in the linear regression model

We seek coefficients w in the linear regression model (Eq. A.11) such that the decodings of the model, 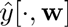, are as close as possible to the experimental SFPDs, *y*[·]. Mathematically, we estimate the regression coefficients w that minimize the least-squares error function

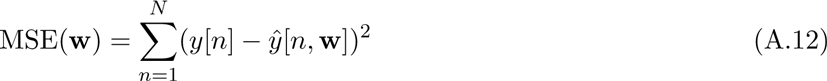

where *N* is the number of epochs. A difficulty in this estimation is that DMP values at neighboring sample points are highly correlated, and correlations increase the variance of ordinary least-squares estimates. In addition, the number of coefficients, *K* in (A.11), is comparable to, and in some cases even larger than, the number of epochs, *N* in (A.12). This further increases the variance of the ordinary least-squares estimates, and when *K* > *N* these estimates are not even defined. To address these problems here we use ridge regression (Hastie et al., 2009, Section 3.4.1), which adds a penalty constraint to the least-squares error function in Eq. A.12, shrinking coefficients estimates toward zero and therefore reducing their variability:

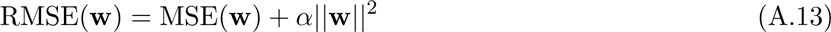

where *α* determines the strength of the constraint (i.e., for larger values of *α* the penalty constraint more strongly bias the coefficient estimates away from minimizing the least-squares error function in Eq. A.12 and toward zero). We call Eq. A.13 the ridge-regression error function.

To find the optimal value w in Eq. A.13 we took a Bayesian approach (Bishop, 2006). We used a Gaussian likelihood function:

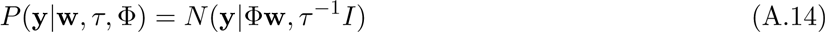

where Φ is the matrix of DMP values (i.e., Φ[*n*, *k*] = *x*[*n*, *k*]) and *τ* is the constant precision of y. We chose a normal-gamma prior for w and *τ*:

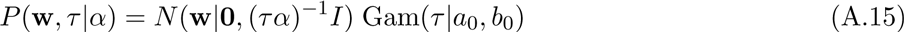

a Gamma hyper-prior for the hyper-parameter *α*:

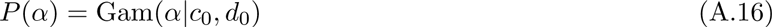

and searched for the values of w that maximized the the log of the posterior distribution:

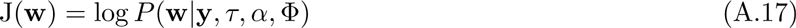

As we prove in Proposition 2, with this choice of likelikhood function and priors, finding the coefficients that maximize the log of the posterior distribution in Eq. A.17 is equivalent to finding the coefficients that minimize the ridge-regression error function in Eq. A.13, as we set to do at the beginning of this section.

Due to the large dimensionality of w, evaluating the posterior distribution in Eq. A.17 is not feasible and one needs to resort to approximation schemes (Bishop, 2006, Chapter 10). These approximations can be stochastic or deterministic. In principle, stochastic approximations, such as Markov Chain Monte Carlo (Metropolis and Ulam, 1949), can give exact evaluations of the posterior distribution, given infinite computational resources, but are often restricted to small-scale problems. Deterministic approximations, are based on approximations of the posterior distribution, and although can never generate exact results, scale well for large problems. Here we use the Variational Bayes deterministic approximation (Bishop, 2006), and implement it as described in Rapela (2016).

We applied a logarithmic transformation to the dependent variable, *y*[*n*], in order to equalize the variance of the residuals (Kutner et al., 2005, Section 3.9), and standardized the dependent variable and regressors to have zero mean and unit variance (Kutner et al., 2005, Section 7.5). Highly influential trials (i.e., trials with a Cook’s distance larger than 4/(N-K-1), where N and K are the number of trials and regressors) were deleted before estimating the model parameters (Kutner et al., 2005, Section 10.4).

##### Proposition 2

Given the likelihood function in Eq. A.14, and the priors in Eq. A.15 and Eq. A.16, the coefficients w maximizing the logarithm of the posterior distribution J(w) in Eq. A.17 minimize the ridge-regression error function RMSE(w) in Eq. A.13.

*Proof*. We first rewrite the joint pdf *P*(y, w, τ, α, Φ) as:

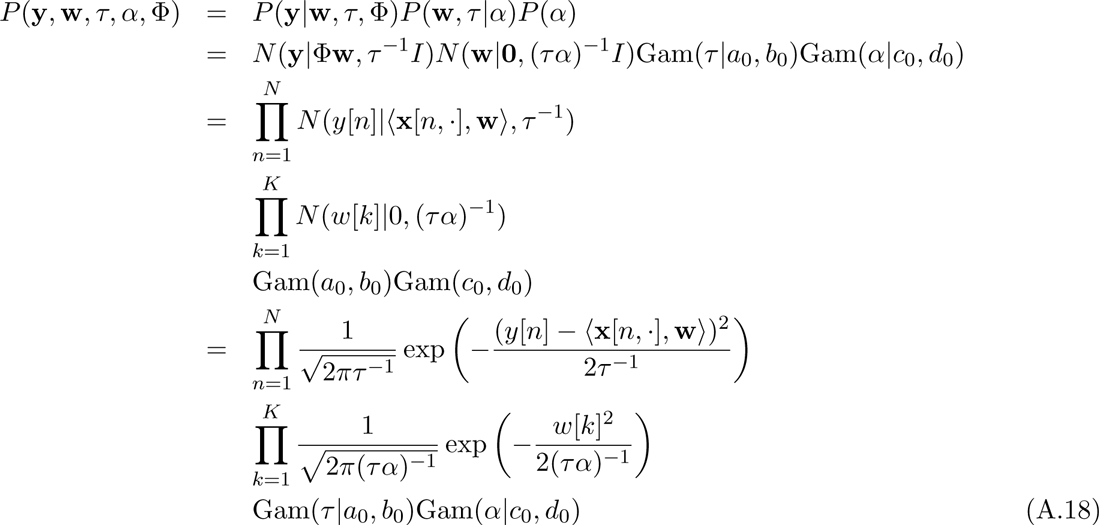

The first equality comes from applying Bayes rule, the second one from substituting Eq. A.14, Eq. A.15, and Eq. A.16 into the first equality, the third one from the fact that the Gaussian distributions in the second equality are independent, and the last one from the definition of a Gaussian distribution.

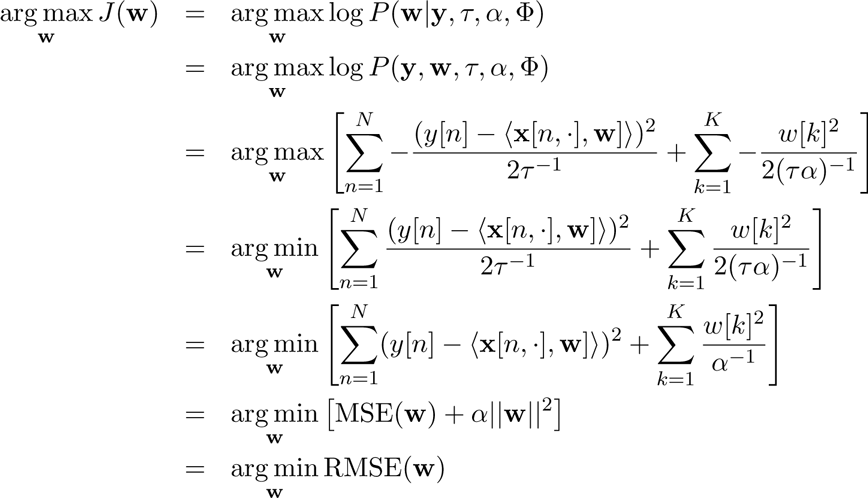

The first equality comes from the definition of *J*(w) in Eq. A.17, the second one from the fact that, by Bayes rule, *P*(w|y, τ, α, Φ) equals *P*(y, w, τ, α, Φ) times a factor independent of w and thus arg max *P*(w|y, τ, α, Φ) = arg max *P* (y, w, τ, α, Φ), the third one by discarding the terms not including w in Eq. A.18 and taking the logarithm, the fourth one from the fact that the maximum of a function is the minimum of the negative of that function, the fifth one from the fact that arg min of a function does not change by multiplying this function by a constant, the sixth one from the definition of MSE(w) in Eq. A.12, and the final one from the definition of RMSE(w) in Eq. A.13.

#### A.5.9 Adjustment for multiple comparisons in correlation tests

For a family of hypothesis *H*_1_ vs. 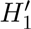, *H*_2_ vs. 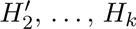 vs. 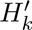, we aim to control the Familywise Error Rate (FWE) defined as FWE=P(Reject at least one *H*_i_|all *H*_i_ are true). For this we define adjusted p-values, 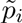, *i* = 1, …, *k*, and we reject *H*_i_ at FWE=α if and only if 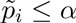. We denote observed values with lowercase and the corresponding random variables with uppercase. The definition of the adjusted p-value, 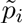, should guarantee that FWE=α. That is, α=FWE=P(Reject at least one *H*_i_ at level α|all *H*_i_ are true)=P(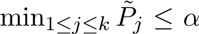|all *H*_i_ are true). We define 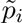 = *P*(min_1≤*j*≤*k*_ *P*_*j*_ ≤ *p*_*i*_|all *H*_i_ are true), where *p*_i_ is the observed non-adjusted p-value. When the p-values *P*_*j*_ are independent it can be shown (Westfall and Young, 1993, p. 47) that the previous definition leads to an exact multiple comparison method (i.e., FWE=α).

In our tests of correlation, the hypothesis 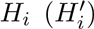 states that the correlation coefficient between two sequences {*a*_1_, …, *a*_*n*_} and {*b*_1_, …, *b*_*n*_} corresponding to the ith cluster, standard modality, and attended modality is zero (different from zero). To compute the adjusted p-value 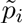 we use a resampling procedure. We first generate nResamples=5,000 samples of min_1≤*j*≤*k*_ *P*_*j*_ under the null hypothesis that all *H*_*i*_ are true and then estimate 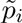 as the proportion of samples smaller or equal than the observed non-adjusted p-value *p*_*i*_. To generate a sample of min_1≤*j*≤*k*_ *P*_*j*_ under the null hypothesis, for each cluster, standard modality, and attended modality we: (1) obtain the sequences {*a*_1_, …, *a*_*n*_} and {*b*_1_, …, *b*_*n*_} to test for significant correlation (e.g., *a*_*i*_=“correlation coefficient between models’ decodings and SFPDs for a subject *i*” and *b*_i_=“error rate for subject *i*” in Figure 5a), (2) shuffle the sequence {*b*_1_, …, *b*_*n*_} (to be under the null hypothesis), (3) compute the p-value of the correlation coefficient between the {*a*_1_, …, *a*_*n*_} and the shuffled {*b*_1_, …, *b*_*n*_}. The sample of min_1≤*j*≤*k*_ *P*_*j*_ under the null hypothesis is then the minimum of all p-values generated in step (3) across all clusters, standard modalities, and attended modalities.

We adjusted for multiple comparison correlation tests between (a) models’ decodings and SFPDs (Figure 2f, colored dots in Figure 3, Table A.4), (b) models’ decoding powers and subjects’ behavioral measures (Figure 5, daggers in Figure 3, Tables A.5 and A.6), and (c) subjects’ peak ITC values and subjects’ behavioral measures (Figure A.2, Tables A.1 and A.2). We begun by defining the family of hypothesis *H*_1_ vs. 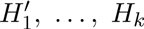 vs. 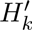 in the definition of the FWE, see above. In (a) we used 19 families of hypothesis, one per subject. The family for subject s contained hypothesis concerning correlation coefficients between model decodings and SFPDs, of all models for subject s across clusters, standard modalities, and attended modalities, that were significantly different from the intercept only model (p<0.01; likelihood-ratio permutation test, Section A.5.14). The null ith hypothesis for subject *s* was 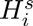={the correlation coefficient between models’ decodings and SFPDs for subject s and the ith combination of cluster, standard modality, and attended modality is zero}. The mean and standard deviation of the number of pairs of hypothesis in a family of hypothesis for a subject was 29.37±12.64. In (b) we used one family of hypothesis for error rates and another one for mean reaction times. Each of these families contained hypothesis regarding the correlation between models’ decoding powers and subjects’ behavioral measures (error rates and mean reaction times) across all clusters, standard modalities, and attended modalities. Each of these correlations was evaluated across all models corresponding to a cluster, standard modality, and attended modality, as in Figure 5. The ith null hypothesis for behavioral measure *b* was 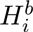={the correlation coefficient between models’ decoding accuracies and subjects’ behavioral measure b for the ith combination of cluster, standard modality, and attended modality equals is zero}. Each family contained 56 pairs of hypothesis (14 clusters × 2 standard modalities × 2 attended modalities). The multiple comparisons procedure in (c) was as that in (b) but using ITC values at the corresponding peak ITC frequencies (Section A.5.4) instead of models’ decoding powers.

**Table A.4:**
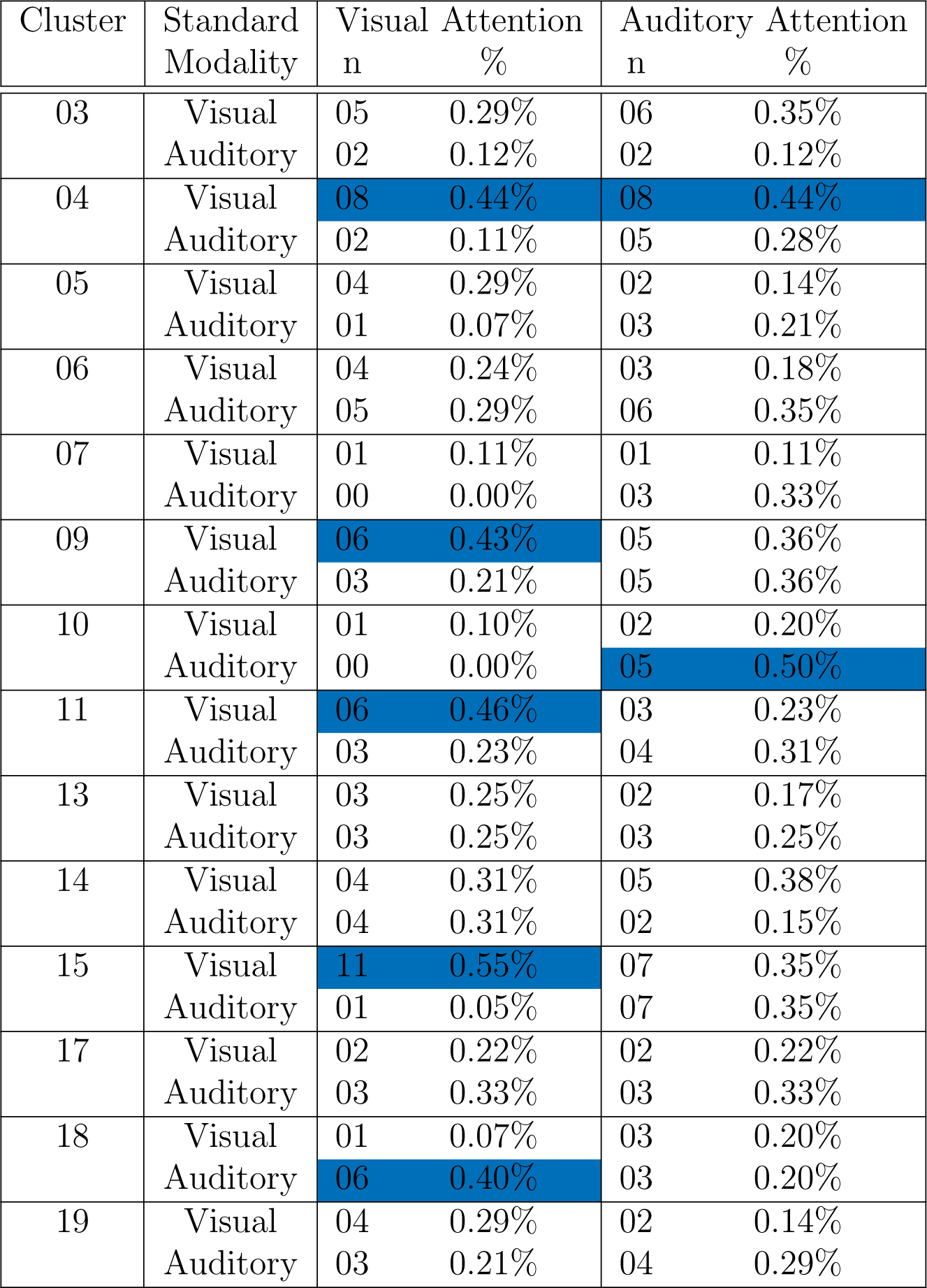
Number (n) and proportion (%) of models with significant correlations (adj p<0.05) between models’ decodings and experimental SFPDs. Cells highlighted in color corresponds to clusters, standard modalities, and attended modalities with more than 40% of significant correlations

**Table A.5:** Correlations between the decoding accuracy of models and subjects’ detection error rates. The decoding accuracy of a model is quantified with the correlation coefficient between decodings of the model and experimental SFPDs. Each cell shows the correlation coefficient, r, and p-values unadjusted, p, and adjusted, adj p, for multiple comparisons. Blue cells highlight correlations with p<0.05 and the red cell corresponds to adj p=0.05.

| Cluster | Standard Modality | Visual Attention |  |  | Auditory Attention |  |  |
| --- | --- | --- | --- | --- | --- | --- | --- |
| | | $r$ | $p$ | $\text{adj}_p$ | $r$ | $p$ | $\text{adj}_p$ |
| 03 | Visual | 0.29 | 0.3054 | 1.00 | 0.30 | 0.2714 | 1.00 |
|  | Auditory | -0.07 | 0.8022 | 1.00 | -0.20 | 0.4766 | 1.00 |
| 04 | Visual | -0.64 | 0.0048 | 0.38 | 0.16 | 0.5212 | 1.00 |
|  | Auditory | 0.06 | 0.8360 | 1.00 | -0.12 | 0.6572 | 1.00 |
| 05 | Visual | -0.50 | 0.0916 | 1.00 | -0.70 | 0.0162 | 0.77 |
|  | Auditory | -0.22 | 0.4846 | 1.00 | -0.10 | 0.7612 | 1.00 |
| 06 | Visual | -0.54 | 0.0336 | 0.95 | -0.33 | 0.2068 | 1.00 |
|  | Auditory | 0.15 | 0.5650 | 1.00 | -0.23 | 0.3694 | 1.00 |
| 07 | Visual | -0.74 | 0.0444 | 0.98 | 0.36 | 0.3490 | 1.00 |
|  | Auditory | -0.17 | 0.6548 | 1.00 | -0.59 | 0.1016 | 1.00 |
| 09 | Visual | -0.20 | 0.4902 | 1.00 | -0.43 | 0.1456 | 1.00 |
|  | Auditory | -0.34 | 0.2412 | 1.00 | -0.25 | 0.3942 | 1.00 |
| 10 | Visual | -0.29 | 0.4274 | 1.00 | 0.49 | 0.1706 | 1.00 |
|  | Auditory | -0.18 | 0.6250 | 1.00 | -0.05 | 0.8794 | 1.00 |
| 11 | Visual | 0.47 | 0.1218 | 1.00 | -0.29 | 0.3104 | 1.00 |
|  | Auditory | -0.43 | 0.1388 | 1.00 | -0.34 | 0.2586 | 1.00 |
| 13 | Visual | -0.19 | 0.5626 | 1.00 | 0.26 | 0.4390 | 1.00 |
|  | Auditory | -0.23 | 0.5264 | 1.00 | 0.11 | 0.7402 | 1.00 |
| 14 | Visual | -0.20 | 0.4964 | 1.00 | 0.34 | 0.2938 | 1.00 |
|  | Auditory | 0.36 | 0.2404 | 1.00 | -0.21 | 0.5538 | 1.00 |
| 15 | Visual | 0.23 | 0.3648 | 1.00 | -0.08 | 0.7376 | 1.00 |
|  | Auditory | -0.38 | 0.1298 | 1.00 | -0.09 | 0.7108 | 1.00 |
| 17 | Visual | 0.06 | 0.8684 | 1.00 | -0.29 | 0.4408 | 1.00 |
|  | Auditory | 0.11 | 0.7900 | 1.00 | -0.08 | 0.8194 | 1.00 |
| 18 | Visual | -0.10 | 0.7612 | 1.00 | -0.11 | 0.7054 | 1.00 |
|  | Auditory | 0.07 | 0.8038 | 1.00 | -0.25 | 0.3922 | 1.00 |
| 19 | Visual | -0.07 | 0.8098 | 1.00 | -0.91 | 0.0004 | 0.05 |
|  | Auditory | -0.62 | 0.0216 | 0.85 | -0.07 | 0.8002 | 1.00 |

**Table A.6:** Correlations between the decoding accuracy of models and subjects’ mean reaction times. Same format as Table A.5.

| Cluster | Standard Modality | Visual Attention |  |  | Auditory Attention |  |  |
| --- | --- | --- | --- | --- | --- | --- | --- |
|  |  | r | p | adj-p | r | p | adj-p |
| 03 | Visual | -0.30 | 0.2574 | 1.00 | 0.02 | 0.9392 | 1.00 |
|  | Auditory | -0.25 | 0.3218 | 1.00 | 0.16 | 0.5410 | 1.00 |
| 04 | Visual | -0.15 | 0.6102 | 1.00 | 0.22 | 0.3654 | 1.00 |
|  | Auditory | -0.04 | 0.8712 | 1.00 | 0.14 | 0.5598 | 1.00 |
| 05 | Visual | 0.13 | 0.6412 | 1.00 | -0.57 | 0.0318 | 0.84 |
|  | Auditory | 0.09 | 0.7596 | 1.00 | -0.24 | 0.4150 | 1.00 |
| 06 | Visual | -0.15 | 0.5712 | 1.00 | -0.14 | 0.5892 | 1.00 |
|  | Auditory | 0.48 | 0.0534 | 0.96 | -0.16 | 0.5414 | 1.00 |
| 07 | Visual | 0.20 | 0.5940 | 1.00 | -0.12 | 0.7456 | 1.00 |
|  | Auditory | 0.11 | 0.7720 | 1.00 | -0.03 | 0.9420 | 1.00 |
| 09 | Visual | -0.01 | 0.9686 | 1.00 | 0.09 | 0.7458 | 1.00 |
|  | Auditory | 0.13 | 0.6486 | 1.00 | 0.11 | 0.6886 | 1.00 |
| 10 | Visual | -0.40 | 0.2706 | 1.00 | 0.54 | 0.1098 | 1.00 |
|  | Auditory | 0.36 | 0.3294 | 1.00 | -0.12 | 0.7308 | 1.00 |
| 11 | Visual | 0.13 | 0.6674 | 1.00 | -0.02 | 0.9430 | 1.00 |
|  | Auditory | -0.45 | 0.1306 | 1.00 | 0.14 | 0.6354 | 1.00 |
| 13 | Visual | 0.12 | 0.6910 | 1.00 | 0.58 | 0.0524 | 0.95 |
|  | Auditory | -0.01 | 0.9682 | 1.00 | 0.35 | 0.2622 | 1.00 |
| 14 | Visual | -0.34 | 0.2598 | 1.00 | -0.03 | 0.9210 | 1.00 |
|  | Auditory | -0.25 | 0.4040 | 1.00 | -0.07 | 0.7996 | 1.00 |
| 15 | Visual | -0.04 | 0.8536 | 1.00 | -0.00 | 0.9890 | 1.00 |
|  | Auditory | -0.07 | 0.7526 | 1.00 | -0.06 | 0.8010 | 1.00 |
| 17 | Visual | -0.23 | 0.5416 | 1.00 | -0.07 | 0.8538 | 1.00 |
|  | Auditory | 0.03 | 0.9172 | 1.00 | -0.10 | 0.7754 | 1.00 |
| 18 | Visual | -0.56 | 0.0270 | 0.80 | 0.14 | 0.6256 | 1.00 |
|  | Auditory | 0.01 | 0.9728 | 1.00 | -0.13 | 0.6502 | 1.00 |
| 19 | Visual | -0.57 | 0.0336 | 0.86 | -0.35 | 0.2044 | 1.00 |
|  | Auditory | -0.16 | 0.5780 | 1.00 | 0.18 | 0.5258 | 1.00 |

#### A.5.10 Calculation of robust correlation coefficients and corresponding p-values

We used skipped measures of correlation (Wilcox, 2012) to characterize the association between pairs of variables in a way that was resistant to the presence of outliers. The calculation of these measures, and the estimation of their bootstrap confidence intervals, followed the procedures described in (Wilcox, 2012; Pernet et al., 2013).

Skipped correlations are obtained by checking for the presence of outliers, removing them, and applying some correlation coefficient to the remaining data (Wilcox, 2012, Chapter 9). We identified and removed bivariate outliers. Bivariate outliers were found using a projection method. The underlying idea behind projection methods is that, if a mutlivariate data point is an outlier, then it should be an outliers along some one-dimensional projection of all data points. For each data point X_*i*_, the projection method used in this manuscript orthogonally projected all data points onto the line joining a robust estimate of the center of the data cloud (given by the medians of the correlated variables) and X_*i*_, and then found the outliers of this one-dimensional projection using the boxplot rule (which is based on interquartiles distances, Frigge et al., 1989). A data point was declared outlier if it was an outlier in any one-dimensional projection. Bivariate outliers were selected using the function outpro freely available from the web site of Dr. Rand Wilcox at http://dornsife.usc.edu/labs/rwilcox/software/.

To estimate the association between single-trial decoding errors and SFPDs (e.g., Figure 2f), we used the skipped Pearson correlation coefficient (i.e., after outlier removal, the remaining data points were correlated using the Pearson product moment correlation coefficient; function pcor from Dr. Wilcox’s website). The use of this correlation coefficient requires that the marginals of the data are approximately normal, as it was the case in the previous data.

The averaged behavioral data tended to be bimodally distributed, with a group of subjects displaying better performance and another group showing worse performance. Thus, the averaged behavioral data was not approximately normal, and it was not possible to use the skipped Pearson correlation coefficient to asses its association with models’ decodings. Instead, we quantified this association using the skipped Spearman correlation coefficient (i.e., after outlier removal, the remaining data points were correlated using the Spearman rank correlation coefficient; function spear from Dr. Wilcox’s website). The use of this correlation coefficient does not assume that the distribution of the correlated variables is bivariate normal, but requires that their association be monotonic, as it was the case for our averaged data.

To compute p-values corresponding to hypothesis tests of robust correlation coefficients we used permutation tests following the procedure given in Pernet et al. (2013). Given a dataset of two sequences to correlate, we (1) removed bivariate outliers from this dataset, (2) computed the correlation coefficient between the sequences in the outliers-removed dataset (using functions pcor and spear from Dr. Wilcox’s website for Pearson and Spearman correlation coefficients, respectively), (3) computed nResamples=5000 correlation coefficients between the first sequence and a random permutation of the second sequence in the outliers-removed dataset (again using functions pcor or spear), and (4) we calculated a two-sided p-value as the proportion of correlation coefficients computed in step (3) whose absolute value was larger than that of the correlation coefficient computed in step (2).

#### A.5.11 ANOVA procedure

The ANOVA procedures described below started with a model with a large number of independent factors. We selected a subset of these factors using a stepwise procedure based on the Akaike Information Criterion, as implemented in the function stepAIC of the package MASS of R (R Core Team, 2012). We then tested for significant differences in the mean of the dependent variable across the selected subset of independent factors. We report the F-value and p-value of all significant effects. For the significant main effects we also report the p-value of posthoc tests for differences across the levels of the corresponding factors, and the sign of these differences.

##### Attentional effects in ANOVAs

Attentional effects in the SFP effect on ITPC could appear in ANOVAs as significant differences between models estimated from attended and unattended stimuli of the same or different standard modalities (e.g., a significant difference between models estimated from epochs aligned to the presentation time of visual standards with attention directed to the visual or auditory modality, or a significant difference between models estimated with attention directed to the visual modality from epochs aligned to the presentation of visual or auditory standards). Here we only study attentional effects in ANOVAs by comparing models estimated from attended and unattended stimuli of the same standard modality. We do so because differences in models estimated from stimuli of different standard modalities could be due to differences in brain responses to stimuli of distinct modalities and unrelated to attention.

As mentioned in Section 5.3, for any standard modality, the mean number of epochs aligned to the presentation of attended stimuli was larger than the mean number of epochs aligned to the presentation of unattended stimuli. This difference invalidates the comparison between models estimated from attended and unattended stimuli of the same standard modality, since any discrepancy between these models could simply be due to differences in the amount of data used to estimate them. Thus, attentional effects in the SFP effect on ITPC were investigated using ANOVAs on datasets where the number of epochs aligned to attended and unattended stimuli had been equalized, as described in Section 5.3.

##### Non-repeated-measures ANOVA

We performed non-repeated-measures ANOVAs using the function aov of the package stats of R (R Core Team, 2012). We removed from the analysis those trials for which the absolute value of the standardized residuals was greater than the 95% quantile of the normal distribution. When a significant main effect was observed in the ANOVA we created a set of confidence intervals on the differences between the means of the levels of the corresponding factor with 95% family-wise probability of coverage using the Tukey’s ‘Honest Significance Difference’ method, as implemented in the function TukeyHSD of the package stats of R (R Core Team, 2012). For the pairs of levels with a significant difference in the mean of the dependent variable we reported the p-value, corrected for multiple comparison, of the test of the hypothesis that the previous difference is distinct from zero. We also reported the sign of this difference extracted from the sign of the corresponding confidence interval.

##### Repeated-measures ANOVA

We performed repeated-measures ANOVAs by estimating mixed-effects models [Pinheiro and Bates, 2000] with the function lme of package nlme of R (R Core Team, 2012). Trials for which the absolute value of the normalized residuals (i.e., standardized residuals pre-multiplied by the inverse square-root factor of the estimated error correlation matrix) was greater than the 95% quantile of the normal distribution were removed from the analysis. When a significant main effect was observed in the ANOVA, we performed posthoc hypothesis tests using the function glht from the package multcomp of R (R Core Team, 2012). This function controls for multiple comparisons by using multivariate t, or Gaussian, distributions to compute p-values associated with multiple hypothesis (Bretz et al., 2010).

#### A.5.12 Further information on ANOVAs

Maximum SFPD (Section A.4): We performed an initial repeated-measures ANOVA (see *Repeated-measures ANOVA* in Section A.5.11) using data from all models significantly different from the intercept-only model (p<0.01, likelihood-ratio permutation test, Section A.5.14) with the natural logarithm of the maximum SFPD as dependent variable (we selected this transformation of the dependent variable to normalize the distribution of the ANOVA residuals). The input to the stepwise selection procedure was a model with standard modality, attended modality, their interaction, and cluster of ICs as independent variables. This procedure selected a model with standard modality, attended modality, and their interaction as independent variables. An ANOVA on the selected model showed significant main effects for attended modality (F(1,615)=15.15, p=0.0001) and for the interaction between standard modality and attended modality (F(1,515)=0.0122, p=0.0122). Since this interaction indicates an attentional effect, we repeated the previous analysis in a dataset with equalized number of epochs aligned to the presentation of attended an unattended standards (see *Attentional effect in ANOVAs* in Section A.5.11). With the equalized dataset, the model selection procedure selected a model with only attended modality as independent factor, and an ANOVA on this model showed a significant main effect of this factor (F(1,579)=8.45, p=0.0038). A posthoc analysis indicated that the maximum SFPD was shorter for the visual than the auditory attended modality (z=2.91, p=0.0018).

**Absolute value of the correlation coefficient between models’ decoding accuracies and subjects averaged behavioral measures** (Section 2.4): We performed a non-repeated-measures ANOVA (see *Non-repeated-measures ANOVA* in Section A.5.11) using data from all models significantly different from the intercept-only model (p<0.01, likelihood-ratio permutation test, Section A.5.14) with the absolute value of the correlation coefficient between models’ decoding accuracies and subjects mean reaction times or error rates as dependent variable. The input to the stepwise selection procedure was a model with standard modality, attended modality, their interaction, cluster of ICs, and type of behavioral measure (i.e., error rate or mean reaction times) as independent variables. This procedure selected a model with standard modality and type of behavioral measures as independent variables. An ANOVA on the selected model showed significant main effects of attended modality (F(1,109)=8.22, p=0.005) and of the type of behavioral measure (F(1,109)=7.26, p=0.0082). A posthoc analysis showed that the mean of the absolute value of the correlation coefficient was larger for the visual than the auditory standard modality (p=0.005; Tukey test) and larger for error rates than mean reaction times (p=0.0082; Tukey test).

**SFPD** (Section 5.3): We performed a repeated-measures ANOVA using data from all epochs with the cubic root of the SFPD as dependent variable (we selected this transformation of the dependent variable to normalize the distribution of the ANOVA residuals). The input to the stepwise selection procedure was a model with standard modality, attended modality, and their interaction as independent variables. This procedure selected a model with all independent variables. An ANOVA on the selected model showed a significant main effect of attended modality (F(1,16264)=234.31, p<0.0001). A posthoc analysis showed that the mean SFPD was shorter in epochs where attention was directed to the visual than to the auditory modality (z=15.31; p<2e-16).

**Time of Peak Coefficient** (Section 2.5): We performed a first repeated-measures ANOVA using data from all models significantly different from the intercept-only model (p<0.01, likelihood-ratio permutation test, Section A.5.14) with the time of the largest peak as dependent variable. The input to the stepwise selection procedure was a model with standard modality, attended modality, group of clusters (as shown in Figure 3), and their triple interaction as independent variables. This procedure selected all independent variables and their interactions, but in a subsequent ANOVA only the effect due to the interaction between standard modality and groups of clusters (F(4, 230)=2.9026, p=0.0227) and the main effect of attended modality (F(1, 230)=4.2590, p=0.0402) remained significant. A posthoc analysis on attended modality showed that the peak occurred earlier for auditory than visual attention (z=1.842, p=0.0327, left black asterisk in Figure 6a). To disentangle the interaction between standard modality and group of clusters we performed a second and a third repeated-measures ANOVA using only models corresponding to visual and auditory standards, respectively.

The input to the stepwise procedure for the second repeated-measures ANOVA for models corresponding to visual standards was a model with peak time as dependent variable and attended modality, group of clusters, and their interaction as independent variables. This procedure selected all independent variables and interactions, but a subsequent ANOVA showed that only the main effects of attended modality (F(1,135)=11.1939, p=0.0011) and groups of clusters (F(4,135)=4.8073, p=0.0022) were significant. A posthoc analysis revealed that the peak of coefficients occurred earlier when attention was oriented to the visual than auditory modality (z=3.512, p=0.000455) and in central than occipital brain regions (z=4.163, p=3.14e-5, red asterisks in Figure 6b).

The input to the stepwise procedure for the third repeated-measures ANOVA for models corresponding to auditory standards was the same as that for models corresponding to the visual modality. This procedure found no significant variable.

From Figure 6 it is apparent that the strongest modulations of the timing of the SFP effect on ITPC occur in the central group of clusters. To test this observation statistically, we performed a fourth repeated-measures ANOVA restricted to models corresponding to the central group of clusters. The input to the stepwise procedure was a model with peak time as dependent variable and standard modality, attended modality, and their interaction as independent variables. This procedure selected the two independent variables and their interaction. However, the significance of the interaction term disappeared when we repeated this analysis with a dataset with equal number of epochs aligned to the presentation of attended and unattended standards (see *Attentional effect in ANOVAs* in Section A.5.11), so we did not used the interaction term in the following. A repeated-measures ANOVA found significant main effects for standard modality (F(1, 43)=14.2173, p=0.0005) and for attended modality (F(1, 43)=4.5061, p=0.0396). A posthoc analysis showed that the peak coefficient occurred earlier in models corresponding to auditory standards (z=3.406, p=0.000659; black asterisks in Figure 6b), and in models estimated with attention directed to the auditory modality (z=2.178, p=0.029360; black asterisk next to Central cluster group in Figure 6a).

#### A.5.13 Identification of peaks in coefficients

We identified positive and negative peaks in the coefficients of decoding models significantly different from the intercept-only model (p<0.01, likelihood-ratio permutation test, Section A.5.14). We only searched for peaks at coefficients significantly different from zero (95% bootstrap CI for regression coefficients, Section A.5.14), and occurring after 200 ms (to avoid possible effects from the warning signal). We filtered continuous subsets of the coefficients with a finite impulse response filter of type I, order two, and cutoff frequency of three times the peak ITC frequency. We selected in the filtered coefficients the local maxima larger than zero (local minima smaller than zero) as the positive (negative) peaks. From all positive and negative peaks, and from all continuous subsets of the coefficients, we selected the peak with maximum absolute value.

#### A.5.14 Additional statistical information

Most statistical figures presented in this paper are based on the bootstrap method (Efron and Tibshirani, 1993).

**Crossvalidated decodings** All models decodings were cross validated using the leave-one-out method with the function `crossval` of the package `bootstrap` of `R` (R Core Team, 2012).

**95% bootstrap CIs for regression coefficients** (Figure 2e) We performed 2,000 ordinary bootstrap resamples of the trials (with the function `boot` of the package `boot` of `R` (R Core Team, 2012)), and for each resample we estimated a set of regression coefficients, as described in Section A.5.7. Having estimated 2,000 sets of regression coefficients, we computed 95% percentile confidence intervals with the function `boot.ci` of package `boot` of `R` (R Core Team, 2012).

**95% bootstrap CI for difference in paired means** of correlation coefficients obtained from the original versus a surrogate dataset (Figure A.1). For each pair of correlation coefficients (obtained by correlating SFPDs and their decodings in the original and surrogate datasets) we subtracted the correlation coefficient from the original minus the surrogate dataset. Then we performed 2,000 ordinary resamples (with the function `boot` of the package `boot` of `R` (R Core Team, 2012)) and computed a 95% percentile bootstrap confidence interval with the function `boot.ci` of package `boot` of `R` (R Core Team, 2012).

**95% bootstrap CI for difference in averaged DMP** between trials with the longest and shortest SFPD (Figures 4a-c). We calculated 2,000 stratified bootstrap resamples of the 20% of trials with the shortest and longest SFPDs, using the function `boot` of the package `boot` of `R` (R Core Team, 2012) with the `strata` option). For each resample, and for each sample point, we subtracted from the mean DMP across the 20% of trials with longest SFPD the mean DMP across the 20% of trials with shortest SFP. Having estimated 2,000 bootstrap differences in average DMP values between trials with the longest and shortest SFPD, we computed a 95% percentile bootstrap confidence interval for these differences with the function `boot.ci` of package `boot` of `R` (R Core Team, 2012).

**Likelihood-ratio permutation test for linear model We** used it to test the alternative hypothesis, H1, that there is association between the dependent and independent variables of a linear regression model against the alternative one, H0, that there is no association. The statistic for this test is the logarithm of the ratio between the likelihood of the data given a full model and that given a null model. A full model associates all independent variables with the dependent one, as in (A.11), while a null model associates only a constant term with the dependent variable. The likelihood of the data given a model is derived from (A.14). The test was conducted as follows. We first measured the value of the test statistic in the original dataset, *T*_*obs*_. Then, we built the distribution of the test statistic under H0, by generating R=2000 datasets where there was no association between the dependent and independent variables (by permuting the dependent value assigned to independent values) and measuring the test statistic in these datasets. The one-sided p-value of the test is the proportion of samples of the test statistic under H0 that are larger than *T*_*obs*_.

**Normalized cross correlation** (Section 2.3) We normalized the cross correlation (at lag zero) between two time series in such a way that the normalized correlation of a time series with itself is one. We defined:

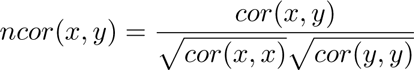

where *x* and *y* are two time series, and *ncor*(*x*, *y*) and *cor*(*x*, *y*) are the normalized and unnormalized correlations, respectively, between these time series.

### A.6 Supplementary figures

Figure A.7 shows average DMPs for the 20% trials with the shortest and longest SFPDs in ICs from the left parieto-occipital cluster 04.. Figure A.8 plots examples of deviant foreperiod effects on reaction times and detectability. Figure A.9 plots axial, coronal, and sagital views of all clusters. Figure A.10 plots the distribution of deviant latencies and SFPDs.

**Figure A.7:**
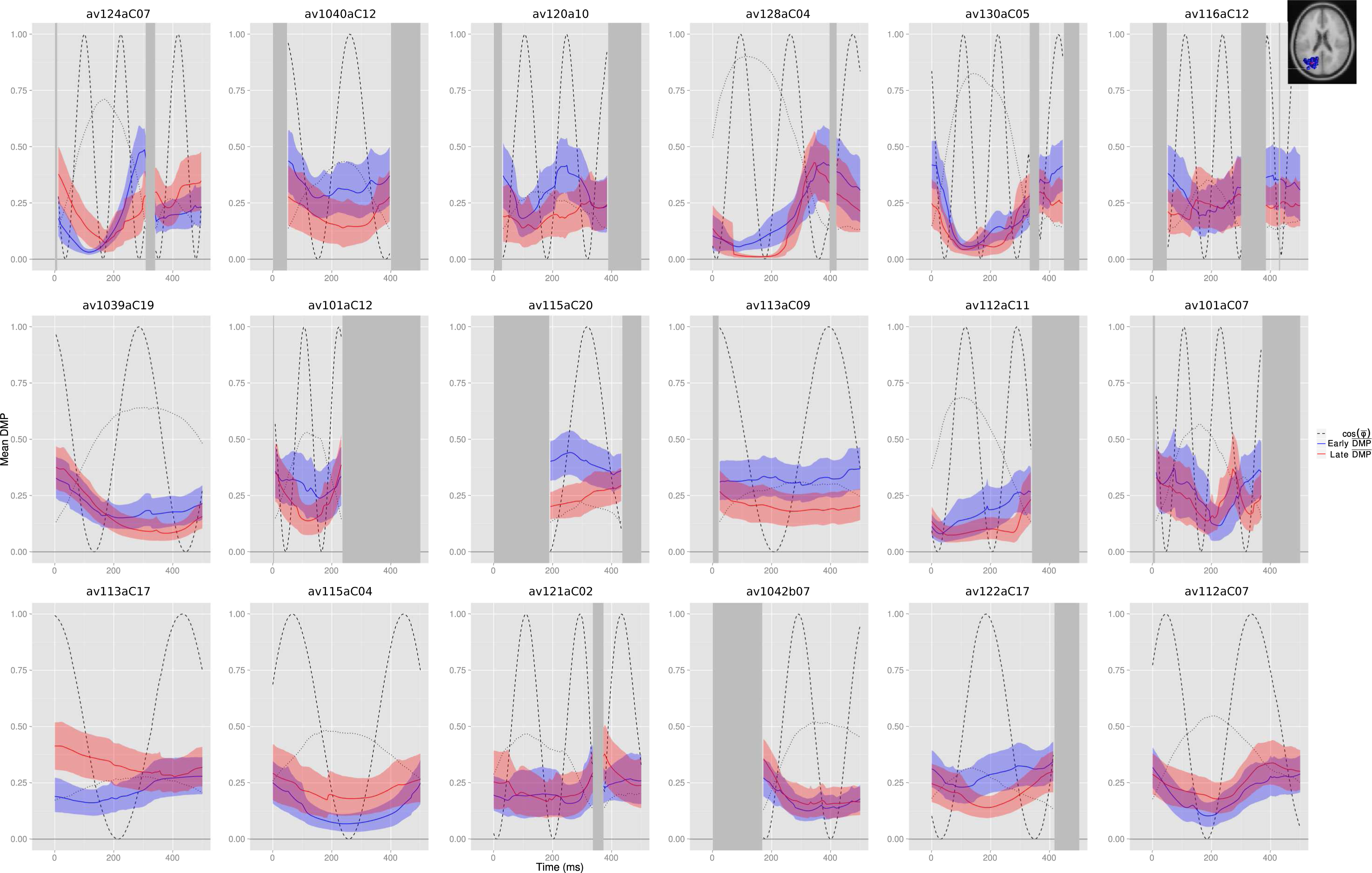
Average DMPs for the 20% trials with the shortest (blue traces) and longest (red traces) SFPDs computed from trials from the left-parieto-occipital cluster 04 (top-right inset) and attended visual stimuli. Panels are sorted from left to right and top to bottom by decreasing decoding power of the corresponding model. These averages suggest that the averaged DMP oscillates at a frequency around 1 Hz and that the phase of these oscillations at time zero is different between trials closest and furthest away from the warning signal.

**Figure A.8:**
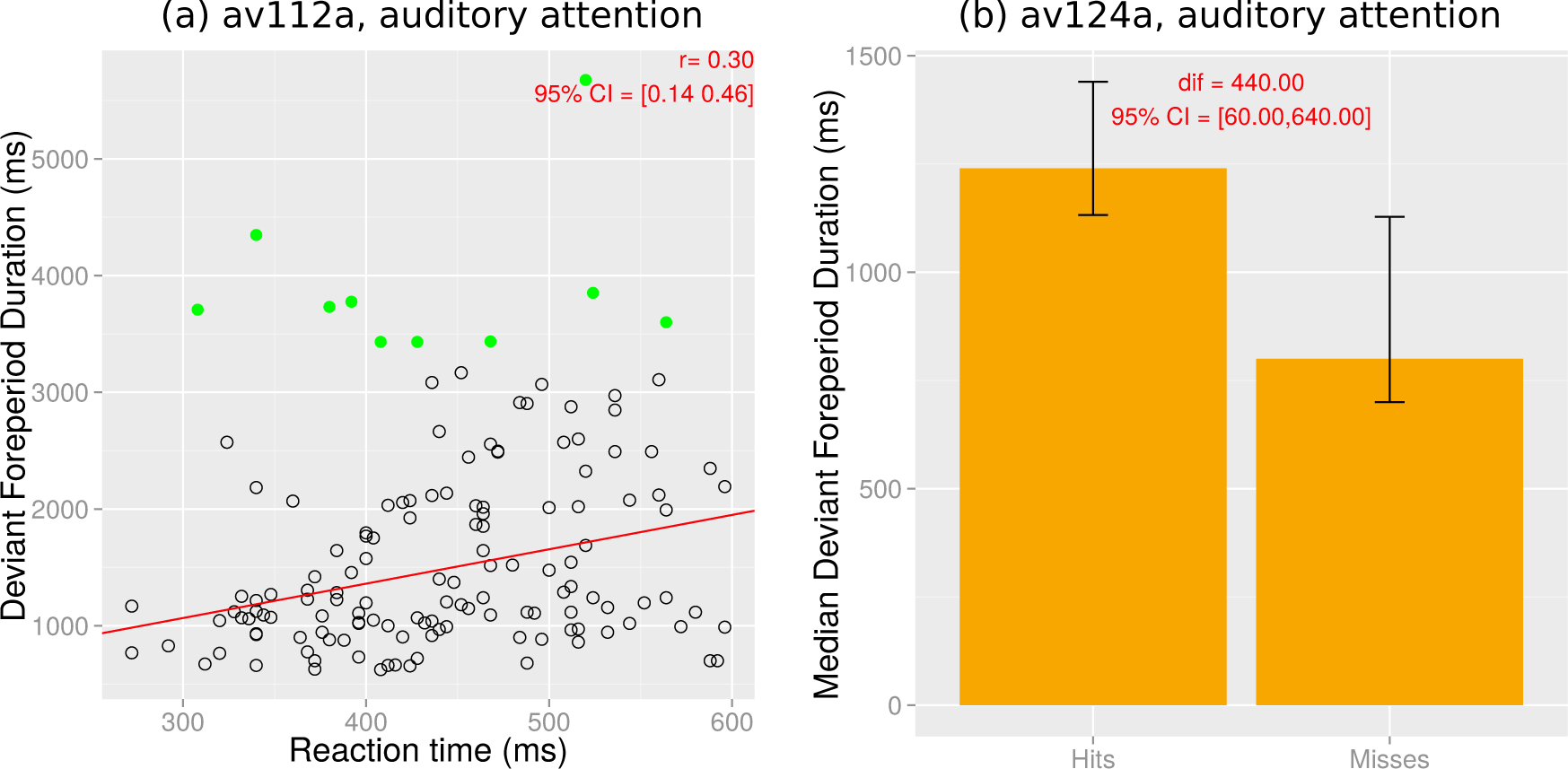
Examples of significant deviant foreperiod effects on reaction times (a) and detectability (b). (a) Deviant foreperiod durations as a function of subject reaction times, for subject av112a and the auditory attended modality. Points colored in green indicate outliers detected in the computation of the robust correlation coefficient (Section A.5.10). (b) Median deviant foreperiod duration for hits and misses, for subject av124a and the auditory attended modality.

**Figure A.9:**
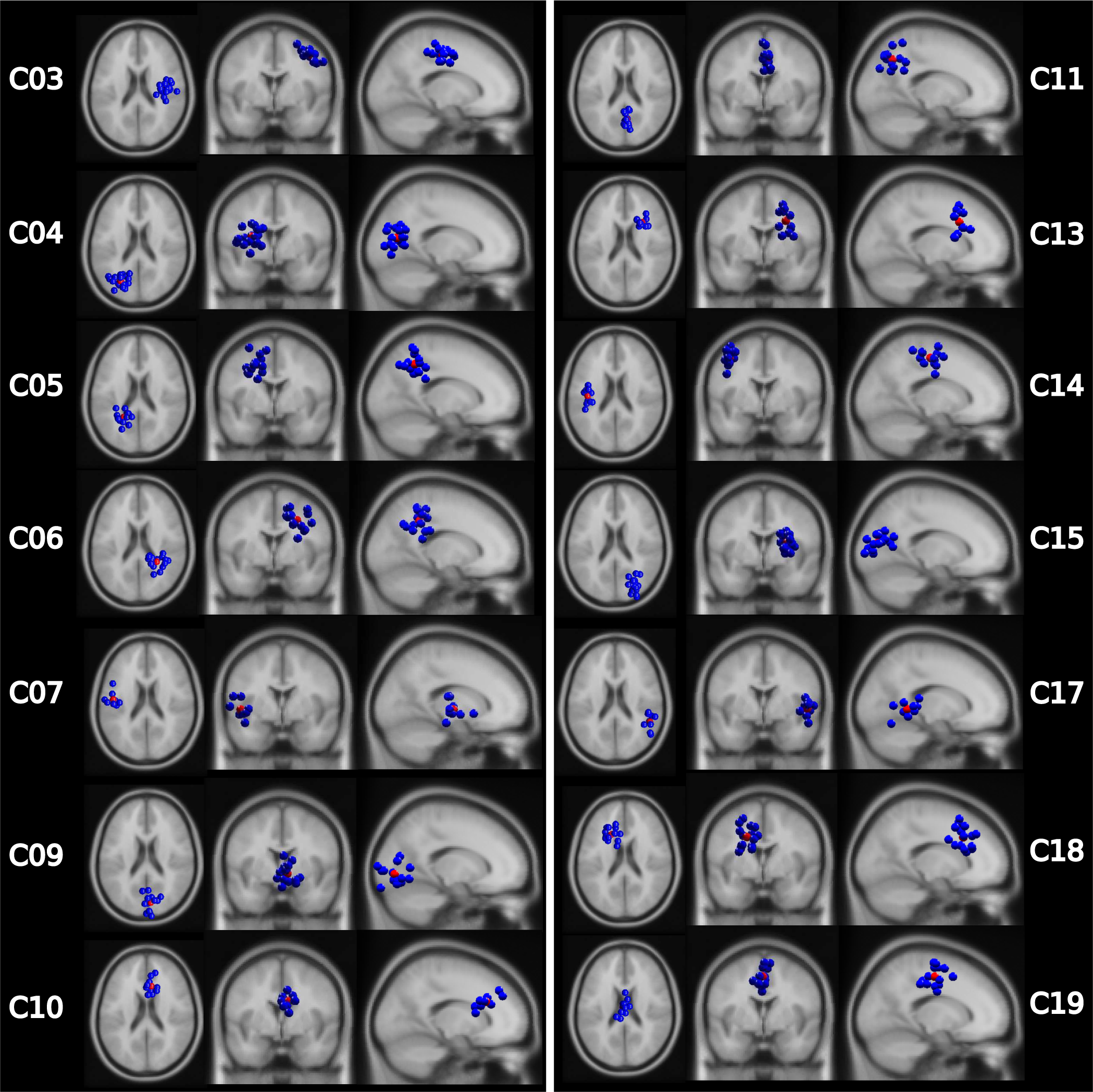
Clusters if ICs. Left, center, and right columns correspond to axial, coronal, and sagital views, respectively.

**Figure A.10:**
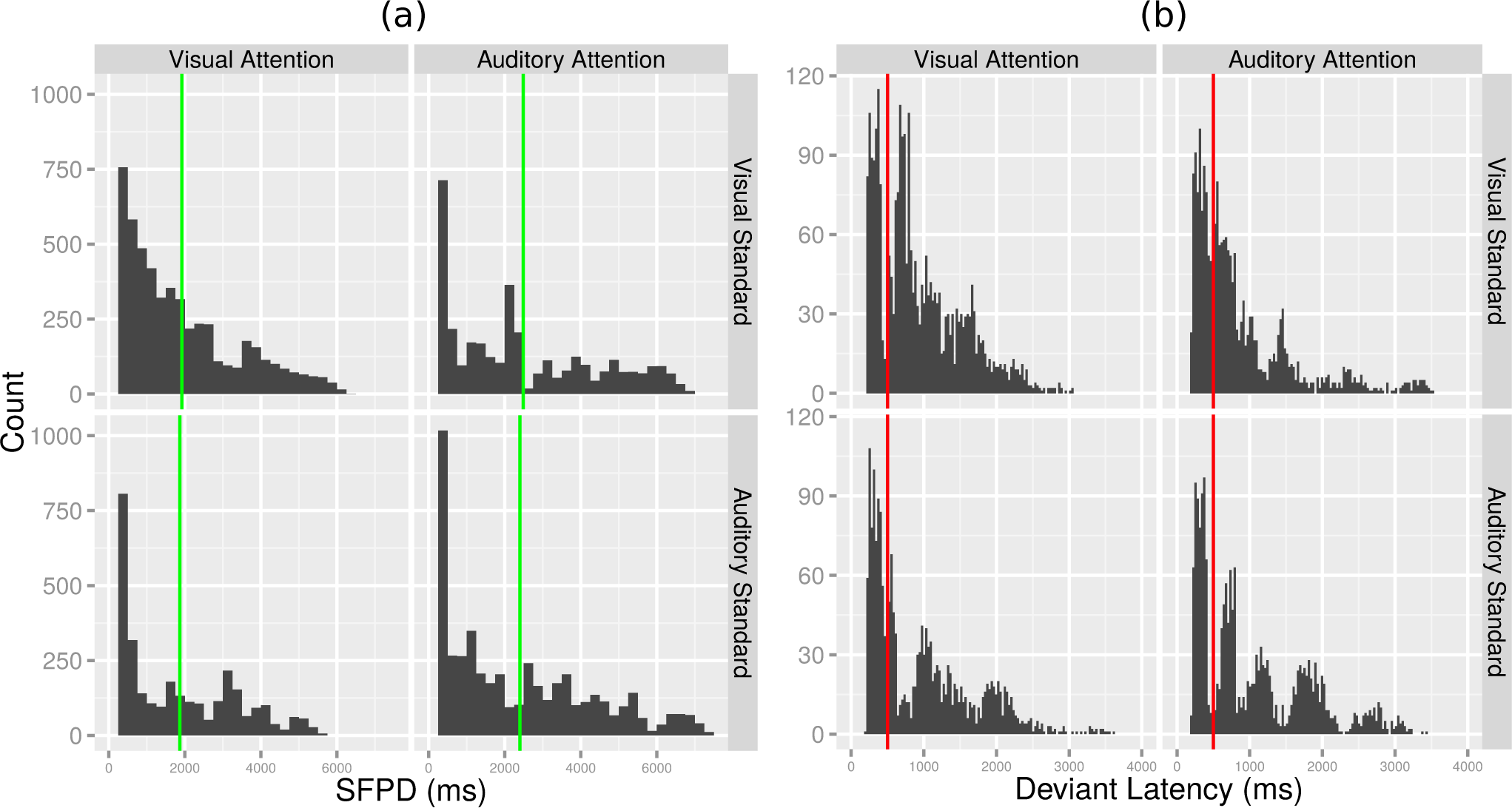
Distributions of SFPDs (a) and of latencies of deviants (b) across standard modalities (rows) and attended modalities (columns). (a) The mean SFPD in epochs corresponding to visual attention (1.9 sec, left panels in (a)) is significantly shorter than that in epochs corresponding to auditory attention (2.4 sec, right panels in (a)), as for the optimal maximum SFPDs of models (see Section A.4). (b) To avoid possible response-related movements artifacts in the EEG, epochs including deviants with latencies shorter than 500 ms were excluded from the analysis. The number of such deviants is given by the sum of counts to the left of the red vertical lines in (b).

### A.7 Supplementary tables

Table A.3 provides the Talairach coordinates and anatomical labels of the centroids of the clusters in Figure 3. For each cluster, standard modality, and attended modality, Table A.4 gives the number of models with decodings significant correlated with SFPDs.

### A.8 Code and sample data

To facilitate the application of the single-trial decoding methods described in this manuscript to other EEG studies, we provide the R (R Core Team, 2012) code implementing these methods, the EEG data from the left parieto-occipital cluster 04, and instruction on how to use this code to analyze this data at http://sccn.ucsd.edu/∼rapela/avshift/codeAndSampleData/

